# Expression of Angiotensin converting enzyme 2 (ACE2) in the dorsolateral striatum is critical for the temporal coordination of acetylcholine and dopamine, and motor learning

**DOI:** 10.1101/2025.10.25.684469

**Authors:** Aria R. Walls, Dustin R. Zuelke, Andreas H. Kottmann

**Affiliations:** Department of Molecular, Cellular and Biomedical Sciences, CUNY School of Medicine, City University of New York, New York, NY, USA; City University of New York Graduate Center, Molecular, Cellular and Developmental Subprogram, New York, NY, USA; City University of New York Graduate Center, Neuroscience Collaborative, New York, NY, USA

**Keywords:** Angiotensin Converting Enzyme 2, Cholinergic Interneurons, Temporal Coordination of Dopamine and Acetylcholine, Motor Learning, Parkinson Disease

## Abstract

Neuropeptides are key modulators of adult neurocircuits, balancing their sensitivity to both excitation and inhibition, and fine-tuning fast neurotransmitter action under physiological conditions. We found that the mono-carboxypeptidase angiotensin-converting enzyme 2 (ACE2), which can take part in producing and/or processing of several neuroactive peptides in the mammalian brain and is best known for converting the pro-inflammatory peptide angiotensin II (Ang II) to the stress-ameliorating and neuro-protective peptide angiotensin 1–7 (Ang 1–7), is broadly expressed in the striatum. Cholinergic interneurons (CIN) of the striatum, known to express multiple peptide receptors also co-expressed the corresponding and functionally opposing receptors angiotensin type 1 receptor (AT1R) for Ang II and mas receptor (MasR) for Ang 1–7. Accordingly, the conditional, semi acute ablation and/or local pharmacological inhibition of ACE2 increased the frequency of acetylcholine (ACh) bursts, reduced the amplitude of dopamine (DA)-modulated pausing of CIN activity, and disrupted the temporal coordination of extracellular levels of ACh and DA during burst events. Further, ablation of ACE2 in the DLS biased directional movement and impaired motor skill learning. Coinjection of the AT1R inhibitor losartan and the dopamine D2 receptor (D2R) agonist quinpirole reduced steady state level cholinergic activity in an additive manner, and proximity ligation supported close spatial association of AT1R and D2R on CINs. Together our study provides evidence that striatal produced ACE2 impinges on the dynamics of ACh and DA and impacts action selection and motor learning through functional and structural interactions of peptidergic and dopaminergic signaling on CIN.

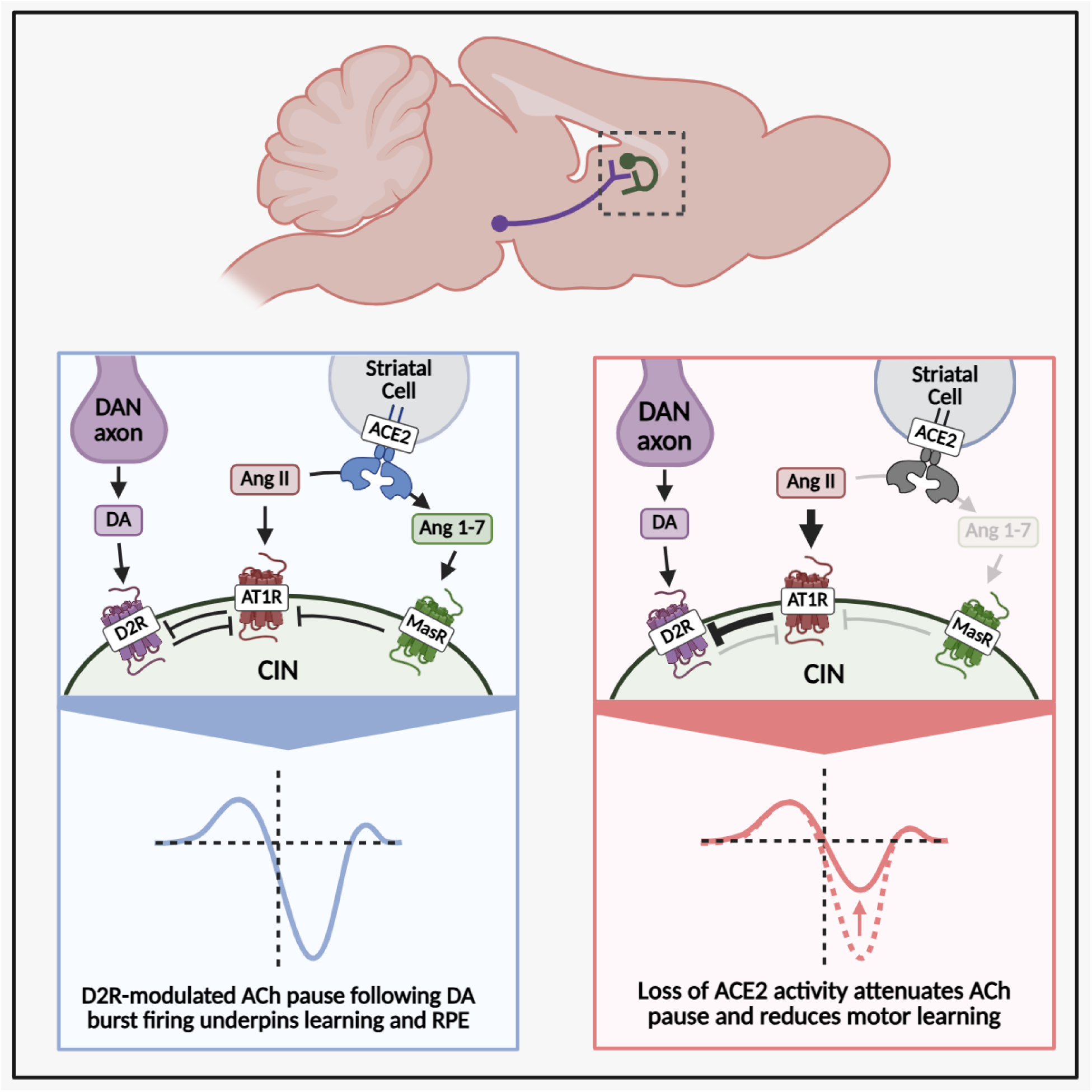
Graphical Abstract.

## Introduction

Neuropeptides constitute the largest, most diverse and evolutionary oldest group of neural messengers^1^. They play conserved roles in the control of behavior, learning, and motivational states^2–4^. Biophysically, neuropeptides introduce unique properties into neural circuits. Their slower diffusion, prolonged receptor activation, and vulnerability to depletion when release rates exceed synthesis rates expand the temporal dynamics of neuromodulation.^5–7^ These characteristics allow neuropeptide-mediated circuits to operate across broader timescales, endowing them with functional flexibility that circuits relying solely on small-molecule transmitters lack. It is now well established that most neurons across animal phyla co-secrete neuropeptides in addition to small molecule transmitters and co-express receptors for multiple neuropeptides.^8,9^ Despite their ubiquity, still little is known about how networks of neuropeptide pathways are organized and interact among themselves or with small molecule transmitters in the brain.

The striatum, the input structure of the basal ganglia, is a hub for associative learning where anti correlated fluctuations in dopamine (DA) and acetylcholine (ACh) dynamically regulate behavior^10,11^: Phasic DA release is widely understood as a teaching or reinforcement signal encoding positive reward prediction errors and driving synaptic potentiation.^12–17^ In contrast, ACh signals, especially their elevations, have been implicated in extinction learning, cue devaluation, action interruption and behavioral flexibility^18^ providing the counterbalance to reinforced associations when they are no longer yielding reward. This anti-correlated physiology is particularly prominent in the dorsolateral striatum (DLS).^12^ Here, transient ACh dips (also called the “conditioned pause”) create a window during which bursts of DA release can induce synaptic plasticity.^15,16^ During extinction, the pattern reverses: ACh rises while DA decreases, supporting the unlearning of cues that are no longer rewarded.^18^ Thus, ACh secreting cholinergic interneurons (CIN) have emerged as critical regulators of learning in the striatum whose inputs powerfully gate and contextualize DA signals. Accordingly, CIN receive input from over 40 neuromodulatory, mostly peptidergic, systems, many of which remain poorly characterized.^19,20^ This complex signaling landscape is thought to be a critical mediator of the striatum’s ability to acquire and express context appropriate behaviors.^19–23^

We find here that the mono-carboxypeptidase angiotensin-converting enzyme 2 (ACE2) is broadly expressed in the DLS. Among the about 100 known neuroactive peptides in the mammalian brain^24^, ACE2 takes part in the production and/or processing of members of the dynorphin-, neurotensin-, β-casomorphin-, kinetensin-, grehlin– and amyloid-β peptide families, many of which engage receptors known to be expressed by CIN.^25–29^ In addition, ACE2 converts the pro-inflammatory peptide angiotensin II (Ang II) to the stress-ameliorating and neuroprotective angiotensin 1–7 (Ang 1–7) of the renin angiotensin system (RAS) (Table 1) and we report here, that CIN co-express the functionally opposing receptors angiotensin 1 receptor (AT1R), which binds Ang II and MasR, which binds Ang 1-7. Thus, given its broad substrate range, ACE2 activity in the striatum is poised to effect the integration of multiple neuromodulatory inputs that converge on CIN.

**Table 1.**
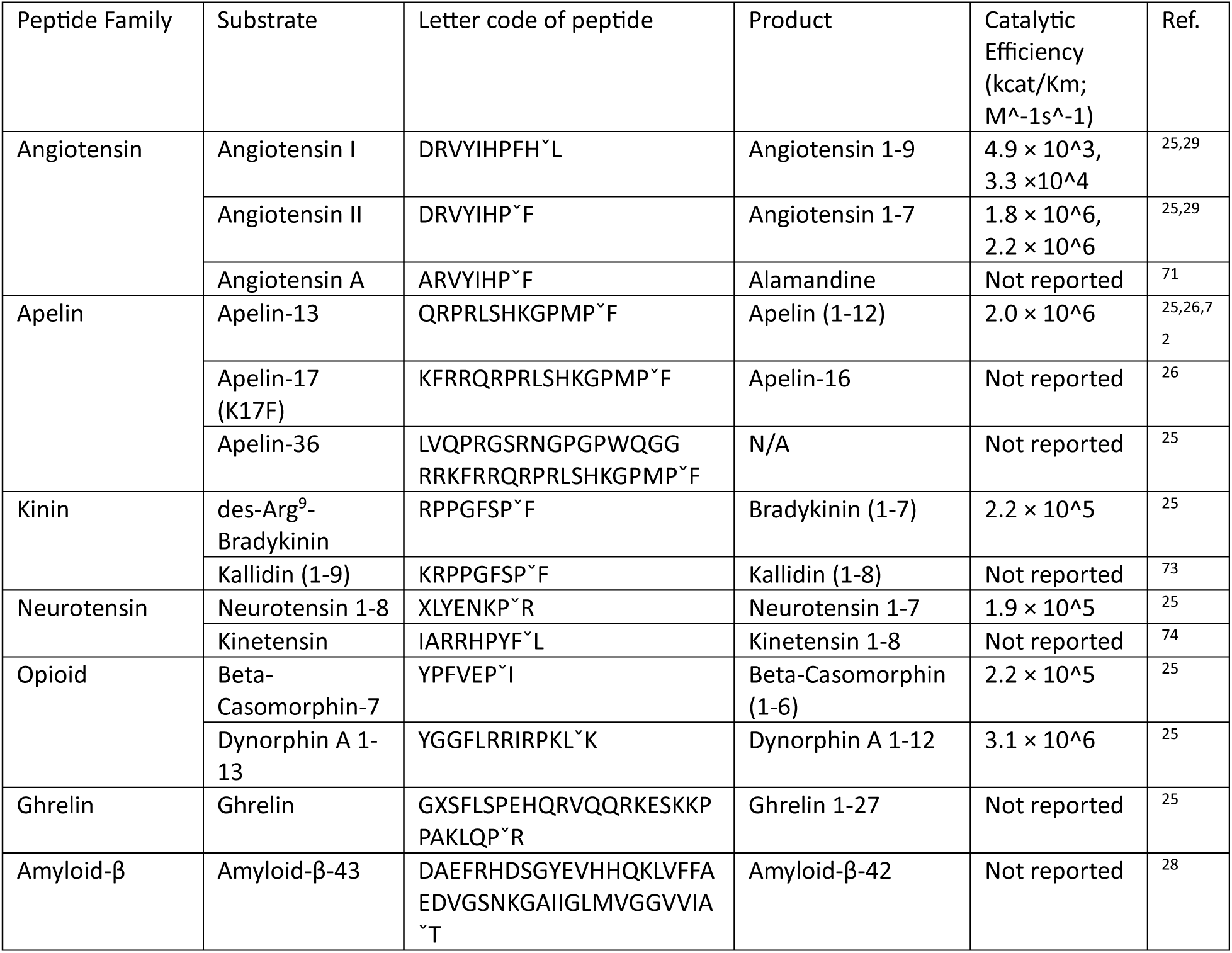
Substrates and products of angiotensin converting enzyme 2 (ACE2)

Here, revealing the physiological relevance of ACE2 enzymatic activity in the striatum, we found that both local, viral-mediated, Cre dependent ablation of ACE2 as well as intrastriatal injection of the ACE2 inhibitor MLN4760 led to a reduction in the amplitude of cholinergic pauses associated with DA release, and disturbed temporal coordination of DA and ACh signals. Accordingly, ablation of ACE2 in the DLS produced changes in motor learning. As an example of a mechanism by which ACE2 dependent peptidergic signaling can influence CIN physiology and impinge on small molecule neurotransmission, we present evidence that AT1R heteromerizes with dopamine 2 receptors (D2R) on CIN blunting D2R dependent inhibition of CIN. Overall, our findings reveal that striatal ACE2 enzymatic activity is a critical determinant of CIN physiology in the adult basal ganglia and effects learning.

## Results

### ACE2 is expressed broadly and AT1R and MasR are co-expressed in Cholinergic Interneurons in the Dorsolateral Striatum

Previous studies have identified components of the canonical renin angiotensin system (RAS) in specific basal ganglia cell types, including dopaminergic neurons in the substantia nigra ^30^ and medium spiny neurons (MSNs) in the striatum.^31^ These studies demonstrated expression of angiotensinogen (Agt), angiotensin II type 1 receptor (AT1R), angiotensin II type 2 receptor (AT2R), and the prorenin receptor (PRR). However, components of the noncanonical RAS pathway have not been thoroughly investigated, leaving open questions about the full spectrum of RAS expression and the range of striatal cell types, such as striatal interneurons, which are receptive to angiotensingeric signaling.

We examined the expression of RAS components in the dorsolateral striatum (DLS) using triple immunofluorescence staining of fixed striatal sections from adult wild-type (WT) mice. Sections were stained for DAPI (nuclei), NeuN (pan-neuronal), ChAT (cholinergic interneurons), and one of the following RAS-related proteins: angiotensin converting enzyme 2 (ACE2), AT1R, AT2R, the mas receptor (MasR), angiotensin IV receptor (AT4R), or mas-related G protein-coupled receptor type D (MrgDR).

All RAS components assessed, except for AT2R, were detected above background in DLS neurons (Fig. 1B-G). The lack of AT2R staining within this region was not attributable to antibody performance, as validation in midbrain DANs, a cell type previously shown to express AT2R^30^, confirmed antibody specificity (<u>Fig. S1B</u>). ACE2 showed broad expression throughout the DLS, localizing to both NeuN-positive and NeuN-negative cells (Fig. 1B – top row, <u>Fig S1C</u>), indicating that its distribution is not restricted to a specific neuronal population or subregion. Several of the RAS receptors showed high expression in a subset of large, sparsely distributed neurons, which co-labeled with ChAT, identifying this subpopulation as cholinergic interneurons (CINs) (Fig. 1B). CINs expressed higher levels of AT1R, MasR, and AT4R compared to non-CIN neurons, whereas MrgDR expression did not differ significantly between these populations (Fig. 1H-K). These differences were consistent across multiple animals (<u>Fig. S1D-E</u>). Furthermore, a greater proportion of CINs expressed AT1R, MasR, and MrgDR, with no difference in AT4R (Fig. 1L-O). Notably, 100% of CINs expressed AT1R and MasR, compared to 93.01 ± 0.41% and 84.84 ± 0.73% of non-cholinergic neurons, respectively (mean ± SEM) (Fig. 1L-M).

**Figure 1.**
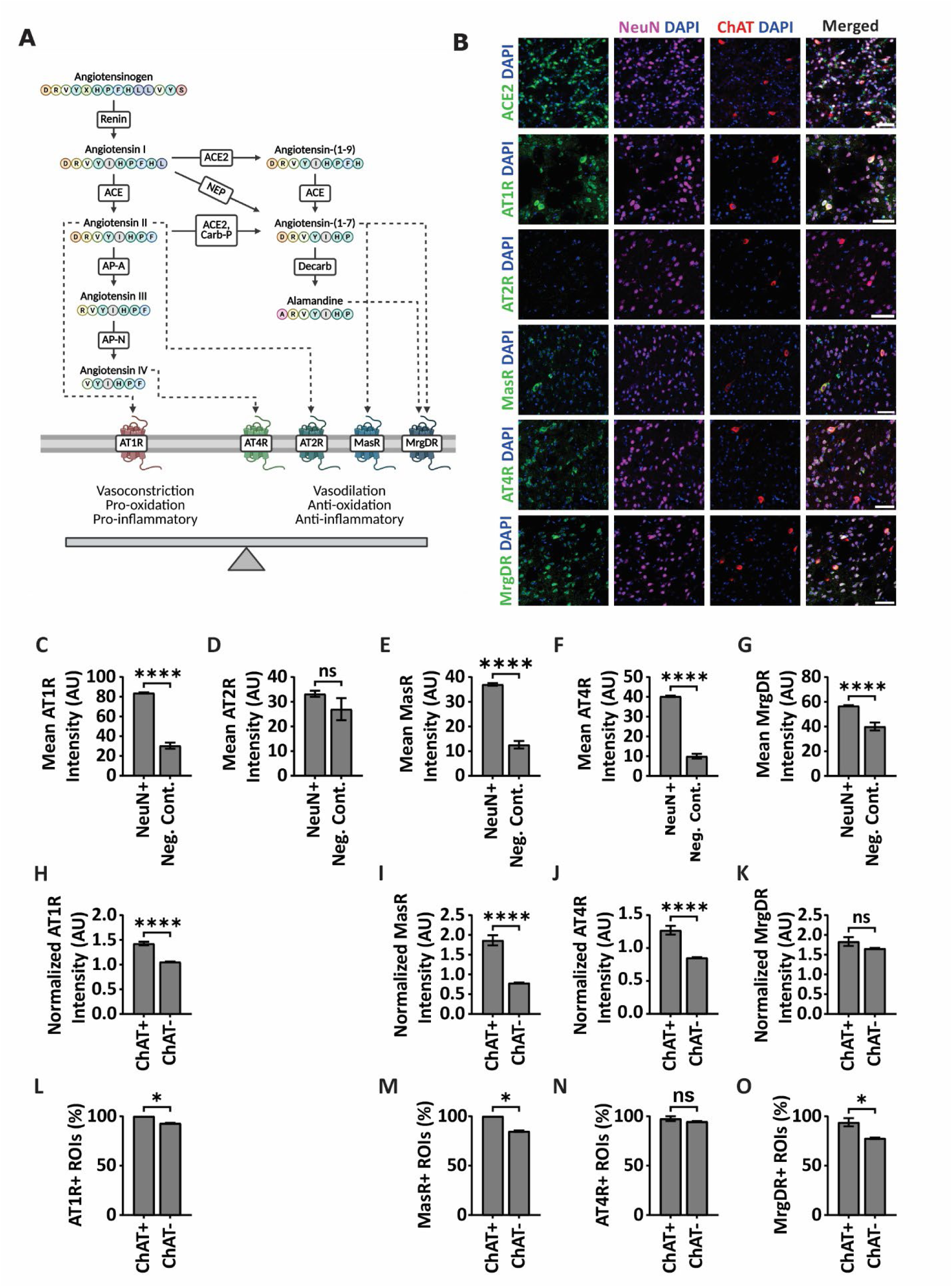
Cell-type-specific expression of RAS components in the dorsolateral striatum. (A) An overview of the renin angiotensin system (RAS) signaling pathway. Angiotensinogen (Agt), the precursor peptide, is cleaved by renin to produce angiotensin I (Ang I), which is then converted by angiotensin-converting enzyme (ACE) into angiotensin II (Ang II), the first active peptide. Ang II signals through two opposing receptors: the angiotensin II type 1 receptor (AT1R) and the angiotensin II type 2 receptor (AT2R). Ang II can be cleaved by angiotensin-converting enzyme 2 (ACE2) or carboxypeptidase P (Carb-P) into angiotensin-(1-7) (Ang 1-7), which primarily binds the mas receptor (MasR) and, to a lesser degree, the mas-related G-protein coupled receptor member D (MrgDR). Ang 1-7 can be further decarboxylated to form alamandine, a ligand of MrgDR. Alternatively, Ang II can be converted into angiotensin III (Ang III) by aminopeptidase A (AP-A), and subsequently into angiotensin IV (Ang IV) by aminopeptidase N (AP-N). Ang IV can bind angiotensin IV receptor (AT4R), also known as insulin-regulated aminopeptidase (IRAP). Functionally, the RAS is divided into two arms: the classical/canonical arm, mediated by AT1R, promotes vasoconstriction, increased reactive oxygen species production, inflammatory signaling and neurodegeneration; and the non-classical/non-canonical arm, mediated by AT2R, MasR, MrgDR, and AT4R, promotes vasodilation, reduced oxidation and inflammatory signaling, and neuroprotection. **(B)** Representative confocal images of dorsolateral striatum (DLS) sections from an adult wild-type (WT) male mouse. Sections were co-stained with NeuN (magenta), ChAT (red), DAPI (blue), and one of the following RAS proteins (green): ACE2, AT1R, AT2R, MasR, AT4R, or MrgDR. Scale bars: 50 µm. **(C-G)** Quantification of mean fluorescence intensity for RAS receptors in NeuN+ neurons of the dorsolateral striatum (DLS). Fluorescence intensity was measured in NeuN+ regions of interest (ROIs) and compared to negative control ROIs located in adjacent DAPI-negative areas. Data are presented as mean ± SEM. Due to violations of normality, statistical comparisons were performed using a two-tailed Mann–Whitney U test. Sample sizes indicate ROIs from one WT adult male (NeuN+ vs. control): **(C)** AT1R (n = 3,857 vs. 30, **** p < 0.0001), **(D)** AT2R (n = 507 vs. 25, p = 0.1422), **(E)** MasR (n = 2,456 vs. 30, **** p < 0.0001), **(F)** AT4R (n = 2,491 vs. 25, **** p < 0.0001), **(G)** MrgDR (n = 2,721 vs. 30, **** p < 0.0001). **(H-K)** NeuN-normalized mean fluorescence intensities of RAS receptors were quantified in ChAT+ NeuN+ and ChAT-NeuN+ ROIs. Fluorescence values were normalized to the NeuN signal within each ROI to control for potential variability in staining and imaging conditions. Due to violations of normality, statistical comparisons were performed using a two-tailed Mann–Whitney U test. Sample sizes indicate ROIs from one WT adult male (ChAT+ NeuN+ vs. ChAT– NeuN+): **(H)** AT1R (n = 68 vs. 3,789, **** p < 0.0001), **(I)** MasR (n = 34 vs. 2,422, **** p < 0.0001), **(J)** AT4R (n = 41 vs. 2,450, **** p < 0.0001), **(K)** MrgDR (n = 33 vs. 2,688, p = 0.1015). **(L-O)** Proportion of receptor-positive ROIs were calculated for each RAS receptor and compared between ChAT+ NeuN+ and ChAT-NeuN+ ROIs. ROIs were classified as receptor-positive if the mean fluorescence intensity exceeded a threshold defined by the maximum fluorescence intensity measured in the negative control ROIs. Due to violations of normality, statistical comparisons were performed using a two-tailed Mann–Whitney U test. Sample sizes indicate ROIs from one WT adult male (ChAT+ NeuN+ vs. ChAT– NeuN+): **(L)** AT1R (n = 68 vs. 3,789, *p = 0.0241), **(M)** MasR (n = 34 vs. 2,422, *p = 0.0119), **(N)** AT4R (n = 41 vs. 2,450, p = 0.5214), **(O)** MrgDR (n = 33 vs. 2,688, *p = 0.0315). See <u>Table S1</u> for full statistical details.

These findings indicate that neurons in the DLS engage both in canonical (AT1R-mediated) and non-canonical (MasR/AT4R/MrgDR-mediated) RAS signaling. The absence of AT2R, a classical counter-regulator of AT1R signaling, underscores the critical role of ACE2, whose enzymatic activity shifts RAS activity from AT1R-driven pro-inflammatory signaling towards MasR/AT4R/MrgDR-mediated anti-inflammatory signaling. Thus, our expression studies indicated that most neuronal subtypes of the striatum are capable of being influenced by RAS peptides, that in part might be produced locally. We tested this possibility by focusing on CIN.

### Local ACE2 Enzymatic Activity Modulates Acetylcholine (ACh) Release and the Temporal Coordination of ACh and Dopamine (DA) in the DLS

CINs are the primary contributor of acetylcholine (ACh) tone in the striatum, which acts as a key modulatory signal on MSNs and dopamine (DA) axons from the midbrain.^32^ The complex reciprocal modulation of ACh and DA is particularly critical for key striatal functions such as motor selection and decision-making.^33,34^ While prior studies have shown that RAS signaling can modulate striatal neurotransmitter release,^35–38^ its specific effects on ACh release and ACh-DA release dynamics remain uncharacterized. Here we employed fiber photometry (FP) to monitor ACh and DA release using genetically encoded G protein-coupled receptor (GPCR) activation based (GRAB) sensors ‒ specifically AAV-delivered ACh3.0 (green M3 receptor-based ACh sensor)^39^ and rDA1h (red D2 receptor-based DA sensor – Fig. 2A-C).^40^

**Figure 2.**
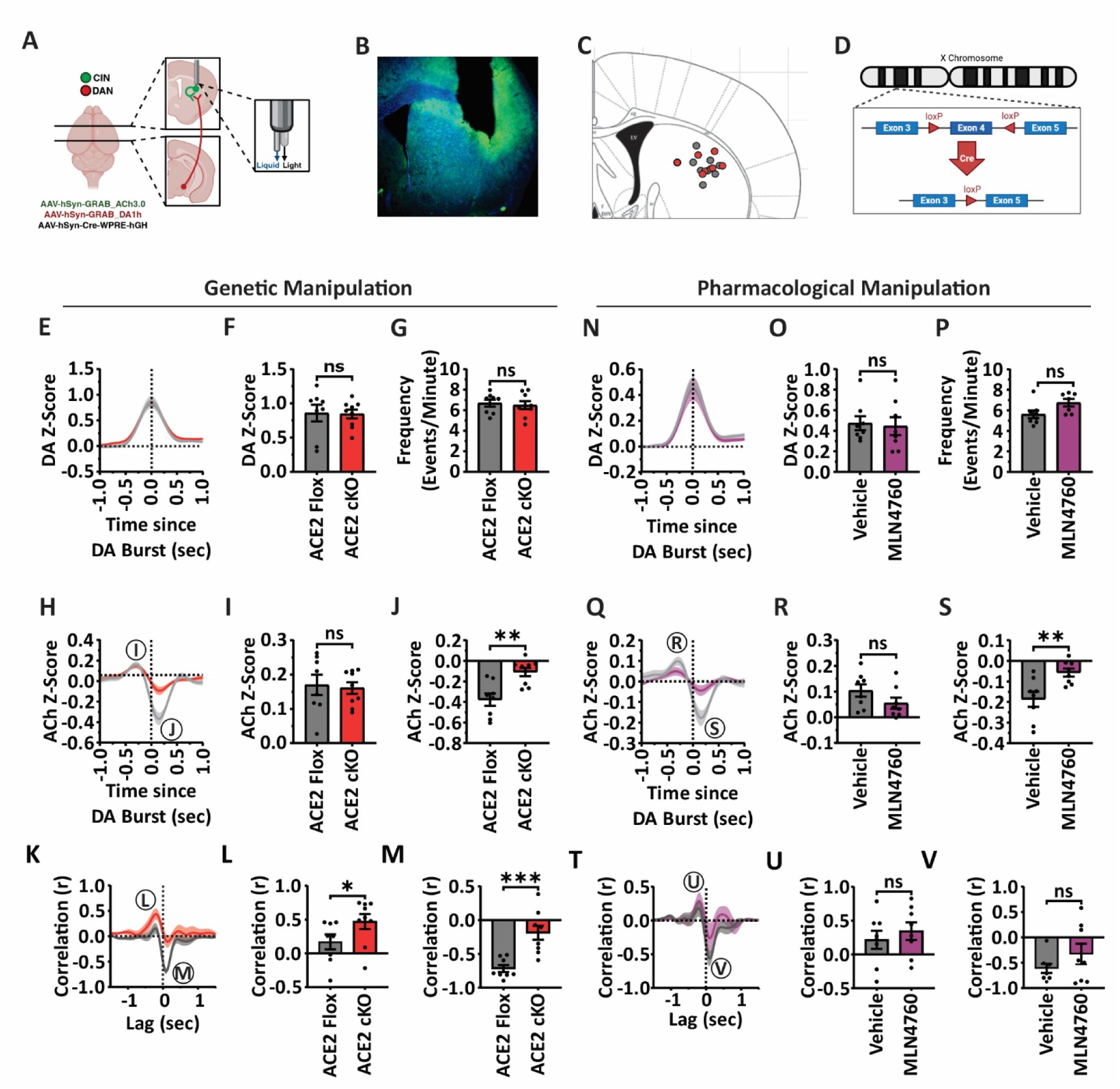
Genetic Loss and Pharmacological Inhibition of ACE2 Enzymatic Activity Disrupts the Acetylcholine Pause Following Dopamine Burst Firing in the DLS. (A) Graphical overview of fiber photometry experimental setup. Mice were implanted with a combined optic fiber– injection cannula targeting the DLS and injected with AAV-hSyn-GRAB_ACh3.0 and AAV-hSyn-GRAB-DA1h, with or without AAV-hSyn-Cre-WPRE-hGH. **(B)** Representative fluorescence microscopy image depicting the fiber optic implantation site and expression of AAV-hSyn-GRAB_ACh3.0. **(C)** Mapped locations of fiber optic implant sites determined by postmortem analysis. **(D)** Schematic of the ACE2 conditional knockout (ACE2 cKO) model. Exon 4 of the ACE2 gene is flanked by loxP sites and is excised upon Cre recombinase expression, resulting in production of an (enzymatically inactive ACE2 protein. **(E–G)** DA burst events were compared between ACE2 Flox control mice (grey; n = 8) and ACE2 cKO mice (red; n = 9). Data are presented as mean ± SEM. Statistical tests were selected based on normality and variance assumptions. **(F)** Amplitude of DA burst events (unpaired two-tailed t-test: p = 0.9147). **(G)** Frequency of DA burst events (unpaired two-tailed t-test: p = 0.7055). **(H-J)** ACh traces during DA burst events were compared between ACE2 Flox control mice (grey; n = 8) and ACE2 cKO mice (red; n = 9). **(I)** Maximum ACh peak amplitude preceding the DA burst (unpaired two-tailed t-test: p = 0.7834). **(J)** Minimum ACh dip amplitude following the DA burst (unpaired two-tailed t-test: **p = 0.0019). **(K-M)** Cross-correlations of ACh lagged relative to DA bursts were compared between ACE2 Flox control mice (grey; n = 8) and ACE2 cKO mice (red; n = 9). **(L)** Maximum correlation magnitude (Mann–Whitney U test: *p = 0.0206). **(M)** Minimum correlation magnitude (unpaired two-tailed t-test: ***p = 0.0005). **(N-P)** DA burst events were compared within ACE2 Flox control mice (n = 7–8) following local injections of cortex buffer vehicle (grey) vs. 0.1 mM MLN4760 (purple). **(O)** Amplitude of DA burst events (paired two-tailed t-test: p = 0.8132). **(P)** Frequency of DA burst events (paired two-tailed t-test: p = 0.0597). **(Q-S)** ACh traces during DA burst events were compared within ACE2 Flox control mice (n = 8) following local injections of vehicle (grey) vs. MLN476 (purple). **(R)** Maximum ACh peak amplitude preceding the DA burst (paired two-tailed t-test: p = 0.0597). **(S)** Minimum ACh dip amplitude following the DA burst (paired two-tailed t-test: **p = 0.0072). **(T–V)** Cross-correlations of ACh lagged relative to DA bursts were compared within ACE2 Flox control mice (n = 8) following local injections of vehicle (grey) vs. MLN4760 (purple). **(U)** Maximum correlation magnitude (paired two-tailed t-test: p = 0.5611). **(V)** Minimum correlation magnitude (paired two-tailed t-test: p = 0.2776). Data represent pooled results from two independent experimental cohorts with consistent findings. See <u>Table S2</u> for full statistical details.

ACE2 activity was manipulated via either genetic ablation of ACE2 within the DLS or local pharmacological inhibition. To accomplish this, we implanted Doric optical fiber multiple fluid injection cannulas in the DLS of ACE2 conditional knockout (cKO) mice in which exon 4 of the *Ace2* gene is flanked by loxP sites allowing for Cre-mediated local deletion, resulting in an enzymatically inactive ACE2 protein (Fig. 2D).^41^ For genetic manipulation, the animals received injections of recombinant AAV9 particles via the fluid injection port of the optofluid implants leading to the expression of Cre from the a-syn promoter (Addgene #105553) and neuronal-specific manipulation of the *Ace2* gene. Selective *Ace2* ablation in the striatum was confirmed by PCR of dissected tissue (<u>Fig. S2A</u>). For the pharmacological manipulation, the clinically used ACE2 inhibitor MLN-4760 was injected through the same port during FP recording to acutely inhibit all local ACE2 activity.

All recording sessions were performed in head-fixed animals and began with a 15-minute habituation period, followed by continuous GRAB fluorescence recording for the remainder of the session. For local drug delivery experiments, a 15-minute baseline was recorded prior to infusion. Mice received either 0.5 µL cortex buffer (vehicle) or 0.1 mM MLN4760, delivered at 4 µL/min through the cannula. Recordings continued for 30 minutes after the end of infusion. DA and ACh burst events were defined as transient increases exceeding 3 median absolute deviations (MADs) above a 150-second moving median baseline.^42^ Three-second windows surrounding each event were extracted and averaged for both sensor channels to characterize CIN–DAN release dynamics (<u>Fig. S3</u>).

We first identified DA burst events from these traces and found that DA burst amplitude and frequency were not significantly affected by ACE2 genetic ablation (Fig. 2E-G) or acute pharmacological inhibition (Fig. 2N-P). We next examined ACh dynamics coinciding with the identified DA bursts. In the DLS, a triphasic response in cholinergic interneurons, including an initial glutamatergic excitation, a D2 receptor–dependent pause, and a rebound excitation, often coincides with phasic dopamine release.^10,12,17,22,23,43^ The amplitude and duration of the ACh peak preceding the DA burst, often reflecting the initial excitation, were not significantly altered by ACE2 genetic ablation (Fig. 2H-I; <u>Fig. S4A</u>) or local pharmacological inhibition in either Flox controls (Fig. 2Q-R; <u>Fig. S4C</u>). However, both genetic ablation and pharmacological inhibition significantly reduced the amplitude of the subsequent ACh dip which reflects a phase of hyper polarization of CIN (Fig. 2J,S) without altering its duration (<u>Fig. S4B,D</u>).

To assess the temporal relationship between ACh and DA signals during these DA burst events in the DLS, we performed cross-correlation analysis. When ACh signals were lagged relative to DA burst events, the cross-correlation exhibited a positive peak at negative lags and a negative peak at positive lags (Fig. 2K,T), reflecting ACh activity preceding the DA release into the striatum, which in turn is followed by an ACh pause, matching the well-described triphasic response of CIN during phasic DA release associated with learning and characterized across multiple techniques including electrophysiology recordings and fiber photometry.^11,43–45^ Cross-correlation revealed that ACE2 genetic ablation increased the positive correlative relationship between the DA burst event and the preceding ACh peak compared to controls (Fig 2K-L). Additionally, ACE2 genetic ablation strongly reduced the anti-correlative relationship between the DA burst and the ACh dip that followed (Fig. 2M).

While the ablation and pharmacological inhibition of ACE2 at the recording site led to similar a reduction in amplitude of the ACh dip (Fig. 2J,S) without altering the DA burst amplitude (Fig. 2F,O), the pharmacological inhibition of ACE2 did not affect the negative correlation at a positive lag (Fig. 2T,V). The different outcomes revealed by the two approaches point to different sources and functions of ACE2 targeted by the neuron biased expression of Syn-Cre and the inhibitor MLN4760 or cKO mice.

We next repeated the event detection analyses above but isolated ACh burst events instead of DA burst events. Similar to ACE2 effects on DA burst events, neither genetic ablation of ACE2 nor acute pharmacological inhibition significantly altered the frequency or amplitude of ACh burst events in the DLS (Fig. 3A-C<u>, J-L</u>). In contrast, genetic ablation of ACE2 significantly reduced the DA dip amplitude preceding the ACh burst without altering the subsequent DA peak amplitude (Fig. 3D-F). Acute pharmacological inhibition of ACE2 showed a similar trend towards reduced DA dip amplitude which approached significance (*p* = 0.0719) but no effect on DA peak amplitude (Fig. 3M-O).

**Figure 3.**
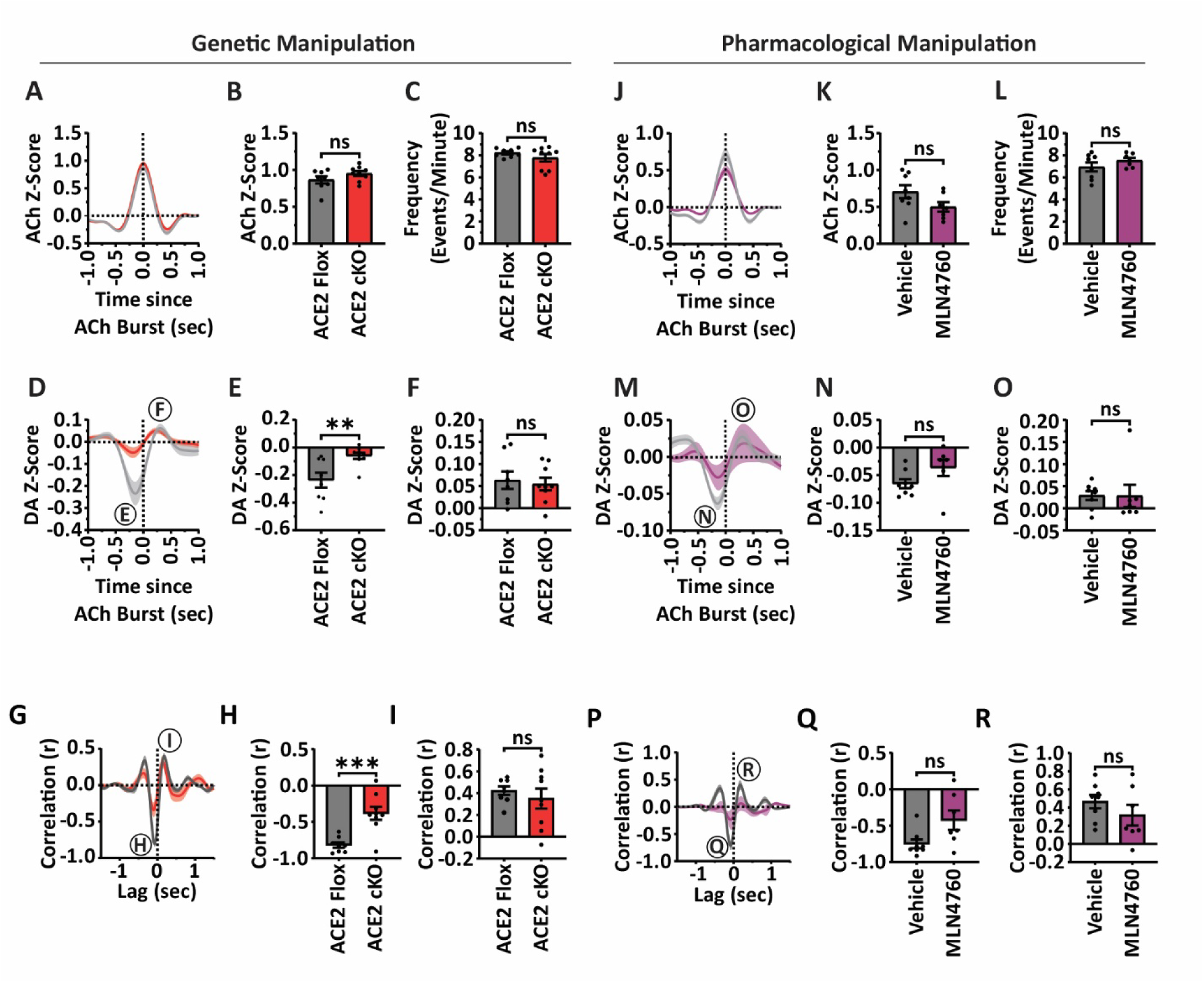
Genetic loss of ACE2 enzymatic activity, but not pharmacological inhibition, diminishes the dopamine dip preceding ACh bursts in the DLS. (A–C) ACh burst events were compared between ACE2 Flox control mice (grey; n = 8) and AAV-Cre–treated ACE2 cKO mice (red; n = 9). Data are presented as mean ± SEM. Statistical tests were selected based on normality and variance assumptions. (B) Amplitude of ACh burst events (unpaired two-tailed t-test: p = 0.1524). (C) Frequency of ACh burst events (Welch’s two-tailed t-test: p = 0.2404). (D–F) ACh burst events were compared within ACE2 Flox control mice (n = 7) following local injections of cortex buffer vehicle (grey) vs. 0.1 mM MLN4760 (purple). (E) Amplitude of ACh burst events (paired two-tailed t-test: p = 0.1091). (F) Frequency of ACh burst events (Wilcoxon matched-pairs signed rank test: p = 0.5781). (G–I) DA traces during ACh burst events were compared between ACE2 Flox controls (grey; n = 8) and ACE2 cKO mice (red; n = 9). (H) Minimum DA dip amplitude preceding the ACh burst (Welch’s two-tailed t-test: *p = 0.0142). (I) Maximum DA peak amplitude following the ACh burst (unpaired two-tailed t-test: p = 0.7177). (J–L) DA traces during ACh burst events were compared within ACE2 Flox control mice (n = 7) following local injections of vehicle (grey) vs. MLN4760 (purple). (K) Minimum DA dip amplitude (paired two-tailed t-test: p = 0.0719). (L) Maximum DA peak amplitude (Wilcoxon matched-pairs signed rank test: p = 0.4688). (M–O) Cross-correlations of DA lagged relative to ACh bursts were compared between ACE2 Flox control mice (grey; n = 8) and ACE2 cKO mice (red; n = 9). (N) Minimum correlation magnitude (Welch’s two-tailed t-test: ***p = 0.0010). (O) Maximum correlation magnitude (Welch’s two-tailed t-test: p = 0.4221). (P–R) Cross-correlations of DA lagged relative to ACh bursts were compared within ACE2 Flox control mice (n = 7) following local injections of vehicle (grey) vs. MLN4760 (purple). (Q) Minimum correlation magnitude (paired two-tailed t-test: p = 0.0945). ® Maximum correlation magnitude (paired two-tailed t-test: p = 0.4608). Data represent pooled results from two independent experimental cohorts with consistent findings. See Table S3 for full statistical details.

We next assessed the temporal correlation between ACh and DA during ACh bursts. When DA signals were lagged relative to ACh burst events, the cross-correlation showed a negative peak at negative lags and a positive peak at positive lags (Fig. 3G,P), consistent with an initial dip in extracellular DA predicting ACh burst firing, followed by a rebound in DA release. ACE2 ablation significantly reduced the anti-correlative relationship between ACh and DA when DA was negatively lagged, representing a weakening of the predictive relationship of a DA dip being followed by an ACh peak (Fig. 3G-H). This effect was not replicated by acute pharmacological inhibition of ACE2 (Fig. 3P,Q). The positive correlative relationship between ACh and DA when DA was positively lagged was not significantly altered by ACE2 ablation or pharmacological inhibition (Fig. 3I,R).

Because ablation and pharmacological inhibition of ACE2 did not result in equivalent effects on all features analyzed suggesting the existence of distinct pools of ACE2, we combined the two approaches by treating ACE2 cKO mice with MLN4760. When isolating DA burst events, the additional pharmacological inhibition of ACE2 in the cKO mice had no effect on DA burst amplitude or frequency (Fig. 4A-C). Likewise, DA burst-associated ACh dynamics as well as ACh-DA cross correlations were unaffected by the additional inhibition of ACE2 (Fig. 4D-I). However, when isolating ACh bursts, we found the frequency of these events markedly increased (Fig. 4L), an effect not seen with either ablation or inhibition alone (Fig. 3C,L). Further, while the dynamics of DA release associated with the ACh burst, including the DA dip prior and the peak after the ACh burst, were not affected (Fig. 4M-O), the cross-correlation analysis between DA and ACh revealed a qualitative difference. When DA signals were lagged relative to ACh burst events, the cross-correlation showed a negative peak at negative lags and a positive peak at positive lags as seen before (Fig. 4P, Fig. 3G-P), consistent with an initial dip in extracellular DA predicting ACh burst firing, followed by a rebound in DA release. However, in addition to this pattern, the further inhibition of ACE2 in ACE2 cKO mice resulted in a wide second positive peak at positive lags suggesting that ACh bursts in this condition predict long duration DA activity that is not seen with ACE2 ablation alone (Fig. 3G).

**Figure 4.**
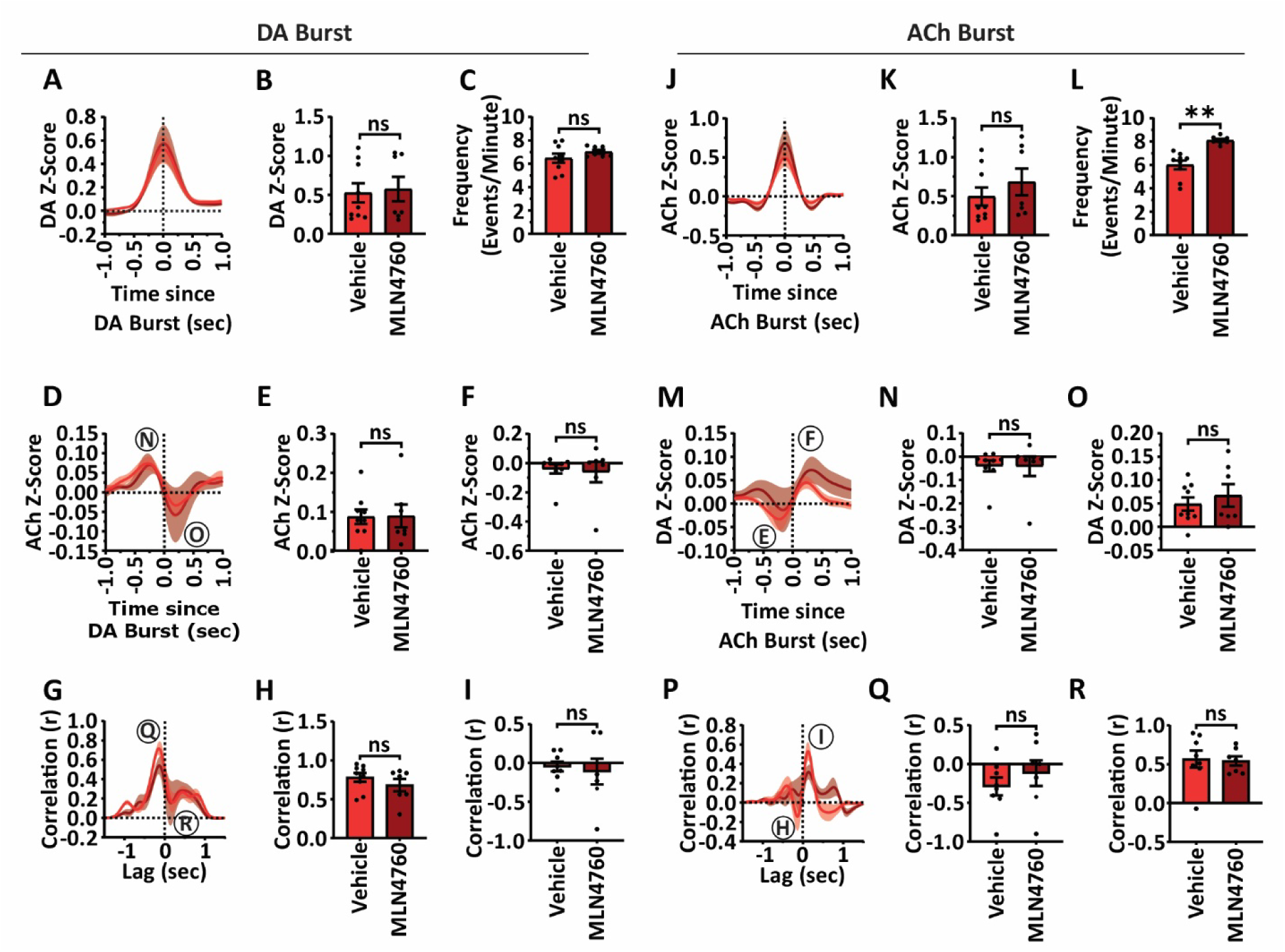
ACE2 Inhibitor MLN4760 Produces Minimal Effects in ACE2 Conditional Knockout Mice. (A-C) DA burst events were compared within ACE2 cKO mice (n = 7) following injections of vehicle (red) vs. MLN4760 (dark red). **(B)** DA burst amplitude (paired two-tailed t-test: p = 0.5145). **(C)** DA burst frequency (paired two-tailed t-test: p = 0.5841). **(D-F)** ACh traces during DA burst events were compared within ACE2 cKO mice (n = 7) after injections of vehicle (red) vs. MLN4760 (dark red). **(E)** Maximum ACh peak amplitude preceding the DA burst (paired two-tailed t-test: p = 0.9639). **(F)** Minimum ACh dip amplitude following the DA burst (paired two-tailed t-test: p = 0.8720). **(G-I)** Cross-correlations of ACh lagged relative to DA bursts were compared within ACE2 cKO mice (n = 6–7) after injections of vehicle (red) vs. MLN4760 (dark red). **(H)** Maximum correlation magnitude (paired two-tailed t-test: p = 0.3810). **(I)** Minimum correlation magnitude (paired two-tailed t-test: p = 0.5708). **(J-L)** ACh burst events were compared within ACE2 cKO mice (n = 7) following local injections of cortex buffer vehicle (red) vs. 0.1 mM MLN4760 (dark red). Data are presented as mean ± SEM. **(K)** ACh burst amplitude (paired two-tailed t-test: p = 0.0905). **(L)** ACh burst frequency (Wilcoxon matched-pairs signed-rank test: *p = 0.0156). **(M-O)** DA traces during ACh burst events were compared within ACE2 cKO mice (n = 7) after injections of vehicle (red) vs. MLN4760 (dark red). **(N)** Minimum DA dip amplitude preceding the ACh burst (paired two-tailed t-test: p = 0.6400). **(O)** Maximum DA peak amplitude following the ACh burst (paired two-tailed t-test: p = 0.6701). **(P-R)** Cross-correlations of DA lagged relative to ACh bursts were compared within ACE2 cKO mice (n = 7) after injections of vehicle (red) vs. MLN4760 (dark red). **(Q)** Minimum correlation magnitude (paired two-tailed t-test: p = 0.3517). **(R)** Maximum correlation magnitude (paired two-tailed t-test: p = 0.9557).Data represent pooled results from two independent experimental cohorts with consistent findings. See <u>Table S4</u> for full statistical details.

Collectively, these results indicate that local ACE2 enzymatic activity, possibly from multiples sources, plays a critical and specific role in organizing DA-ACh interactions in the DLS.

### AT1R Inhibition Alters D2R-Mediated CIN Activity and associates with D2R on CIN

Dopamine D2 receptor (D2R) signaling shapes the duration and amplitude of the CIN pause.^23^ D2Rs can form functional heteromers with AT1Rs leading to cross modulation at the level of ligand binding between dopamine and Ang II.^46^ In heterologous systems, co-expression of AT1R and D2R increases D2R sensitivity to low concentrations of the D2R agonist quinpirole, while attenuating its response at higher concentrations compared to D2R expression alone.^46^ Our functional observation that ACE2 inhibition or ablation, which should increase extracellular levels of Ang II and promote AT1R activation, reduces dopamine-associated cholinergic pauses alludes to the possibility that functional interaction between AT1R and D2 may act as the mechanism of action for this attenuation.

To investigate whether AT1R activity modulates D2R function in CINs, adult mice received intraperitoneally (I.P.) injected of saline, 5mg/kg of the AT1R inhibitor losartan, 3mg/kg of the D2R agonist quinpirole, or a combination of losartan and quinpirole. Following injection, mice were placed in the open field (OF) for 30 minutes, then perfused for histological analysis. Striatal sections were stained for phosphorylated ribosomal protein S6 (p-rpS6), a marker of CIN activity.^47^

In the open field, quinpirole ‒ alone or in combination with losartan ‒ significantly reduced total distance moved and mean velocity as compared to saline or losartan (<u>Fig. S6</u>), consistent with previous reports demonstrating its suppressive effects on locomotor activity.^48^ No significant differences were observed between saline– and losartan-treated mice, nor between quinpirole alone and the combination of losartan and quinpirole. Striatal tissue from these mice were then stained for p-rpS6. Neither losartan nor quinpirole alone significantly altered p-rpS6 levels compared to saline. However, co-administration of losartan and quinpirole significantly reduced p-rpS6 expression compared to both saline and quinpirole alone (Fig. 5A-B), indicating enhanced suppression of CIN activity. Consistent with the established cross inhibitory relationship of AT1R and D2R^46^ these observations suggest that inhibiting AT1R activity, which normally blunts D2R activity, allowed for stronger D2R activation and inhibition of CIN.

**Figure 5.**
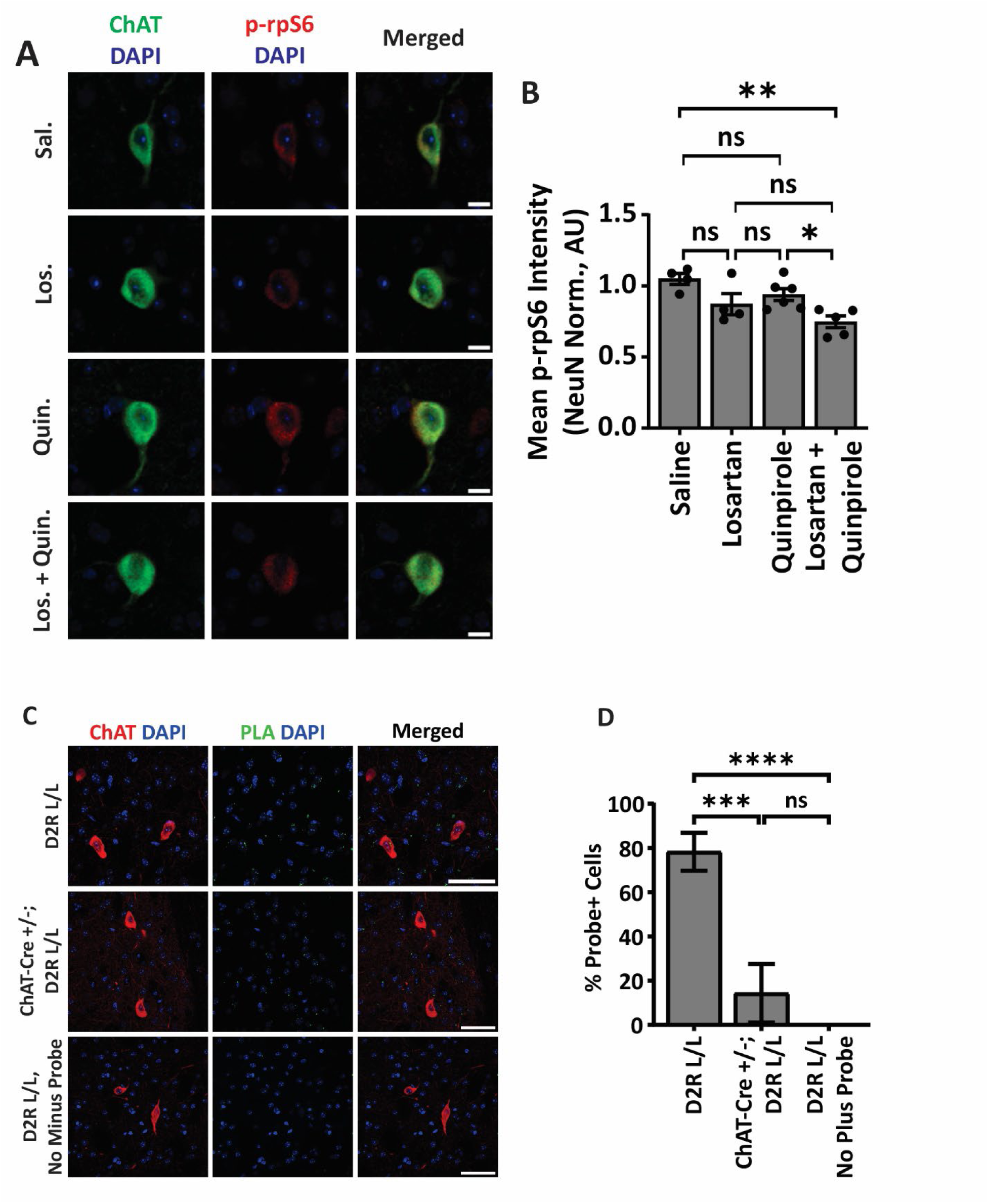
AT1R–D2R Interactions and Pharmacological Modulation of CIN Activity in the DLS. (A) Representative immunofluorescence confocal images of dorsolateral striatum (DLS) sections from adult ACE2 Flox control mice following injection with saline (Sal.), 5 mg/kg losartan (Los.), 3 mg/kg quinpirole (Quin.), or losartan + quinpirole (Los. + Quin.). Sections were co-stained for phosphorylated ribosomal protein S6 (p-rpS6, red), ChAT (green), and DAPI (blue). Scale bars: 10 µm. **(B)** Animal-averaged NeuN-normalized p-rpS6 expression in DLS CINs across the four treatment groups (saline: n = 4; losartan: n = 4; quinpirole: n = 6; losartan + quinpirole: n = 5). Data are presented as mean ± SEM. Statistical analysis was performed using one-way ANOVA (F(3,15) = 6.366, **p = 0.0054). Holm–Šídák’s multiple comparisons test: losartan + quinpirole vs. saline (**p = 0.0045), losartan + quinpirole vs. quinpirole (*p = 0.0475). No other pairwise comparisons were significant. **(C)** Representative immunofluorescence confocal images of DLS sections from adult male mice. Top and bottom: D2R L/L; middle: ChAT-Cre +/– D2R L/L. Sections underwent proximity ligation assay (PLA) to visualize AT1R–D2R heterodimers (green), followed by immunohistochemical staining for ChAT (red) and DAPI (blue). Scale bars: 50 µm. **(D)** Percentage of DLS CINs positive for AT1R–D2R dimers across genotypes and conditions. Groups include D2R L/L with both plus and minus PLA probes, ChAT-Cre +/– D2R L/L with both plus and minus PLA probes (biological control), and D2R L/L with minus PLA probe only (technical control). Data in panels A and B represent pooled results from two independent experimental cohorts with consistent findings. See <u>Table S5</u> for full statistical details.

Previous work has shown that AT1R-D2R heteromers form on rat striatal cells, however cell type specificity of these dimers remains unexplored.^46^ To determine if AT1R-D2R dimers form specifically on CINs, we used proximity ligation assay (PLA), which detects proteins localized within approximately 40nm of each other, followed by immunohistochemical staining for the CIN-marker ChAT and the pan-neuronal marker NeuN. Consistent with previous observations in the striatum^46^, roughly 70% of NeuN+ cells were positive for AT1R-D2R PLA puncta (<u>Fig. S7B</u>). Further, we found that 78.3±8.6% of CINs colocalized with PLA puncta (Fig. 5C-D). Selective ablation of D2R in CIN (ChAT-Cre +/-; D2R L/L mice) resulted in a fivefold reduction of CIN-associated PLA puncta supporting the presence of AT1R-D2R dimers on CINs in the DLS.

These observations point to the possibility that increased stimulation of AT1R by Ang II, the substrate of ACE2, could impinge on DA-dependent modulation of CIN.

### Local ACE2 Activity in the DLS Regulates Directional Motor Output and Supports Motor Learning

The preceding results suggest that local ACE2 in the DLS influences the coordination of DA and ACh signals that are critical for action selection, movement vigor, and motor learning. To assess whether ACE2 activity in the DLS regulates motor output, adult mice unilaterally implanted with Doric’s optical fiber multiple fluid injection cannula in the DLS were assessed in the open field. Prior to virus injection, mice exhibited a slight contralateral turning bias, possibly due to the unilateral implant (Fig. 6A, week 0). Within three weeks following AAV-Cre injection, mice with unilateral ACE2 cKO in the DLS developed a significant ipsilateral turn bias (Fig. 6A-B), driven by a significant increase in the number of ipsilateral (counterclockwise) turns (Fig. 6D) while contralateral (clockwise) turns remained unchanged (Fig. 6C). Notably, this rotational bias occurred without changes in overall motor output, as total distance moved and mean velocity did not differ between groups in either unilaterally (<u>Fig. S8A-B</u>) or bilaterally (<u>Fig. S8D-E</u>) manipulated mice. Likewise, ACE2 ablation did not affect angular path variability, as indicated by similar mean relative meander across groups (<u>Fig. S8C</u>). These findings suggest that ACE2 activity in the DLS supports appropriate motor action selection.

**Figure 6.**
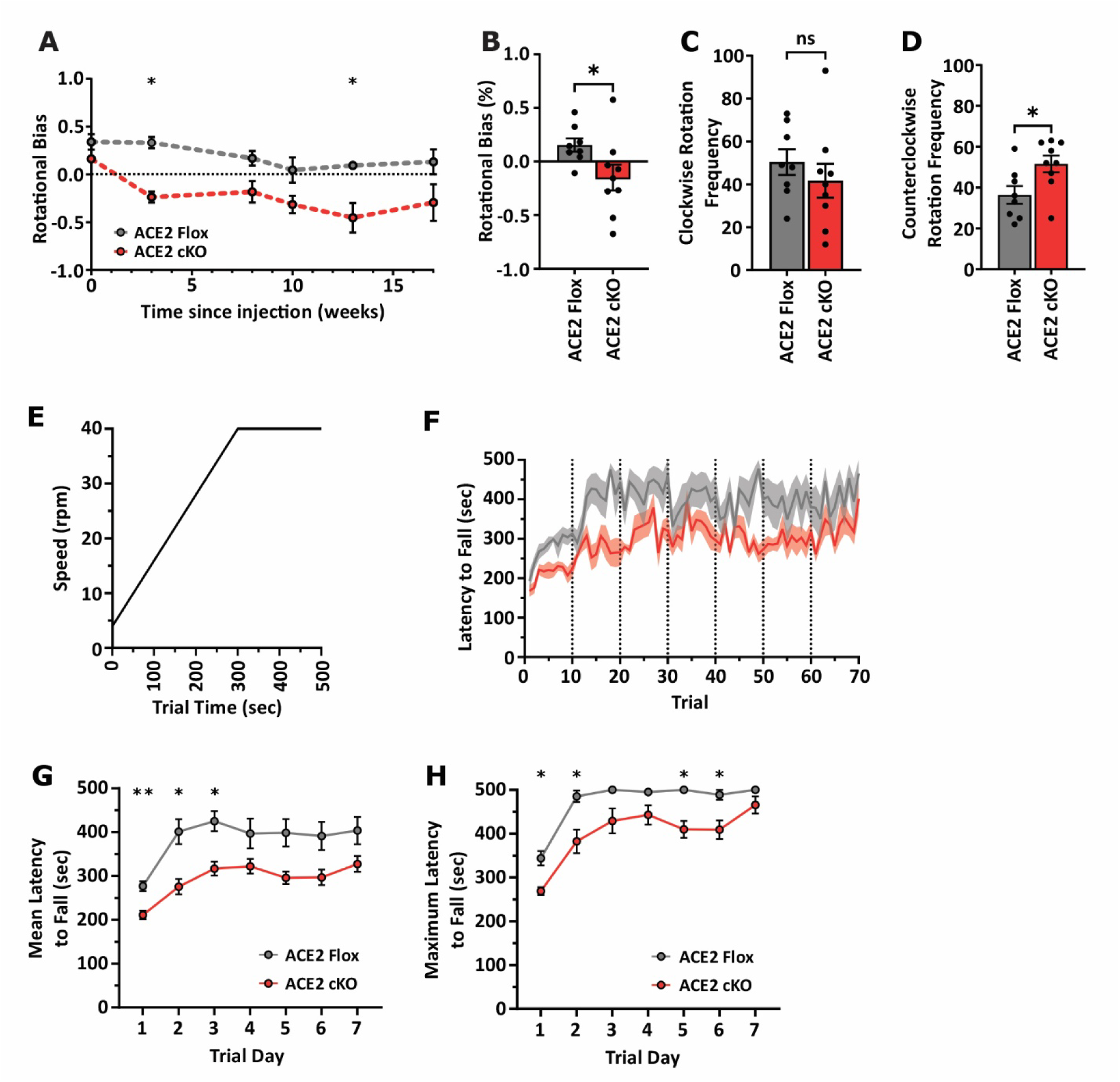
Conditional ACE2 Ablation in the DLS Induces Ipsilateral Rotational Bias and Impairs Motor Learning. (A–D) Rotational motor output was measured in fiber photometry–implanted ACE2 Flox (grey; n = 8) and ACE2 cKO (red; n = 9) mice 14 weeks after unilateral AAV injections. Data are presented as mean ± SEM. (A) Longitudinal analysis of rotational bias was performed in ACE2 Flox (n = 3) and ACE2 cKO (n = 3) mice at multiple time points following unilateral AAV injections. Two-way repeated measures ANOVA revealed main effects of treatment (**p = 0.0011) and time (*p = 0.0292), with no interaction (p = 0.6434). Post hoc Holm–Šídák’s test: significant differences at 3 weeks (*p = 0.0138) and 13 weeks (*p = 0.0160), but not at other time points. (B) Rotational bias ratio (unpaired two-tailed t-test: *p = 0.0485). (C) Clockwise rotation frequency (unpaired two-tailed t-test: p = 0.4034). (D) Counterclockwise rotation frequency (unpaired two-tailed t-test: *p = 0.0228). (E) Representative acceleration profile of the rotarod (4–40 rpm over 300 s, followed by 40 rpm for 200 s). **(F–H)** Latency to fall in ACE2 Flox (grey; n = 6) and ACE2 cKO (red; n = 9) mice, six weeks post-bilateral AAV injections. **(F)** Performance across ten trials per day over seven consecutive days. **(G)** Mean latency to fall across training days. Two-way repeated measures ANOVA with Geisser–Greenhouse correction revealed main effects of treatment (***p = 0.0007) and time (****p < 0.0001), with no interaction (p = 0.5403). Holm–Šídák’s test: significant differences on days 1 (**p = 0.0059), 2 (*p = 0.0230), and 3 (*p = 0.0201), but not on later days. **(H)** Maximum latency to fall across training days in ACE2 Flox (n = 6) and ACE2 cKO (n = 9) mice. Mixed-effects model (REML) with Geisser–Greenhouse correction revealed main effects of treatment (***p = 0.0006) and time (****p < 0.0001), with no interaction (p = 0.4741). Holm–Šídák’s test: significant differences on days 1 (*p = 0.0224), 2 (*p = 0.0259), 5 (*p = 0.0111), and 6 (*p = 0.0259), but not on other days. Data represent pooled results from two independent experimental cohorts with consistent findings. See <u>Table S6</u> for full statistical details.

To see if these deficits in natural locomotion applied to more regimented motor learning tasks, we investigated whether ACE2 ablation impairs motor learning in a rotarod task. Adult ACE2 floxed mice were bilaterally injected with either AAV-eGFP (controls) or AAV-Cre-eGFP (cKO). Six weeks after viral delivery, motor learning was assessed using an accelerating rotarod task (Fig. 6E), with mice completing 10 trials per day over 7 consecutive days (Fig. 6F). ACE2 cKO mice demonstrated a significant impairment in learning the motor task, as indicated by reduced mean latency to fall during the first three trial days (Fig. 6G). In addition, maximum performance across trials was significantly lower in ACE2 cKO mice compared to Flox controls on days 1, 2, 5, and 6 (Fig. 6G-H). Notably, while both groups showed improvements over time, the ACE2 cKO groups consistently underperformed, suggesting a deficit in both early task acquisition and peak performance of the learned motor behavior.

Together, these findings suggest that local ACE2 activity in the DLS is required for normal directional motor behavior and motor learning.

## Discussion

Previous studies have implicated the RAS in Parkinson’s disease (PD) ^49–55^ and levodopa-induced dyskinesia. ^56–58^ However, whether or not there is physiologically relevant expression of the mono-carboxypeptidase ACE2 in the basal ganglia, a key enzyme for the production of many RAS-related peptidergic neuromodulators and physiological cell stress inhibitors, has not been previously examined (Fig. 1A, Table 1). Our study faithfully verified the previously described expression of RAS components in the basal ganglia and extended this work to CINs, an interneuron population in the striatum critical for both normal function and central to the pathogenesis of PD and LID.^10,11,14^ In addition, we found that conditional, spatially restricted genetic ablation of ACE2 affected the temporal coordination of the neuro-modulators dopamine and acetylcholine, as well as negatively impacted motor learning. These findings are important for several reasons: First, our results suggest that locally produced ACE2 in the striatum serves a physiological role that cannot be substituted by enzymatically active and diffusible ACE2 known to be shed from expressing cells outside of the basal ganglia.^59^ Second, our results support that allosteric, possibly bidirectional, regulation of striatal ACE2 could affect the production, processing and thus balance of multiple, known neuromodulators in the striatum, some of which are known to have opposing physiological effects (Table 1). Third, our results suggest that alterations to striatal ACE2 activity by SARS-CoV-2 spike protein, which is shown to allosterically modulate ACE2 activity^60^ and to persist in the brain of patients following clearance in the periphery^61,62^, could contribute to neurological post-acute sequelae of COVID-19 (nPASC).^63^

One of the most surprising aspects of the phenotypes caused by the ablation or inhibition of ACE2 in the dorsolateral striatum was the specific attenuation of cholinergic “dips” associated with DA bursts. These dips have been linked to periods of hyper-polarizations in CINs that follow elevated burst activity brought on by glutamatergic excitations before returning to tonic baseline activity.^43^ Further, ACE2 ablation specifically impacted the magnitude of the dips but not their duration. This is interesting given the prior observations that magnitude of fluorescent GRAB sensor events can represent the synchrony of underlying neuronal populations.^13^ Since the amplitude of ACh bursts and their frequency is not altered by ACE2 activity loss, excitation of CINs is likely less affected than the D2R-dependent temporal coordination of ACh release among CINs.^22,23^ Consistent with this reasoning, we find that ACE2 loss strongly erodes the ability of DA dynamics to predict ACh changes: A DA burst became less predictive of an ACh dip (Fig. 2M) and a DA dip was less likely to predict an ACh burst (Fig. 3H). These findings suggest that loss of ACE2 enzymatic activity reduced CIN sensitivity to DA, likely through D2R as this receptor is known to bidirectional modulate DA’s inhibitory effect on CINs.^22,23,64^

The anticorrelated dynamics of DA and ACh is critical for reinforcement learning. More specifically, the ACh dip defines a permissive window during which DA can induce Hebbian plasticity at corticostriatal synapses on striatal output neurons in the DLS^14^. Concordant, the reduction in ACh Dip amplitude caused by ACE2 ablation diminishes motor learning (Fig. 6F-H). Our experiments phenocopy the ablation of D2R from CIN, which results in reduced ACh dip amplitude and motor learning.^22^ Thus, these finding’s overlap supports a mechanism wherein ACE2 activity impinges on ACh dynamics during learning through AT1R-D2R cross regulation.

In accordance with this scenario, it has been found previously that AT1R and D2R form functional heteromers capable of bidirectionally modulation.^23,46^ We extended this observation to CIN in two ways: First, our proximity ligation assays support the possibility that AT1R and D2R can form heteromers also on CINs in the DLS (Fig. 5C-D). Second, the pharmacological blockade of AT1R increases the D2R dependent inhibition of CIN activity (Fig. 5A-B) consistent with the possibility that AT1R exerts an inhibitory influence on D2R activity in CIN. AT1R–D2R heteromers appear to form constitutively, as pharmacological inhibition of either receptor does not reduce heteromerization.^46^ However, the activity state of one receptor can modulate the signaling efficacy of the other.^46^ Thus, when ACE2 activity is reduced, local accumulation of Ang II may enhance AT1R signaling within these heteromers, thereby dampening D2R-mediated responses, and reducing CIN sensitivity to midbrain DA.

It is important to note that our ACE2 manipulations will likely also impact other neuropeptide families that are processed by ACE2, including apelins, kinins, dynorphins, neurotensins, ghrelins, and opioids.^25–27^ ^65^ ^66^. Further experiments are required to tease out the role of angiotensin peptide signaling as opposed to other neuropeptide signaling effects in CIN.

Previous work during the COVID-19 pandemic explored the neuro-invasive potential of SARS-CoV-2 in the CNS, including the basal ganglia^67–69^, with evidence of viral particle presence in neurons of post mortem COVID patients.^61,62^ Intraneuronal virus in the basal ganglia supported indirectly the presence of membrane-bound ACE2, the cellular receptor of SARS-CoV-2. Our data reveals that physiologically relevant ACE2 is locally expressed in the DLS. Disruption of this ACE2 function in the DLS in response to SARS-Cov2 exposure ^70^ may contribute to long-term motor and cognitive symptoms observed in neurological post-acute sequalae of COVID-19 (nPASC).

## Figures

**Figure S1.**
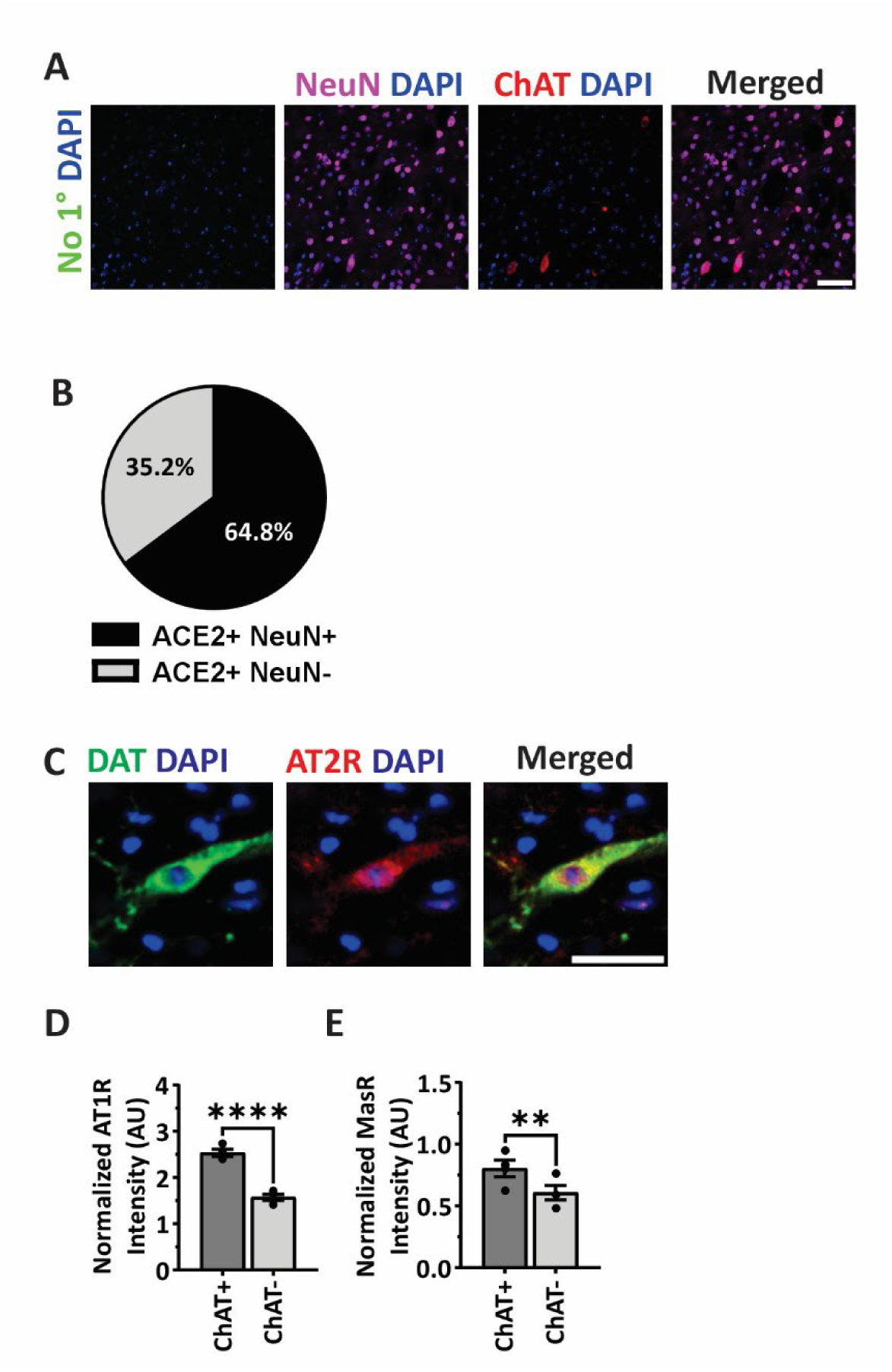
Validation of RAS Antibody Specificity and Receptor Expression in Cholinergic Neurons of the Dorsolateral Striatum (Related to Figure 1) (**A**) Negative control immunostaining in dorsolateral striatum (DLS) sections from an adult wild-type (WT) mouse. Sections were co-stained with NeuN (magenta), ChAT (red), and DAPI (blue) without primary antibody (green). Minimal background fluorescence was observed, confirming the specificity of RAS antibody staining shown in Figure 1. Scale bar: 50 µm. **(B)** Quantification of the proportion of ACE2+ ROIs that were NeuN+ (black) and NeuN-(grey). **(C)** Validation of AT2R antibody by immunostaining a cell type with confirmed AT2R expression. Representative confocal image demonstrates the expected expression pattern. Scale bar: 50 µm. **(D)** AT1R expression in ChAT+ NeuN+ vs. ChAT– NeuN+ ROIs was quantified across multiple animals (n = 4 adult WT males). Data are presented as mean ± SEM. Paired two-tailed t-test: ****p < 0.0001. **(E)** MasR expression in ChAT+ NeuN+ vs. ChAT– NeuN+ ROIs was quantified across multiple animals (n = 4 adult WT males). Data are presented as mean ± SEM. Paired two-tailed t-test: **p = 0.0027. See <u>Table S7</u> for full statistical details.

**Figure S2.**
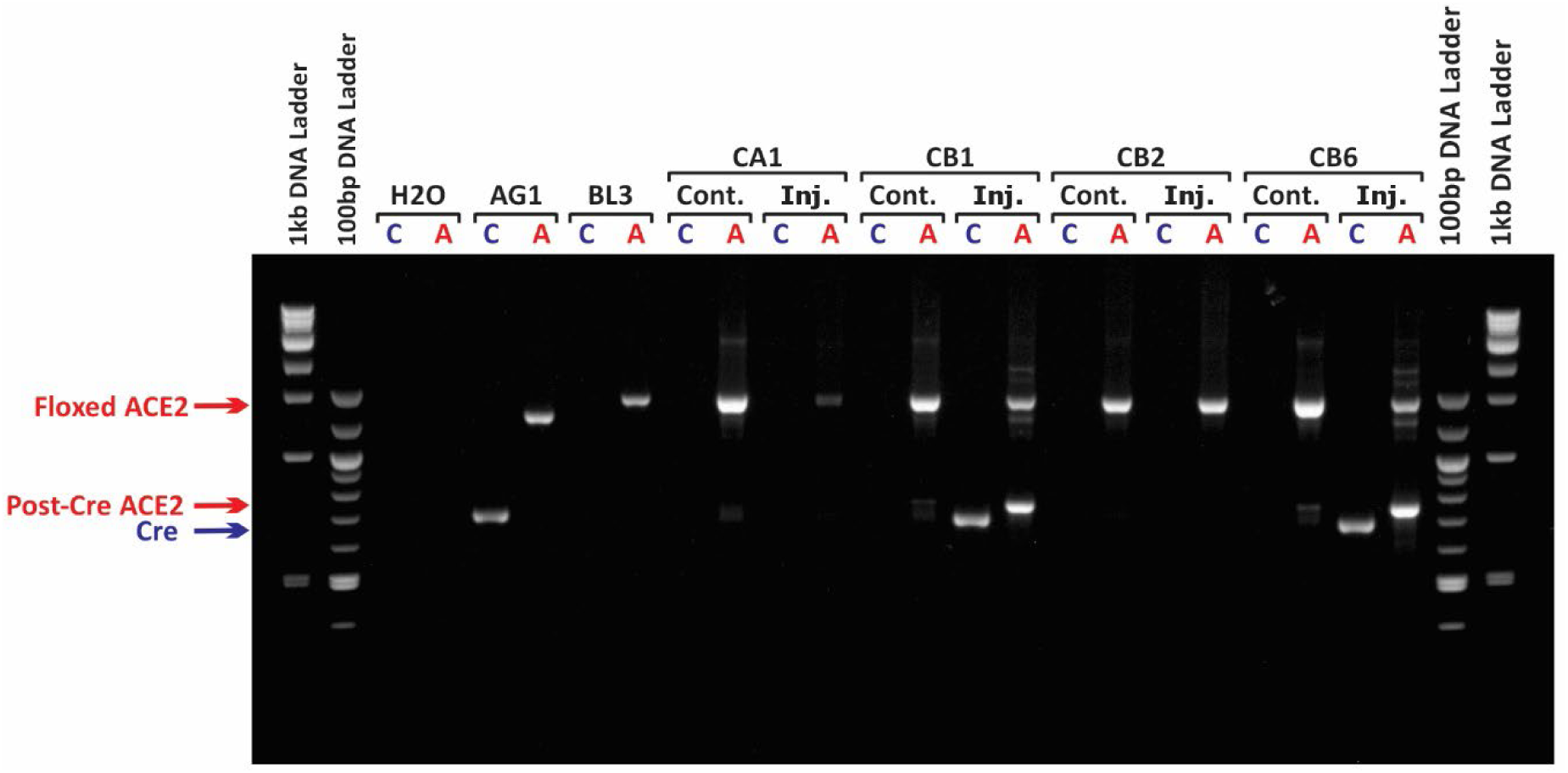
PCR Confirmation of ACE2 cKO (Related to Figure 2-4) A representative PCR gel confirming ACE2 ablation in conditional knockout (cKO) mice. Tissue punches were collected from the dorsolateral striatum (DLS) of both hemispheres – one at the viral injection site (Inj.) and the other from the contralateral control region (Cont.) in the uninjected hemisphere. DNA was extracted and qualitative PCR was performed to detect Cre recombinase and excision of exon 4 in ACE2 conditional knockout (cKO) mice. In ACE2 Flox controls, the intact ACE2 locus was detected, while cKO animals showed successful exon 4 excision. Water (H₂O) served as a no DNA template negative control. AG1 was used as a Cre-positive control; BL3 served as a floxed ACE2-positive control. “C” denotes PCR with Cre primers; “A” denotes PCR with ACE2 primers. NEB 1 kb and Quick-Load 100 bp DNA ladders were used as size markers. Mice CA1 and CB2 were ACE2 Flox controls; CB1 and CB6 were ACE2 cKO.

**Figure S3.**
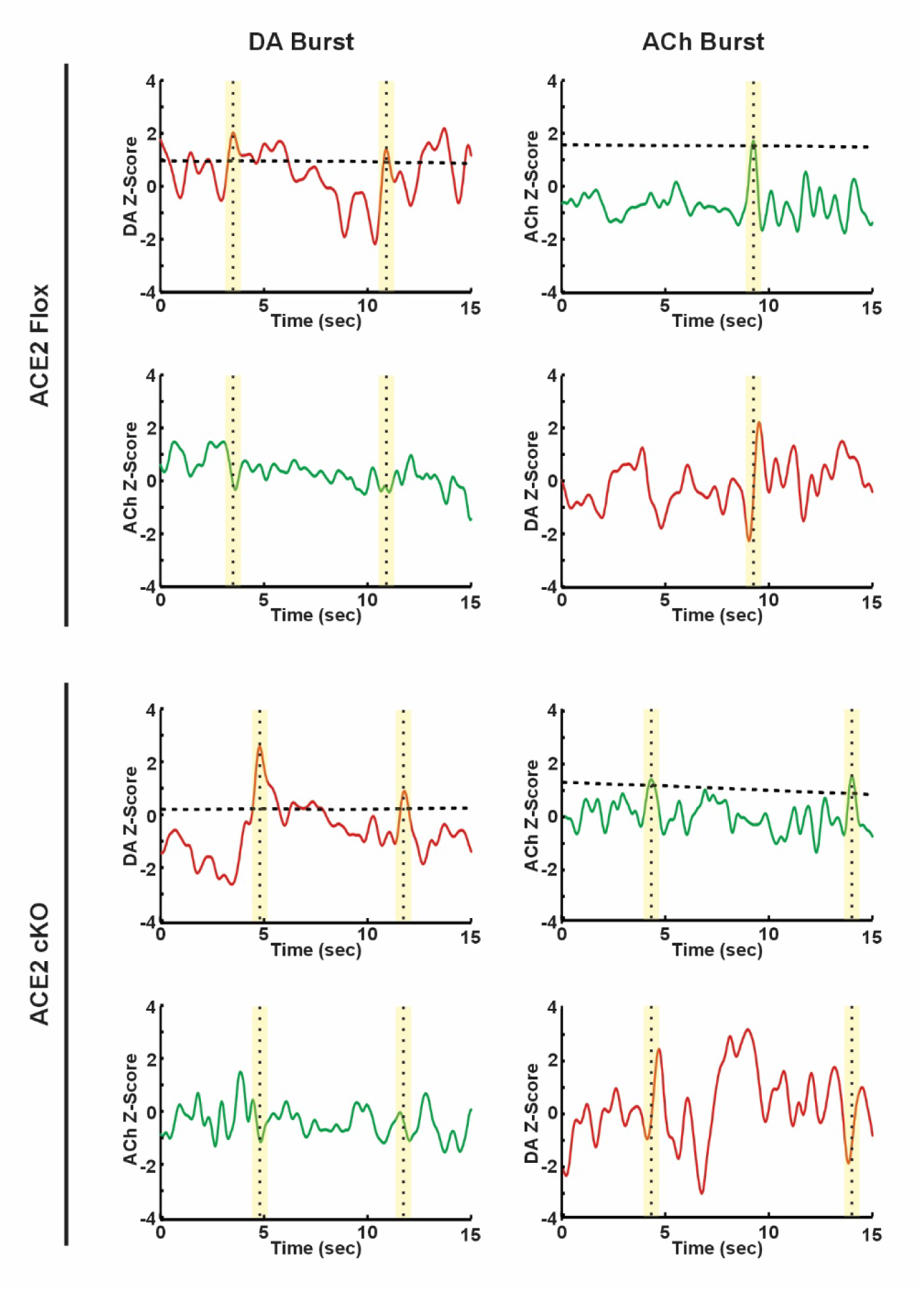
Event Detection Methodology for GRAB-ACh and GRAB-rDA Signals (Related to Figure 2-4) Representative examples of event detection and averaging methodology for dopamine (DA, red traces) and acetylcholine (ACh, green traces) signals in ACE2 Flox (top two rows) and ACE2 cKO (bottom two rows) mice isolating DA bursts (first column) and ACh bursts (second column). Events (vertical dashed lines) were identified as local maxima exceeding 3 median absolute deviations (MAD) above a moving median baseline (horizontal dashed line). For each detected event, 3-s windows centered on the event were extracted from both the guiding channel and the corresponding follow channel.

**Figure S4.**
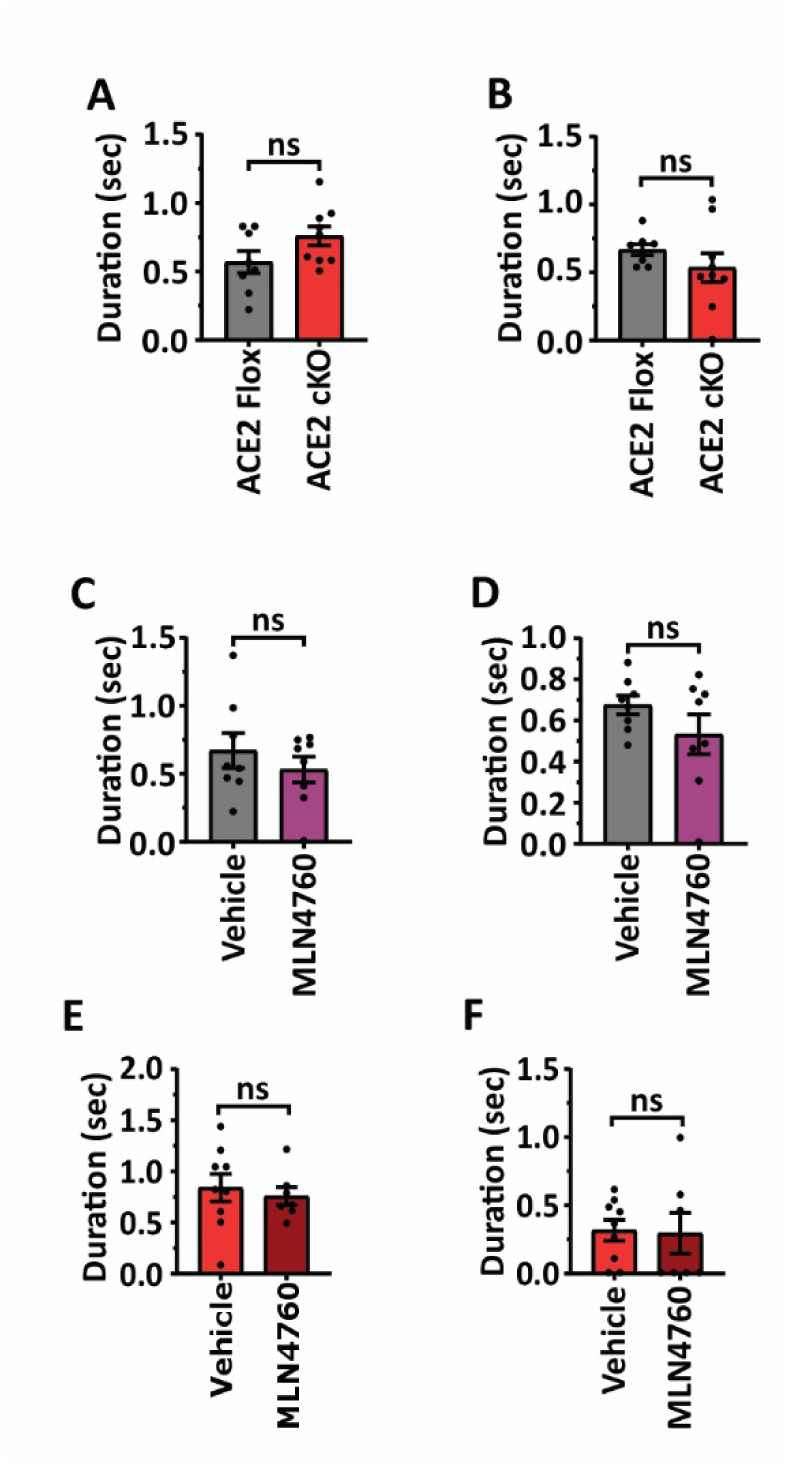
ACE2 ablation or inhibition does not alter ACh peak or dip durations during DA burst events (Related to Figure 2-4) **(A–B)** ACh traces during DA burst events were compared between ACE2 Flox control mice (grey; n = 8) and ACE2 cKO mice (red; n = 9). Data are presented as mean ± SEM. **(A)** Duration of the ACh peak preceding the DA burst (unpaired two-tailed t-test: p = 0.0936). **(B)** Duration of the ACh dip following the DA burst (Welch’s two-tailed t-test: p = 0.2724). **(C–D)** ACh traces during DA burst events were compared within ACE2 Flox control mice (n = 8) following local injections of vehicle (grey) vs. 0.1 mM MLN4760 (purple). **(C)** Duration of the ACh peak preceding the DA burst (paired two-tailed t-test: p = 0.4668). **(D)** Duration of the ACh dip following the DA burst (paired two-tailed t-test: p = 0.2524). **(E–F)** ACh traces during DA burst events were compared within ACE2 cKO mice (n = 7) following local injections of vehicle (red) vs. 0.1 mM MLN4760 (dark red). **(E)** Duration of the ACh peak preceding the DA burst (paired two-tailed t-test: p = 0.9998). **(F)** Duration of the ACh dip following the DA burst (paired two-tailed t-test: p = 0.5051). Data represent pooled results from two independent experimental cohorts with consistent findings. See Table S8 for full statistical details.

**Figure S5.**
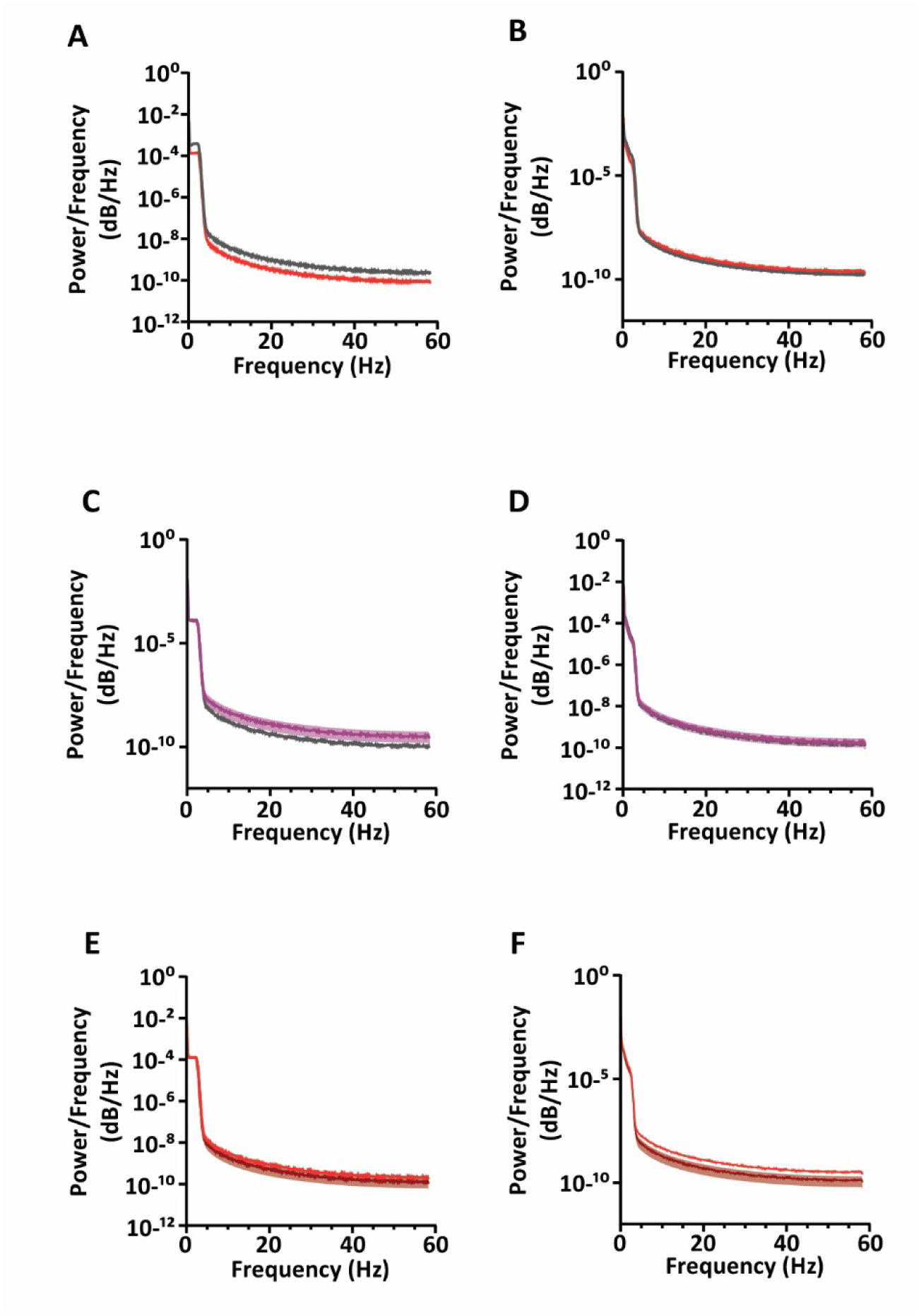
Power Spectral Density (PSD) Analysis of GRAB-ACh and GRAB-rDA Signals. (Related to Figure 2 and 3) (A-B) Power spectral density of GRAB-ACh **(A)** and GRAB-rDA **(B)** traces from ACE2 Flox (grey; n = 8) and ACE2 cKO (red; n = 9) mice. **(C-D)** Power spectral density of GRAB-ACh **(C)** and GRAB-rDA **(D)** traces from ACE2 Flox mice treated with vehicle (grey; n = 8) or 0.1 mM MLN4760 (purple; n = 7). **(E-F)** Power spectral density of GRAB-ACh **(E)** and GRAB-rDA **(F)** traces from ACE2 cKO mice treated with vehicle (red; n = 9) or 0.1 mM MLN4760 (dark red; n = 7).

**Figure S6.**
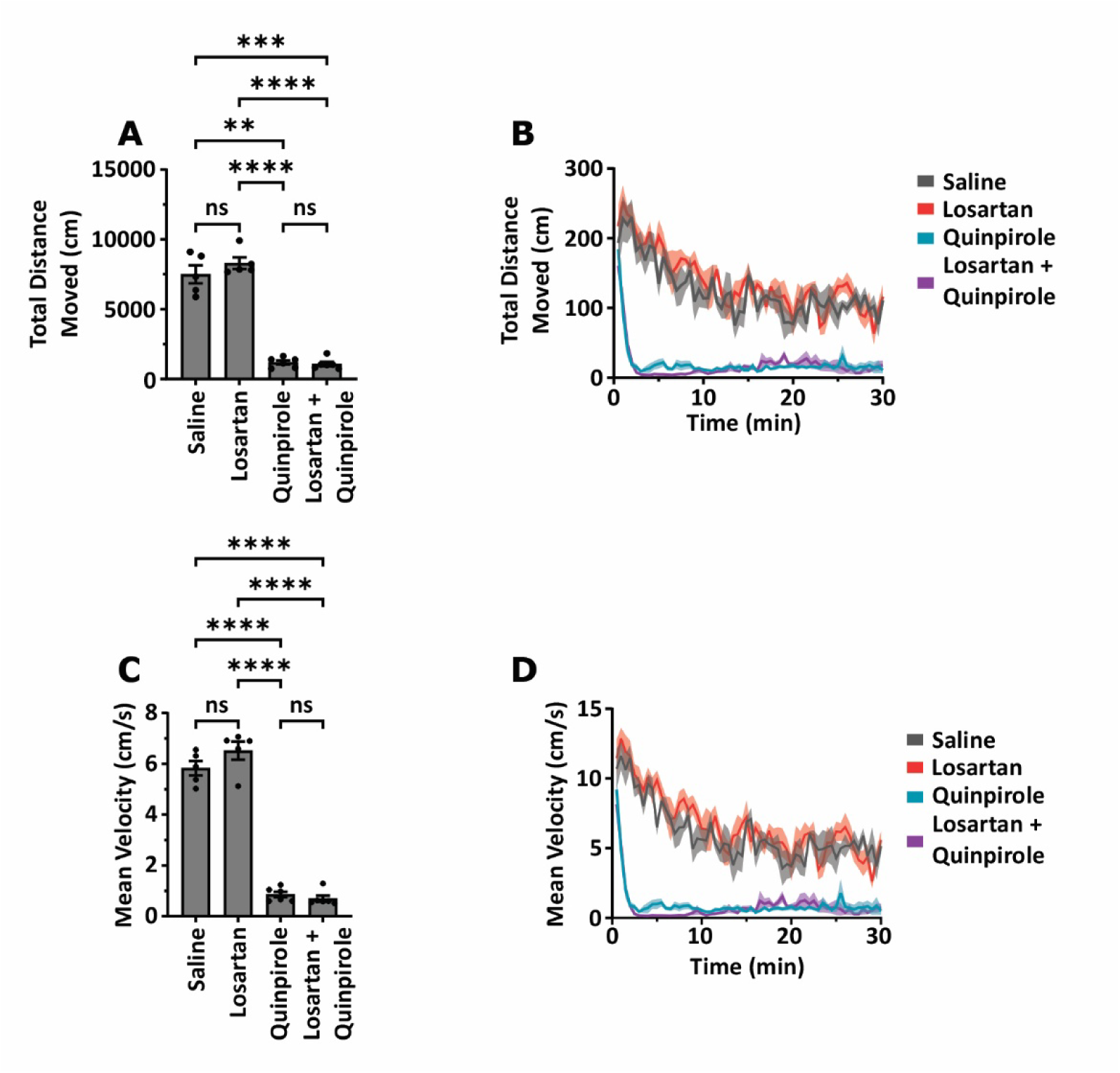
D2R Agonist Quinpirole Reduces Motor Output with or without AT1R Inhibition in Open Field Behavior (Related to Figure 4) (**A–B**) Total distance moved in the open field was measured for 30 minutes immediately following injection with saline (grey; n = 5), 5 mg/kg losartan (red; n = 5), 3 mg/kg quinpirole (blue; n = 6), or losartan + quinpirole (purple; n = 6). Data are presented as mean ± SEM. **(A)** Brown–Forsythe ANOVA (****p < 0.0001) and Welch’s ANOVA (****p < 0.0001). Dunnett’s T3 post hoc comparisons are reported in Table S9. **(C–D)** Mean velocity was measured in the same mice across treatment groups. Data are presented as mean ± SEM. **(C.)** One-way ANOVA (****p < 0.0001). Holm–Šídák’s post hoc comparisons are reported in Table S9. Data represent pooled results from two independent experimental cohorts with consistent findings. See Table S9 for full statistical details.

**Figure S7.**
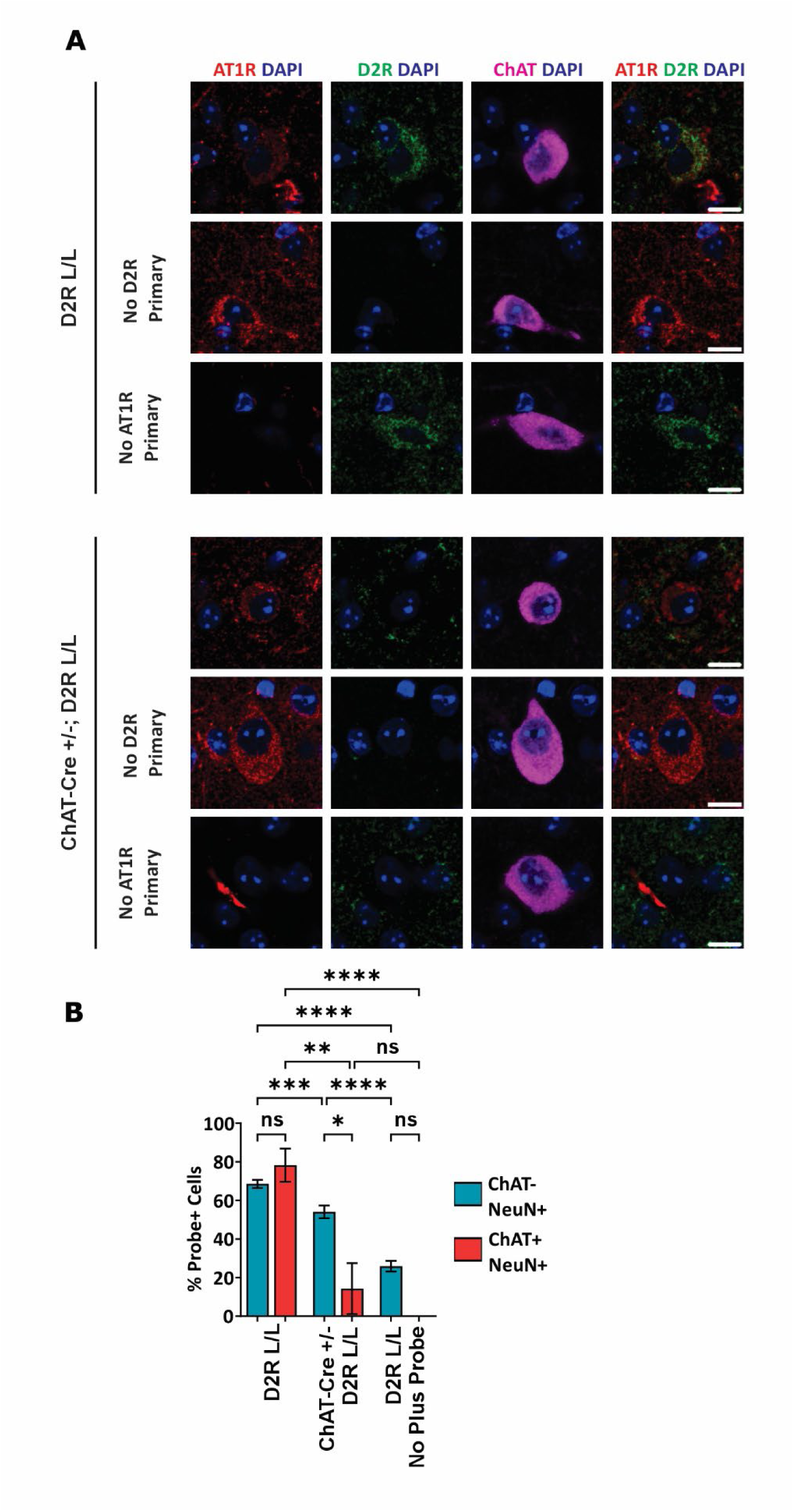
Antibody Validation and Detection of AT1R–D2R Heterodimers in the DLS (Related to Figure 4). (**A**) Representative confocal microscopy images demonstrating specificity of the mouse anti-AT1R and rabbit anti-D2R primary antibodies used for the proximity ligation assay (PLA). AT1R (red), D2R (green), ChAT (magenta), and DAPI (blue) staining are shown. No-primary antibody controls demonstrate minimal background immunoreactivity. In sections from mice with D2R ablated in ChAT-expressing cells, D2R immunoreactivity was reduced. Scale bars: 10 µm. **(B)** Percentage of DLS ChAT-NeuN+ and ChAT+ NeuN+ ROIs positive for AT1R–D2R dimers across genotypes and probe conditions. Groups include D2R L/L with both plus and minus PLA probes, ChAT-Cre +/-; D2R L/L with both plus and minus PLA probes (biological control), and D2R L/L with minus PLA probe only (technical control). Two-way ANOVA revealed significant effects of genotype (****p < 0.0001), cell type (*p = 0.0275), and interaction (*p = 0.0209). Tukey’s multiple comparisons test indicated significant differences between groups. See Table S10 for full statistical details.

**Figure S8.**
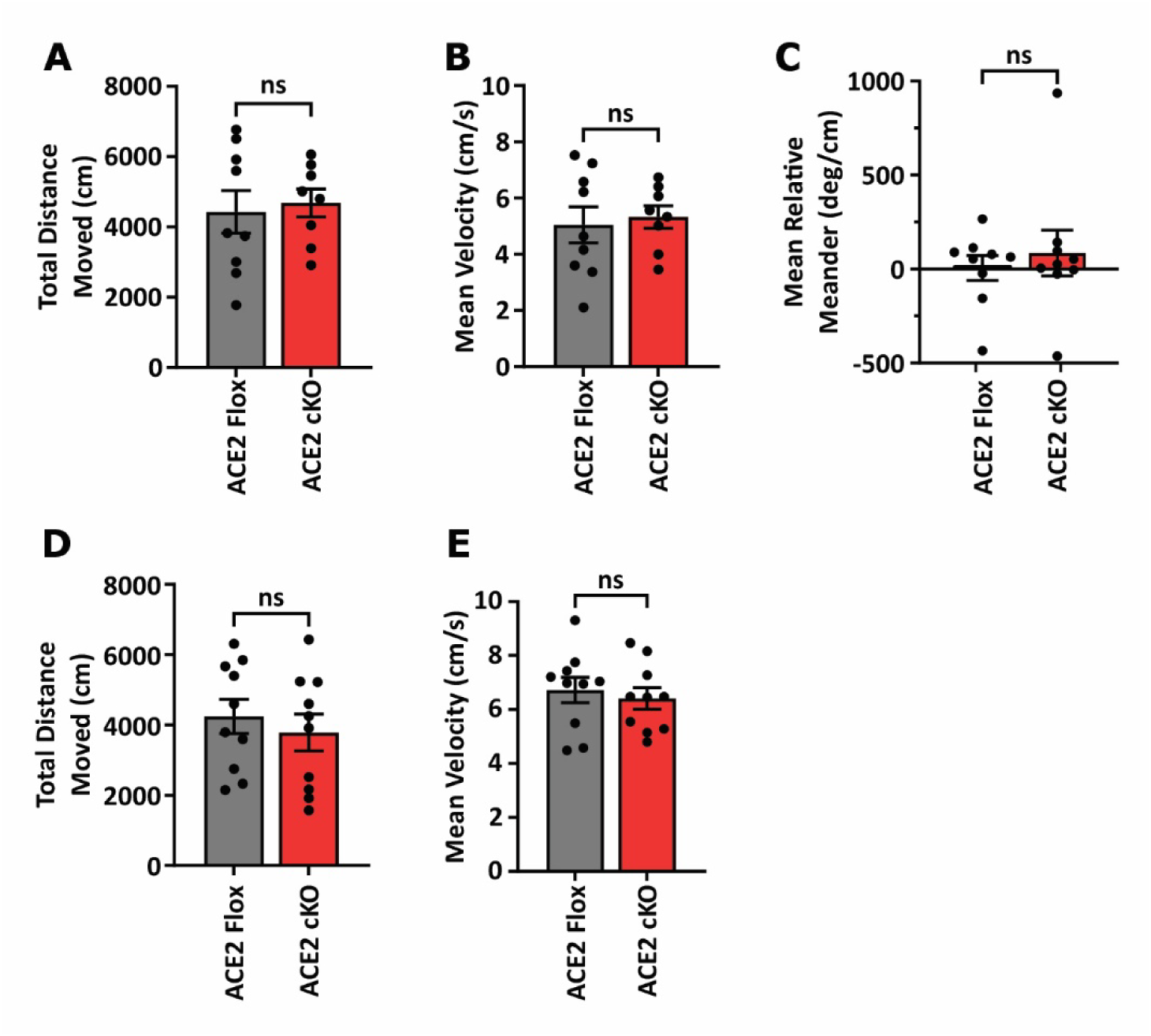
ACE2 Conditional Knockout in the DLS Does Not Impair Motor Output (Related to Figure 5). (**A–C**) Total distance moved, mean velocity, and mean relative meander in fiber photometry–implanted ACE2 Flox (grey; n = 8) and ACE2 cKO (red; n = 9) mice, 14 weeks after unilateral AAV injection. Data are presented as mean ± SEM. **(A)** Total distance moved (unpaired two-tailed t-test: p = 0.7368). **(B)** Mean velocity (unpaired two-tailed t-test: p = 0.7264). **(C)** Mean relative meander (Mann–Whitney U test: p = 0.8633). **(D–E)** Total distance moved and mean velocity in ACE2 Flox (grey; n = 10) and ACE2 cKO (red; n = 10) mice, 4 weeks after bilateral AAV injection. **(D)** Total distance moved (unpaired two-tailed t-test: p = 0.5264). **(E)** Mean velocity (unpaired two-tailed t-test: p = 0.6122). Data represent pooled results from two independent experimental cohorts with consistent findings. See Table S11 for full statistical details.

## Supplemental Tables

**Supplementary Table S1.** (associated with Figure 1). Statistical Table.

|  |  |  |  |  |  |  |  |
| --- | --- | --- | --- | --- | --- | --- | --- |
| **Fig 1C - Mean AT1R Intensity in NeuN+ ROIs vs Negative Control ROIs** |  |  |  |  |  |  |  |
|  | NeuN+ ROIs |  |  | Neg. Cont. ROIs |  |  |  |
|  | Mean | SEM | n | Mean | SEM | n |  |
| Fluorescence Intensity (AU) | 83.86 | 0.5694 | 3857 | 30.47 | 3.068 | 30 | **** |
| Two-tailed Mann-Whitney U test: U = 8732, **** $p < 0.0001$ | | | | | | | |
| **Fig 1D - Mean AT2R Intensity in NeuN+ ROIs vs Negative Control ROIs** |  |  |  |  |  |  |  |
|  | NeuN+ ROIs |  |  | Neg. Cont. ROIs |  |  |  |
|  | Mean | SEM | n | Mean | SEM | n |  |
| Fluorescence Intensity (AU) | 33.24 | 1.259 | 507 | 27.01 | 4.490 | 25 | ns |
| Two-tailed Mann-Whitney U Test: U = 5234, $p = 0.1422$ | | | | | | | |
| **Fig 1E - Mean MasR Intensity in NeuN+ ROIs vs Negative Control ROIs** |  |  |  |  |  |  |  |
|  | NeuN+ ROIs |  |  | Neg. Cont. ROIs |  |  |  |
|  | Mean | SEM | n | Mean | SEM | n |  |
| Fluorescence Intensity (AU) | 37.06 | 0.5335 | 2456 | 12.63 | 1.500 | 30 | **** |
| Two-tailed Mann-Whitney U Test: U = 13694, **** $p < 0.0001$ | | | | | | | |
| **Fig 1F - Mean AT4R Intensity in NeuN+ ROIs vs Negative Control ROIs** |  |  |  |  |  |  |  |
|  | NeuN+ ROIs |  |  | Neg. Cont. ROIs |  |  |  |
|  | Mean | SEM | n | Mean | SEM | n |  |
| Fluorescence Intensity (AU) | 40.29 | 0.3841 | 2491 | 10.00 | 1.205 | 25 | **** |
| Two-tailed Mann-Whitney U Test: U = 2750, **** $p < 0.0001$ | | | | | | | |
| **Fig 1G - Mean MrgDR Intensity in NeuN+ ROIs vs Negative Control ROIs** |  |  |  |  |  |  |  |
|  | NeuN+ ROIs |  |  | Neg. Cont. ROIs |  |  |  |
|  | Mean | SEM | n | Mean | SEM | n |  |
| Fluorescence Intensity (AU) | 56.79 | 0.4565 | 2721 | 40.07 | 3.211 | 30 | **** |
| Two-tailed Mann-Whitney U Test: U = 22017, **** $p < 0.0001$ | | | | | | | |
| **Fig 1H - Mean AT1R Intensity in ChAT+ NeuN+ vs ChAT- NeuN+ ROIs** |  |  |  |  |  |  |  |
|  | ChAT+ NeuN+ ROIs |  |  | ChAT- NeuN+ ROIs |  |  |  |
|  | Mean | SEM | n | Mean | SEM | n |  |
| Fluorescence Intensity (NeuN-normalized AU) | 1.427 | 0.03667 | 68 | 1.055 | 0.006938 | 3789 | **** |
| Two-tailed Mann-Whitney U Test: U = 50026, **** $p < 0.0001$ | | | | | | | |
| **Fig 1I - Mean MasR Intensity in ChAT+ NeuN+ vs ChAT- NeuN+ ROIs** |  |  |  |  |  |  |  |
|  | ChAT+ NeuN+ ROIs |  |  | ChAT- NeuN+ ROIs |  |  |  |
|  | Mean | SEM | n | Mean | SEM | n |  |
| Fluorescence Intensity (NeuN-normalized AU) | 1.866 | 0.1285 | 34 | 0.7837 | 0.01009 | 2422 | **** |
| Two-tailed Mann-Whitney U Test: U = 7256, **** $p < 0.0001$ | | | | | | | |
| **Fig 1J - Mean AT4R Intensity in ChAT+ NeuN+ vs ChAT- NeuN+ ROIs** |  |  |  |  |  |  |  |
|  | ChAT+ NeuN+ ROIs |  |  | ChAT- NeuN+ ROIs |  |  |  |
|  | Mean | SEM | n | Mean | SEM | n |  |
| Fluorescence Intensity (NeuN-normalized AU) | 1.270 | 0.06740 | 41 | 0.8537 | 0.008239 | 2450 | **** |
| Two-tailed Mann-Whitney U Test: U = 22689, **** $p < 0.0001$ | | | | | | | |
| **Fig 1K - Mean MrgDR Intensity in ChAT+ NeuN+ vs ChAT- NeuN+ ROIs** |  |  |  |  |  |  |  |
|  | ChAT+ NeuN+ ROIs |  |  | ChAT- NeuN+ ROIs |  |  |  |
|  | Mean | SEM | n | Mean | SEM | n |  |
| Fluorescence Intensity (NeuN-normalized AU) | 1.834 | 0.1130 | 33 | 1.658 | 0.01243 | 2688 | ns |
| Two-tailed Mann-Whitney U Test: U = 37003, $p = 0.1015$ | | | | | | | |
| **Fig 1L - Percentage of AT1R-positive ChAT+ NeuN+ vs ChAT- NeuN+ ROIs** |  |  |  |  |  |  |  |
|  | ChAT+ NeuN+ ROIs |  |  | ChAT- NeuN+ ROIs |  |  |  |
|  | Mean | SEM | n | Mean | SEM | n |  |
| Percentage ROIs positive for AT1R | 1.000 | 0.000 | 68 | 0.9301 | 0.004144 | 3789 | * |
| Two-tailed Mann-Whitney U Test: U = 119816, * $p = 0.0241$ | | | | | | | |
| **Fig 1M - Percentage of MasR-positive ChAT+ NeuN+ vs ChAT- NeuN+ ROIs** |  |  |  |  |  |  |  |
|  | ChAT+ NeuN+ ROIs |  |  | ChAT- NeuN+ ROIs |  |  |  |
|  | Mean | SEM | n | Mean | SEM | n |  |
| Percentage ROIs positive for MasR | 1.000 | 0.000 | 34 | 0.8485 | 0.007287 | 2422 | * |
| Two-tailed Mann-Whitney U Test: U = 34935, * $p = 0.0119$ | | | | | | | |
| **Fig 1N - Percentage of AT4R-positive ChAT+ NeuN+ vs ChAT- NeuN+ ROIs** |  |  |  |  |  |  |  |
|  | ChAT+ NeuN+ ROIs |  |  | ChAT- NeuN+ ROIs |  |  |  |
|  | Mean | SEM | n | Mean | SEM | n |  |
| Percentage ROIs positive for AT4R | 0.9756 | 0.02439 | 41 | 0.9469 | 0.004530 | 2450 | ns |
| Two-tailed Mann-Whitney U Test: $U = 48785$, $p = 0.5214$ | | | | | | | |
| **Fig 1O - Percentage of MrgDR-positive ChAT+ NeuN+ vs ChAT- NeuN+ ROIs** |  |  |  |  |  |  |  |
|  | ChAT+ NeuN+ ROIs |  |  | ChAT- NeuN+ ROIs |  |  |  |
|  | Mean | SEM | n | Mean | SEM | n |  |
| Percentage ROIs positive for MrgDR | 0.9394 | 0.04218 | 33 | 0.7779 | 0.008019 | 2688 | * |
| Two-tailed Mann-Whitney U Test: $U = 37190$, * $p = 0.0315$ | | | | | | | |

**Supplementary Table S2.** (associated with Figure 2). Statistical Table.

|  |  |  |  |  |  |  |  |
| --- | --- | --- | --- | --- | --- | --- | --- |
| **Fig 2F - Mean DA burst amp. in ACE2 Flox vs. ACE2 cKO mice** |  |  |  |  |  |  |  |
|  | ACE2 Flox mice |  |  | ACE2 cKO mice |  |  |  |
|  | Mean | SEM | n | Mean | SEM | n |  |
| Amplitude (Z-score) | 0.8584 | 0.1215 | 8 | 0.8439 | 0.06465 | 9 | ns |
| Unpaired two-tailed t-test: $t(15) = 0.1090$, $p = 0.9147$ | | | | | | | |
| **Fig 2G - Mean DA burst frequency in ACE2 Flox vs. ACE2 cKO mice** |  |  |  |  |  |  |  |
|  | ACE2 Flox mice |  |  | ACE2 cKO mice |  |  |  |
|  | Mean | SEM | n | Mean | SEM | n |  |
| Frequency (events/min) | 6.692 | 0.3463 | 8 | 6.491 | 0.3813 | 9 | ns |
| Unpaired two-tailed t-test: $t(15) = 0.3852$, $p = 0.7055$ | | | | | | | |
| **Fig 2I - Mean pre-DA burst ACh peak amp. in ACE2 Flox vs. ACE2 cKO mice** |  |  |  |  |  |  |  |
|  | ACE2 Flox mice |  |  | ACE2 cKO mice |  |  |  |
|  | Mean | SEM | n | Mean | SEM | n |  |
| Amplitude (Z-score) | 0.1700 | 0.02956 | 8 | 0.1607 | 0.01694 | 9 | ns |
| Unpaired two-tailed t-test: $t(15) = 0.2799$, $p = 0.7834$ | | | | | | | |
| **Fig 2J - Mean post-DA burst ACh dip amp. in ACE2 Flox vs. ACE2 cKO mice** |  |  |  |  |  |  |  |
|  | ACE2 Flox mice |  |  | ACE2 cKO mice |  |  |  |
|  | Mean | SEM | n | Mean | SEM | n |  |
| Amplitude (Z-score) | -0.3773 | 0.06118 | 8 | -0.1083 | 0.03977 | 9 | ** |
| Unpaired two-tailed t-test: $t(15) = 3.769$, ** $p = 0.0019$ | | | | | | | |
| **Fig 2L - Max. correlation between ACh lagged relative to DA burst in ACE2 Flox vs. ACE2 cKO** |  |  |  |  |  |  |  |
|  | ACE2 Flox mice |  |  | ACE2 cKO mice |  |  |  |
|  | Mean | SEM | n | Mean | SEM | n |  |
| Max. correlation (r) | 0.1697 | 0.1110 | 8 | 0.4711 | 0.4711 | 9 | * |
| Two-tailed Mann-Whitney U Test: U = 12, * $p = 0.0206$ | | | | | | | |
| **Fig 2M - Min. correlation between ACh lagged relative to DA burst in ACE2 Flox vs. ACE2 cKO** |  |  |  |  |  |  |  |
|  | ACE2 Flox mice |  |  | ACE2 cKO mice |  |  |  |
|  | Mean | SEM | n | Mean | SEM | n |  |
| Min. correlation (r) | -0.7170 | 0.05216 | 8 | -0.1878 | 0.1027 | 9 | *** |
| Unpaired two-tailed t-test: $t(15) = 4.416$, *** $p = 0.0005$ | | | | | | | |
| **Fig 2O - Mean DA burst amp. in ACE2 Flox mice treated with vehicle vs 0.1mM MLN4760** |  |  |  |  |  |  |  |
|  | ACE2 Flox mice treated with vehicle |  |  | ACE2 Flox mice treated with MLN4760 |  |  |  |
|  | Mean | SEM | n | Mean | SEM | n |  |
| Amplitude (Z-score) | 0.4733 | 0.06805 | 8 | 0.4451 | 0.08697 | 8 | ns |
| Paired two-tailed t-test: $t(7) = 0.2454$, $p = 0.8132$ | | | | | | | |
| **Fig 2P - Mean DA burst frequency in ACE2 Flox mice treated with vehicle vs 0.1mM MLN4760** |  |  |  |  |  |  |  |
|  | ACE2 Flox mice treated with vehicle |  |  | ACE2 Flox mice treated with MLN4760 |  |  |  |
|  | Mean | SEM | n | Mean | SEM | n |  |
| Frequency (events/min) | 5.603 | 0.3700 | 7 | 6.739 | 0.3642 | 7 | ns |
| Paired two-tailed t-test: $t(6) = 2.096$, $p = 0.0809$ | | | | | | | |
| **Fig 2R - Mean pre-DA burst ACh peak amp. in ACE2 Flox mice treated with vehicle vs 0.1mM MLN4760** |  |  |  |  |  |  |  |
|  | ACE2 Flox mice treated with vehicle |  |  | ACE2 Flox mice treated with MLN4760 |  |  |  |
|  | Mean | SEM | n | Mean | SEM | n |  |
| Amplitude (Z-score) | 0.1040 | 0.02415 | 8 | 0.05479 | 0.02140 | 8 | ns |
| Paired two-tailed t-test: $t(7) = 2.245$, $p = 0.0597$ | | | | | | | |
| **Fig 2S - Mean post-DA burst ACh dip amp. in ACE2 Flox mice treated with vehicle vs 0.1mM MLN4760** |  |  |  |  |  |  |  |
|  | ACE2 Flox mice treated with vehicle |  |  | ACE2 Flox mice treated with MLN4760 |  |  |  |
|  | Mean | SEM | n | Mean | SEM | n |  |
| Amplitude (Z-score) | -0.1860 | 0.03715 | 8 | -0.05656 | 0.01974 | 8 | ** |
| Paired two-tailed t-test: $t(7) = 3.750$, ** $p = 0.0072$ | | | | | | | |
| **Fig 2U - Max. correlation between ACh lagged relative to DA burst in ACE2 Flox mice treated with vehicle vs. 0.1mM MLN4760** |  |  |  |  |  |  |  |

|  | ACE2 Flox mice treated with vehicle |  |  | ACE2 Flox mice treated with MLN4760 |  |  |  |
| --- | --- | --- | --- | --- | --- | --- | --- |
|  | Mean | SEM | n | Mean | SEM | n |  |
| Max. correlation (r) | 0.2160 | 0.1352 | 8 | 0.3441 | 0.1298 | 8 | ns |
| Paired two-tailed t-test: $t(7) = 0.6101, p = 0.5611$ | | | | | | | |
| **Fig 2V – Min. correlation between ACh lagged relative to DA burst in ACE2 Flox mice treated with vehicle vs. 0.1mM MLN4760** |  |  |  |  |  |  |  |
|  | ACE2 Flox mice treated with vehicle |  |  | ACE2 Flox mice treated with MLN4760 |  |  |  |
|  | Mean | SEM | n | Mean | SEM | n |  |
| Min. correlation (r) | -0.6116 | 0.09068 | 8 | -0.3282 | 0.2038 | 8 | ns |
| Paired two-tailed t-test: $t(7) = 1.177, p = 0.2776$ | | | | | | | |

**Supplementary Table S3.** (associated with Figure 3). Statistical Table.

| **Fig 3B - Mean ACh burst amp. in ACE2 Flox vs. ACE2 cKO mice** |  |  |  |  |  |  |  |
| --- | --- | --- | --- | --- | --- | --- | --- |
|  | ACE2 Flox mice |  |  | ACE2 cKO mice |  |  |  |
|  | Mean | SEM | n | Mean | SEM | n |  |
| Amplitude (Z-score) | 0.8659 | 0.04878 | 8 | 0.9527 | 0.03279 | 9 | ns |
| Unpaired two-tailed t-test: $t(15) = 1.508, p = 0.1524$ | | | | | | | |
| **Fig 3C - Mean ACh burst frequency in ACE2 Flox vs. ACE2 cKO mice** |  |  |  |  |  |  |  |
|  | ACE2 Flox mice |  |  | ACE2 cKO mice |  |  |  |
|  | Mean | SEM | n | Mean | SEM | n |  |
| Frequency (events/min) | 8.228 | 0.1403 | 8 | 7.775 | 0.3358 | 9 | ns |
| Welch's two-tailed t-test: $t(10.67) = 1.243, p = 0.2404$ | | | | | | | |
| **Fig 3E - Mean pre-ACh burst DA dip amp. in ACE2 Flox vs. ACE2 cKO mice** |  |  |  |  |  |  |  |
|  | ACE2 Flox mice |  |  | ACE2 cKO mice |  |  |  |
|  | Mean | SEM | n | Mean | SEM | n |  |
| Amplitude (Z-score) | -0.2371 | 0.05417 | 8 | -0.06090 | 0.02228 | 9 | * |
| Welch's two-tailed t-test: $t(9.336) = 3.008, * p = 0.0142$ | | | | | | | |
| **Fig 3F - Mean post-ACh burst DA peak amp. in ACE2 Flox vs. ACE2 cKO mice** |  |  |  |  |  |  |  |
|  | ACE2 Flox mice |  |  | ACE2 cKO mice |  |  |  |
|  | Mean | SEM | n | Mean | SEM | n |  |
| Amplitude (Z-score) | 0.06341 | 0.01968 | 8 | 0.05460 | 0.01426 | 9 | ns |
| Unpaired two-tailed t-test: $t(15) = 0.3685, p = 0.7177$ | | | | | | | |
| **Fig 3H - Min. correlation between DA lagged relative to ACh burst in ACE2 Flox vs. ACE2 cKO** |  |  |  |  |  |  |  |
|  | ACE2 Flox mice |  |  | ACE2 cKO mice |  |  |  |
|  | Mean | SEM | n | Mean | SEM | n |  |
| Min. correlation (r) | -0.8186 | 0.03723 | 8 | -0.3823 | 0.08957 | 9 | *** |
| Welch's two-tailed t-test: $t(10.64) = 4.497$, *** $p = 0.0010$ | | | | | | | |
| **Fig 3I - Max. correlation between DA lagged relative to ACh burst in ACE2 Flox vs. ACE2 cKO** |  |  |  |  |  |  |  |
|  | ACE2 Flox mice |  |  | ACE2 cKO mice |  |  |  |
|  | Mean | SEM | n | Mean | SEM | n |  |
| Max. correlation (r) | 0.4221 | 0.03903 | 8 | 0.3500 | 0.09159 | 9 | ns |
| Welch's two-tailed t-test: $t(10.76) = 0.7244$, $p = 0.4221$ | | | | | | | |
| **Fig 3K- Mean ACh burst amp. in ACE2 Flox mice treated with vehicle vs 0.1mM MLN4760** |  |  |  |  |  |  |  |
|  | ACE2 Flox mice treated with vehicle |  |  | ACE2 Flox mice treated with MLN4760 |  |  |  |
|  | Mean | SEM | n | Mean | SEM | n |  |
| Amplitude (Z-score) | 0.7061 | 0.08793 | 7 | 0.4997 | 0.06544 | 7 | ns |
| Paired two-tailed t-test: $t(6) = 1.880$, $p = 0.1091$ | | | | | | | |
| **Fig 3L - Mean ACh burst frequency in ACE2 Flox mice treated with vehicle vs 0.1mM MLN4760** |  |  |  |  |  |  |  |
|  | ACE2 Flox mice treated with vehicle |  |  | ACE2 Flox mice treated with MLN4760 |  |  |  |
|  | Mean | SEM | n | Mean | SEM | n |  |
| Frequency (events/min) | 6.950 | 0.4060 | 7 | 7.543 | 0.2305 | 7 | ns |
| Two-tailed Wilcoxon matched-pairs signed rank test: $W = 8.000$, $p = 0.5781$ | | | | | | | |
| **Fig 3N - Mean pre-ACh burst DA dip amp. in ACE2 Flox mice treated with vehicle vs 0.1mM MLN4760** |  |  |  |  |  |  |  |
|  | ACE2 Flox mice treated with vehicle |  |  | ACE2 Flox mice treated with MLN4760 |  |  |  |
|  | Mean | SEM | n | Mean | SEM | n |  |
| Amplitude (Z-score) | -0.06558 | 0.008153 | 7 | -0.03664 | 0.01492 | 7 | ns |
| Paired two-tailed t-test: $t(6) = 2.181$, $p = 0.0719$ | | | | | | | |
| **Fig 3O - Mean post-ACh burst DA peak amp. in ACE2 Flox mice treated with vehicle vs 0.1mM MLN4760** |  |  |  |  |  |  |  |
|  | ACE2 Flox mice treated with vehicle |  |  | ACE2 Flox mice treated with MLN4760 |  |  |  |
|  | Mean | SEM | n | Mean | SEM | n |  |
| Amplitude (Z-score) | 0.02890 | 0.01020 | 7 | 0.02807 | 0.02520 | 7 | ns |
| Two-tailed Wilcoxon matched-pairs signed rank test: $W = -10$, $p = 0.4688$ | | | | | | | |
| **Fig 3Q - Max. correlation between ACh lagged relative to DA burst in ACE2 Flox vs. ACE2 cKO** |  |  |  |  |  |  |  |
|  | ACE2 Flox mice |  |  | ACE2 cKO mice |  |  |  |
|  | Mean | SEM | n | Mean | SEM | n |  |
| Max. correlation (r) | 0.1697 | 0.1110 | 8 | 0.4711 | 0.4711 | 9 | * |
| Two-tailed Mann-Whitney U Test: $U = 12$, * $p = 0.0206$ | | | | | | | |
| **Fig 3R - Min. correlation between ACh lagged relative to DA burst in ACE2 Flox vs. ACE2 cKO** |  |  |  |  |  |  |  |
|  | ACE2 Flox mice |  |  | ACE2 cKO mice |  |  |  |
|  | Mean | SEM | n | Mean | SEM | n |  |
| Min. correlation (r) | -0.7170 | 0.05216 | 8 | -0.1878 | 0.1027 | 9 | *** |
| Unpaired two-tailed t-test: $t(15) = 4.416$, *** $p = 0.0005$ | | | | | | | |

**Supplementary Table S4.** (associated with Figure 4). Statistical Table.

| Fig 4B - Mean DA burst amp. in ACE2 cKO mice treated with vehicle vs 0.1mM MLN4760 |  |  |  |  |  |  |  |
| --- | --- | --- | --- | --- | --- | --- | --- |
|  | ACE2 cKO mice treated with vehicle |  |  | ACE2 cKO mice treated with MLN4760 |  |  |  |
|  | Mean | SEM | n | Mean | SEM | n |  |
| Amplitude (Z-score) | 0.5260 | 0.1225 | 7 | 0.5744 | 0.1578 | 7 | ns |
| Paired two-tailed t-test: $t(6) = 0.6925, p = 0.5145$ | | | | | | | |
| Fig 4C - Mean DA burst frequency in ACE2 cKO mice treated with vehicle vs 0.1mM MLN4760 |  |  |  |  |  |  |  |
|  | ACE2 cKO mice treated with vehicle |  |  | ACE2 cKO mice treated with MLN4760 |  |  |  |
|  | Mean | SEM | n | Mean | SEM | n |  |
| Frequency (events/minute) | 6.474 | 0.3911 | 7 | 7.022 | 0.1654 | 7 | ns |
| Paired two-tailed t-test: $t(6) = 0.5784, p = 0.5841$ | | | | | | | |
| Fig 4E - Mean pre-DA burst ACh peak amp. in ACE2 cKO mice treated with vehicle vs 0.1mM MLN4760 |  |  |  |  |  |  |  |
|  | ACE2 cKO mice treated with vehicle |  |  | ACE2 cKO mice treated with MLN4760 |  |  |  |
|  | Mean | SEM | n | Mean | SEM | n |  |
| Amplitude (Z-score) | 0.08727 | 0.01877 | 7 | 0.08960 | 0.02926 | 7 | ns |
| Paired two-tailed t-test: $t(6) = 0.04714, p = 0.9639$ | | | | | | | |
| Fig 4F - Mean post-DA burst ACh dip amp. in ACE2 cKO mice treated with vehicle vs 0.1mM MLN4760 |  |  |  |  |  |  |  |
|  | ACE2 cKO mice treated with vehicle |  |  | ACE2 cKO mice treated with MLN4760 |  |  |  |
|  | Mean | SEM | n | Mean | SEM | n |  |
| Amplitude (Z-score) | -0.04033 | 0.03097 | 7 | -0.06133 | 0.06987 | 7 | ns |
| Paired two-tailed t-test: $t(6) = 0.1682, p = 0.8720$ | | | | | | | |

**Fig 4H – Max. correlation between ACh lagged relative to DA burst in ACE2 cKO mice treated with vehicle vs 0.1mM MLN4760**
|  | ACE2 cKO mice treated with vehicle |  |  | ACE2 cKO mice treated with MLN4760 |  |  |  |
| --- | --- | --- | --- | --- | --- | --- | --- |
|  | Mean | SEM | n | Mean | SEM | n |  |
| Max. correlation (r) | 0.7822 | 0.05656 | 7 | 0.6813 | 0.07928 | 7 | ns |
| Paired two-tailed t-test: $t(6) = 0.9453, p = 0.3810$ | | | | | | | |

**Fig 4I – Min. correlation between ACh lagged relative to DA burst in ACE2 cKO mice treated with vehicle vs 0.1mM MLN4760**
|  | ACE2 cKO mice treated with vehicle |  |  | ACE2 cKO mice treated with MLN4760 |  |  |  |
| --- | --- | --- | --- | --- | --- | --- | --- |
|  | Mean | SEM | n | Mean | SEM | n |  |
| Min. correlation (r) | -0.04665 | 0.06238 | 7 | -0.1101 | 0.1648 | 7 | ns |
| Paired two-tailed t-test: $t(6) = 0.5994, p = 0.5708$ | | | | | | | |

**Fig 4K- Mean ACh burst amp. in ACE2 cKO mice treated with vehicle vs 0.1mM MLN4760**
|  | ACE2 cKO mice treated with vehicle |  |  | ACE2 cKO mice treated with MLN4760 |  |  |  |
| --- | --- | --- | --- | --- | --- | --- | --- |
|  | Mean | SEM | n | Mean | SEM | n |  |
| Amplitude (Z-score) | 0.4952 | 0.1163 | 7 | 0.6833 | 0.1714 | 7 | ns |
| Paired two-tailed t-test: $t(6) = 2.015, p = 0.0905$ | | | | | | | |

**Fig 4L - Mean ACh burst frequency in ACE2 cKO mice treated with vehicle vs 0.1mM MLN4760**
|  | ACE2 cKO mice treated with vehicle |  |  | ACE2 cKO mice treated with MLN4760 |  |  |  |
| --- | --- | --- | --- | --- | --- | --- | --- |
|  | Mean | SEM | n | Mean | SEM | n |  |
| Frequency (events/min) | 5.998 | 0.3755 | 7 | 8.102 | 0.1281 | 7 | * |
| Two-tailed Wilcoxon matched-pairs signed rank test: $W = 28.00, * p = 0.0156$ | | | | | | | |

**Fig 4N - Mean pre-ACh burst DA dip amp. in ACE2 cKO mice treated with vehicle vs 0.1mM MLN4760**
|  | ACE2 cKO mice treated with vehicle |  |  | ACE2 cKO mice treated with MLN4760 |  |  |  |
| --- | --- | --- | --- | --- | --- | --- | --- |
|  | Mean | SEM | n | Mean | SEM | n |  |
| Amplitude (Z-score) | -0.03955 | 0.02350 | 7 | -0.04096 | 0.04187 | 7 | ns |
| Paired two-tailed t-test: $t(6) = 0.4923, p = 0.6400$ | | | | | | | |

**Fig 4O - Mean post-ACh burst DA peak amp. in ACE2 cKO mice treated with vehicle vs 0.1mM MLN4760**
|  | ACE2 cKO mice treated with vehicle |  |  | ACE2 cKO mice treated with MLN4760 |  |  |  |
| --- | --- | --- | --- | --- | --- | --- | --- |
|  | Mean | SEM | n | Mean | SEM | n |  |
| Amplitude (Z-score) | 0.04849 | 0.01421 | 7 | 0.06709 | 0.02411 | 7 | ns |
| Paired two-tailed t-test: $t(6) = 0.4477, p = 0.6701$ | | | | | | | |
| **Fig 4Q - Min. correlation between DA lagged relative to ACh burst in ACE2 cKO mice treated with vehicle vs 0.1mM MLN4760** |  |  |  |  |  |  |  |
|  | ACE2 cKO mice treated with vehicle |  |  | ACE2 cKO mice treated with MLN4760 |  |  |  |
|  | Mean | SEM | n | Mean | SEM | n |  |
| Min. correlation (r) | -0.2887 | 0.1160 | 7 | -0.1173 | 0.1658 | 7 | ns |
| Paired two-tailed t-test: $t(6) = 1.010, p = 0.3517$ | | | | | | | |
| **Fig 4R - Max. correlation between DA lagged relative to ACh burst in ACE2 cKO mice treated with vehicle vs 0.1mM MLN4760** |  |  |  |  |  |  |  |
|  | ACE2 cKO mice treated with vehicle |  |  | ACE2 cKO mice treated with MLN4760 |  |  |  |
|  | Mean | SEM | n | Mean | SEM | n |  |
| Max. correlation (r) | 0.5666 | 0.1108 | 7 | 0.5420 | 0.06035 | 7 | ns |
| Paired two-tailed t-test: $t(6) = 0.05787, p = 0.9557$ | | | | | | | |

**Supplementary Table S5.** (associated with Figure 5). Statistical Table.

| **Fig 5b – Animal-averaged NeuN-normalized p-rpS6 expression in dorsolateral striatal cholinergic interneurons (CINs) across four treatment groups: saline, 5 mg/kg losartan, 3 mg/kg quinpirole, and combined losartan + quinpirole.** |  |  |  |  |  |
| --- | --- | --- | --- | --- | --- |
|  |  | Mean | SEM | n |  |
| Fluorescence intensity (NeuN-normalized AU) | Saline | 1.048 | 0.03846 | 4 |  |
|  | Losartan | 0.8708 | 0.07384 | 4 |  |
|  | Quinpirole | 0.9379 | 0.04176 | 6 |  |
|  | Losartan + Quinpirole | 0.7468 | 0.04178 | 5 |  |
| Ordinary one-way ANOVA: $F(3, 15) = 6.366, ** p = 0.0054$ | | | | | |
| Post Hoc Holm-Šídák's multiple comparisons test:<br>Saline vs Losartan: $t(15) = 2.352, p = 0.1246$<br>Saline vs Quinpirole: $t(15) = 1.601, p = 0.2790$<br>Saline vs Losartan + Quinpirole: $t(15) = 4.214, ** p = 0.0045$<br>Losartan vs Quinpirole: $t(15) = 0.9758, p = 0.3446$<br>Losartan vs Losartan + Quinpirole: $t(15) = 1.735, p = 0.2790$<br>Quinpirole vs Losartan + Quinpirole: $t(15) = 2.962, * p = 0.0475$ | | | | | |

**Fig 5b – Percentage of CINs positive for AT1R-D2R dimers across genotype and PLA processing**
|  |  | Mean | SEM | n |  |
| --- | --- | --- | --- | --- | --- |
|  | D2R L/L w/ plus & minus probes | 78.3 | 8.60 | 23 |  |
|  | ChAT-Cre +/- D2R L/L w/ plus & minus probes | 14.3 | 13.2 | 7 |  |
|  | D2R L/L w/ only minus probe | 0 | 0 | 10 |  |
| | | Ordinary one-way ANOVA: $F(2, 37) = 21.54$, **** $p < 0.0001$ | | | |
| | | Post Hoc Tukey's multiple comparison test $q(df, k) = \text{value}$<br>D2R L/L vs. ChAT-Cre +/-; D2R L/L: $q(37) = 6.03$, *** $p = 0.0004$<br>D2R L/L vs. D2R L/L No Plus Probe: $q(37) = 8.407$, **** $p < 0.0001$<br>ChAT-Cre +/-; D2R L/L vs. D2R L/L No Plus Probe: $q(37) = 1.180$, $p = 0.6843$ | | | |

**Supplementary Table S6.**
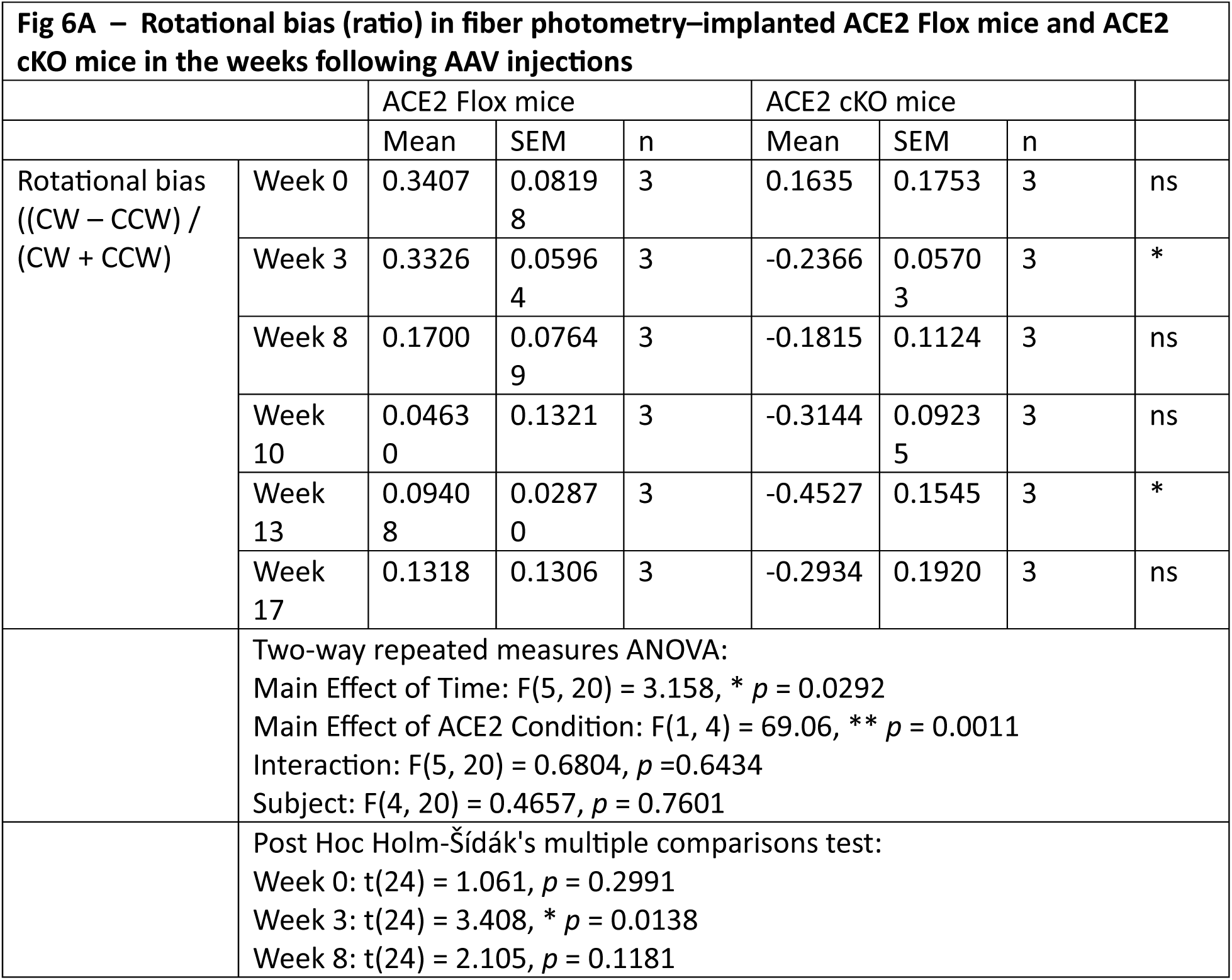
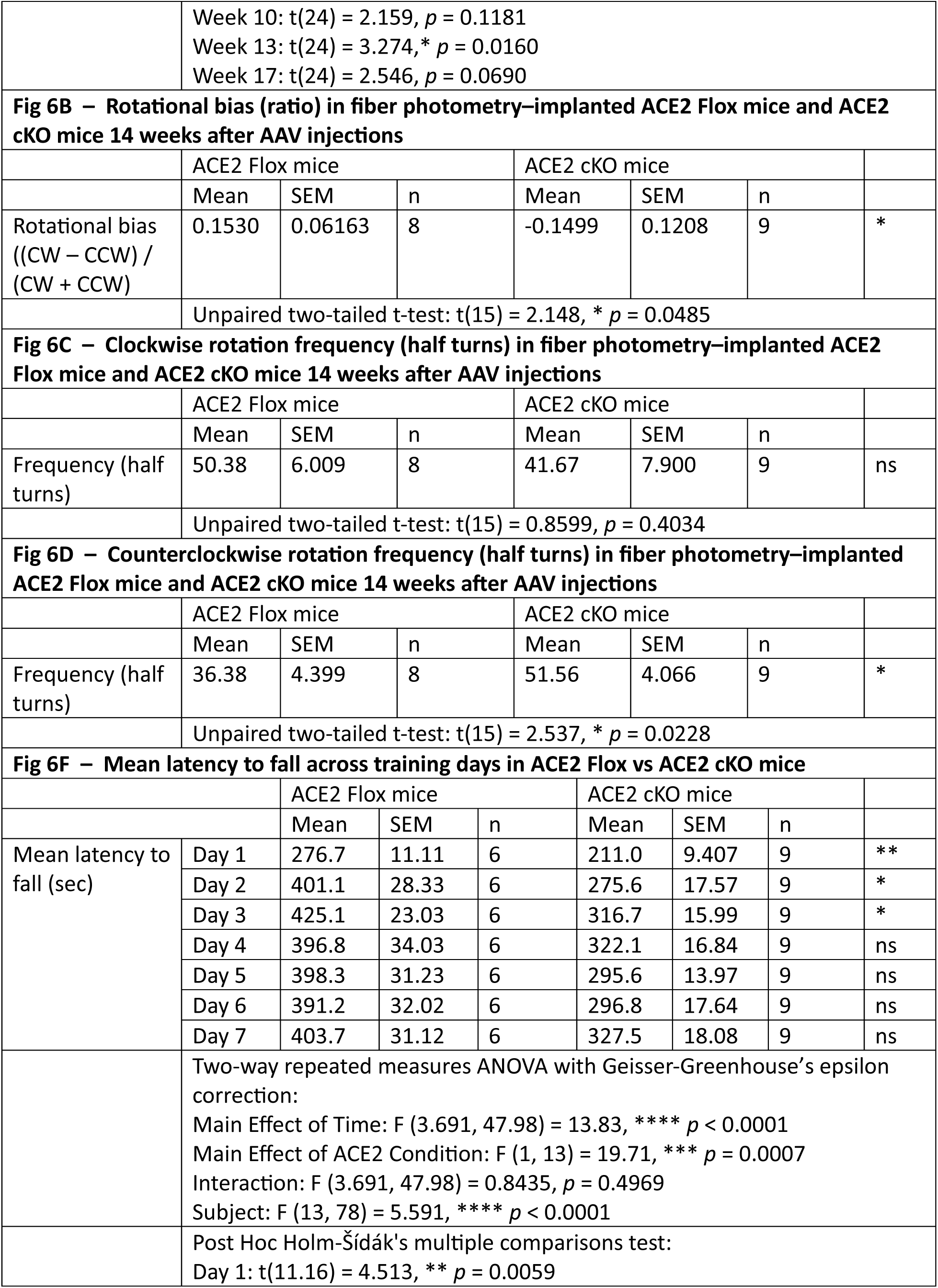
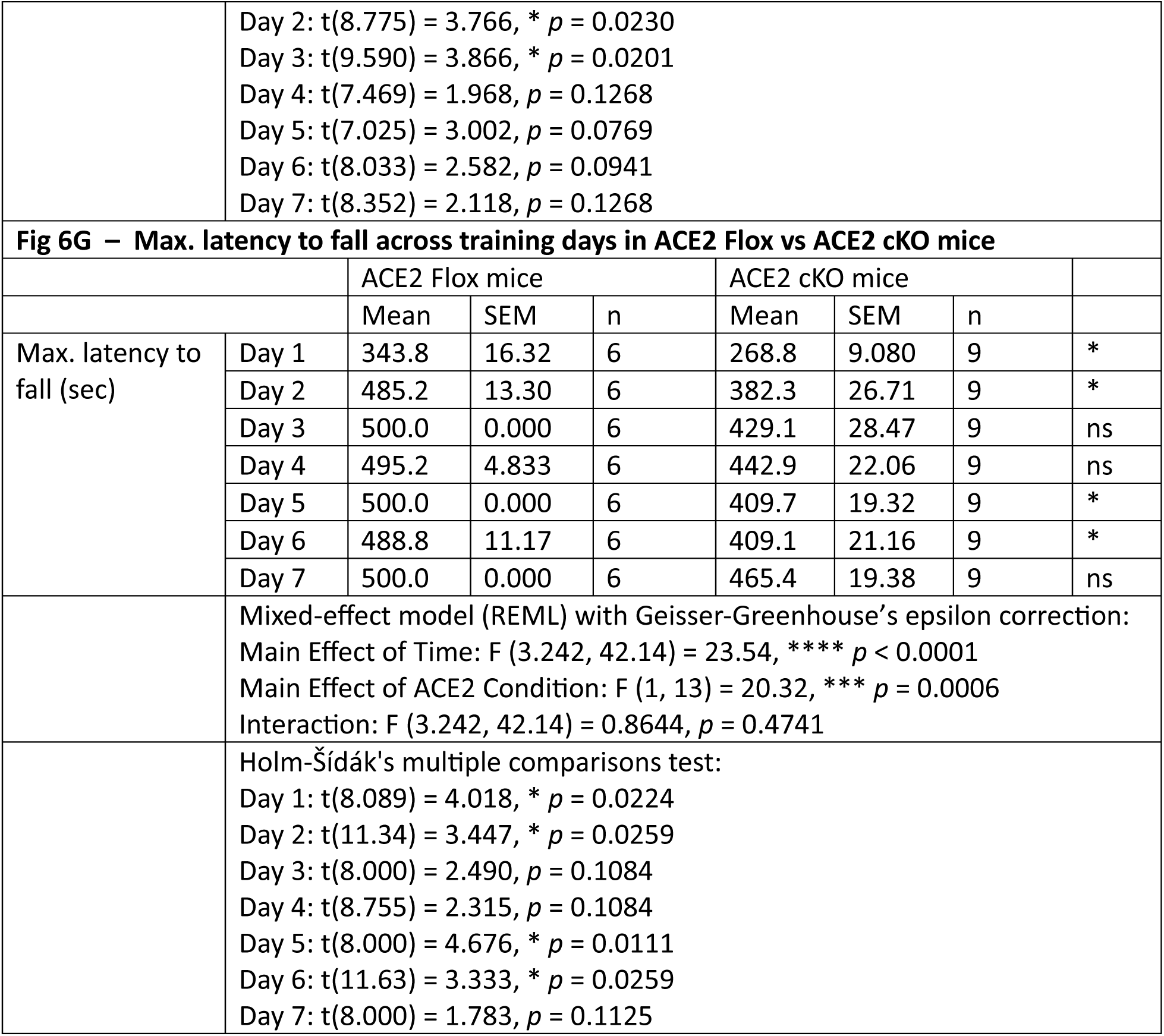
(associated with Figure 6). Statistical Table.

**Supplementary Table S7.**
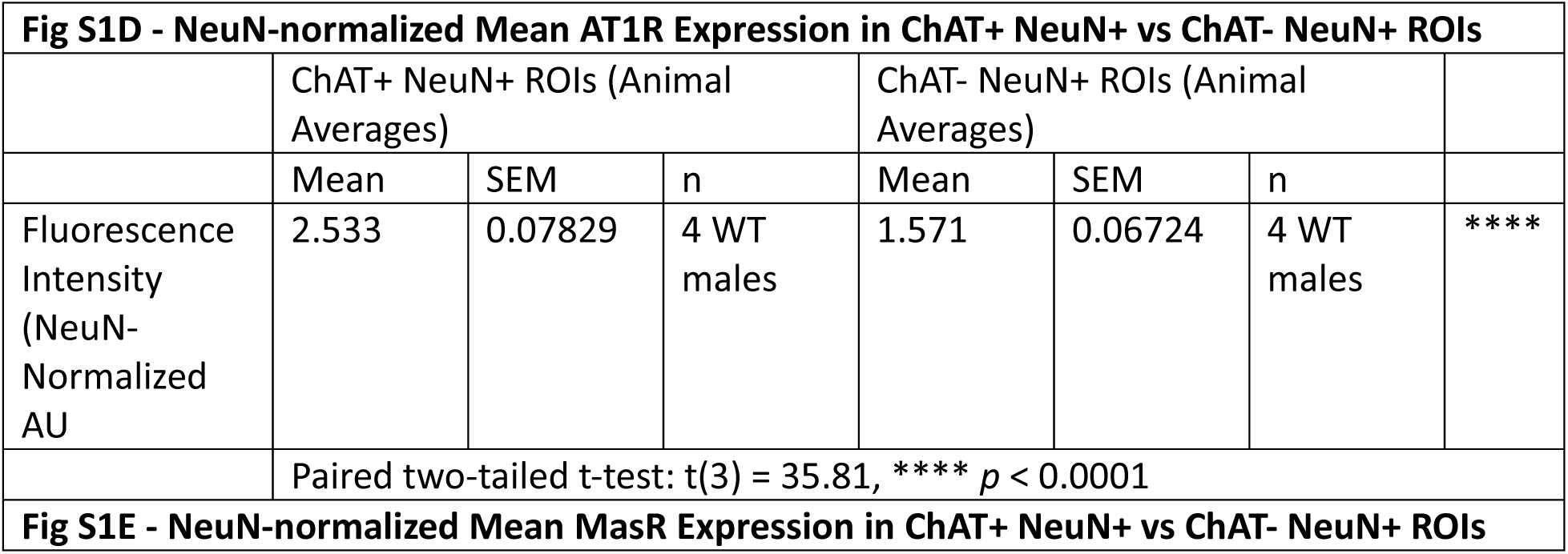
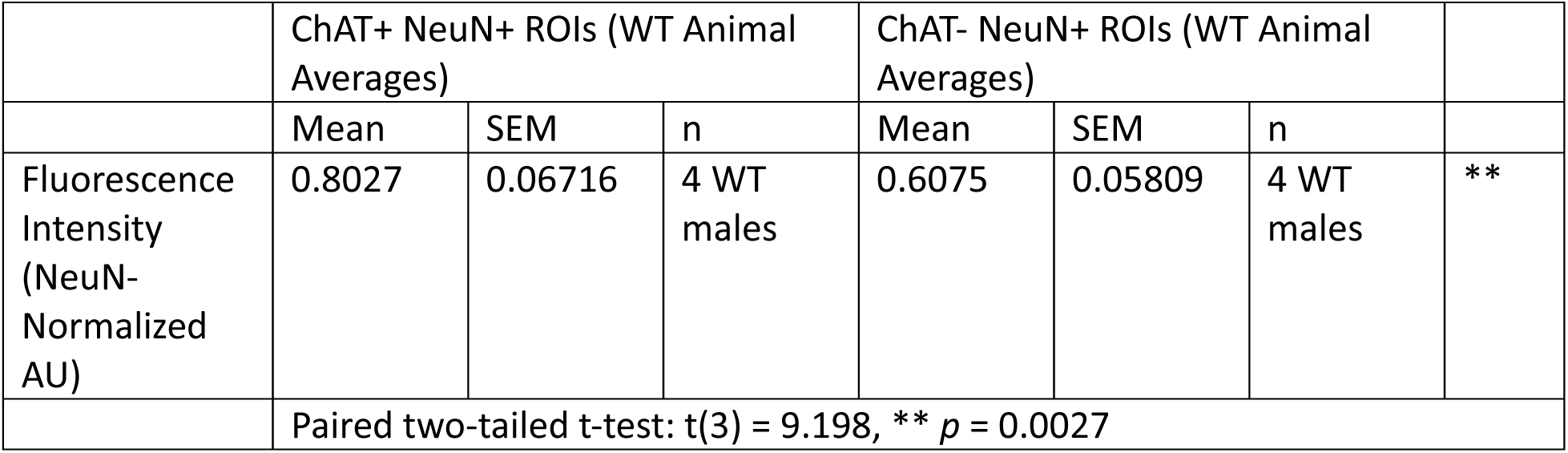
(associated with Figure S1). Statistical Table.

**Supplementary Table S8.**
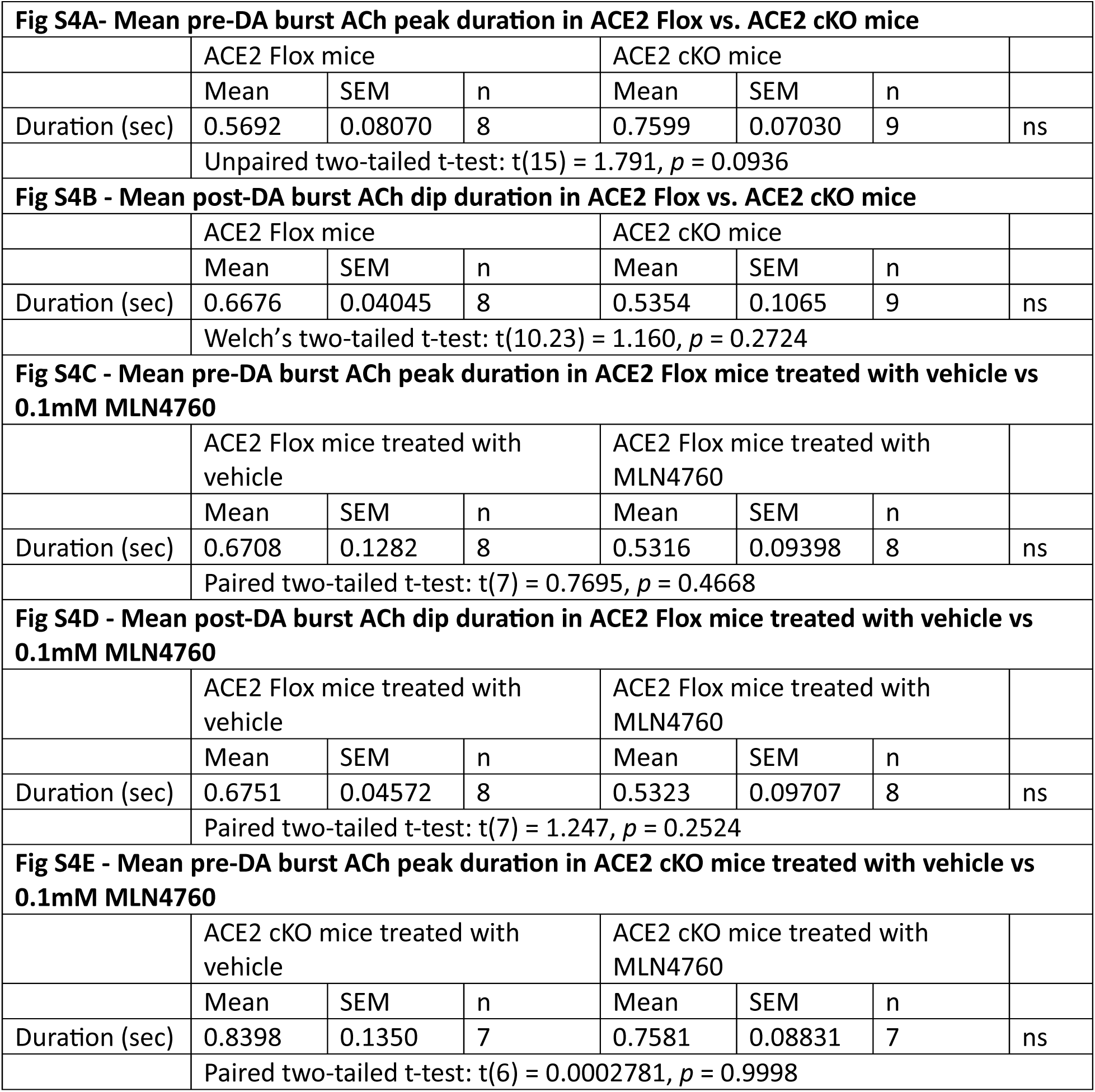
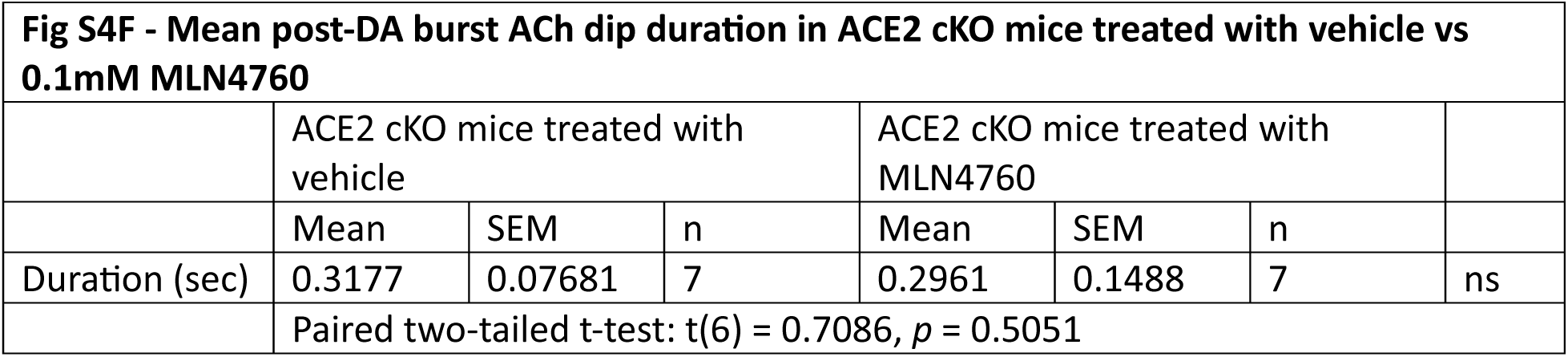
(associated with Figure S3). Statistical Table.

**Supplementary Table S9.** (associated with Figure S6). Statistical Table.

| **Fig S6A – Total distance moved during the 30-minute open field trial across four treatment groups: saline, 5 mg/kg losartan, 3 mg/kg quinpirole, and losartan + quinpirole** |  |  |  |  |  |
| --- | --- | --- | --- | --- | --- |
|  |  | Mean | SEM | n |  |
| Total distance moved (cm) | Saline | 7499 | 641.2 | 5 |  |
|  | Losartan | 8294 | 415.4 | 5 |  |
|  | Quinpirole | 1185 | 144.7 | 6 |  |
|  | Losartan + Quinpirole | 1072 | 160.1 | 6 |  |
| Welch's ANOVA tests & Brown-Forsythe ANOVA test:<br>Welch's ANOVA: $W(3, 8.670) = 104.5, **** p < 0.0001$<br>Brown-Forsythe ANOVA: $F(3, 8.099) = 102.3, **** p < 0.0001$ | | | | | |
| Post Hoc Dunnett's T3 test:<br>Saline vs Losartan: $t(6.854) = 1.040, p = 0.8630$<br>Saline vs Quinpirole: $t(4.409) = 9.606, ** p = 0.0029$<br>Saline vs Losartan + Quinpirole: $t(4.500) = 9.725, *** p = 0.0009$<br>Losartan vs Quinpirole: $t(4.971) = 16.16, **** p < 0.0001$<br>Losartan vs Losartan + Quinpirole: $t(5.185) = 16.22, **** p < 0.0001$<br>Quinpirole vs Losartan + Quinpirole: $t(9.900) = 0.5221, p = 0.9942$ | | | | | |
| **Fig S6C – Mean velocity during the 30-minute open field trial across four treatment groups: saline, 5 mg/kg losartan, 3 mg/kg quinpirole, and losartan + quinpirole** |  |  |  |  |  |
|  |  | Mean | SEM | n |  |
| Mean velocity (cm/s) | Saline | 5.833 | 0.2842 | 5 |  |
|  | Losartan | 6.521 | 0.3571 | 5 |  |
|  | Quinpirole | 0.8561 | 0.1095 | 6 |  |
|  | Losartan + Quinpirole | 0.6948 | 0.1204 | 6 |  |
| One-way ANOVA:<br>$F(3, 18) = 194.0, **** p < 0.0001$ | | | | | |
| Post Hoc Holm-Šídák's multiple comparisons test:<br>Saline vs Losartan: $t(18) = 2.071, p = 0.1032$<br>Saline vs Quinpirole: $t(18) = 15.66, **** p < 0.0001$<br>Saline vs Losartan + Quinpirole: $t(18) = 16.16, **** p < 0.0001$<br>Losartan vs Quinpirole: $t(18) = 17.82, **** p < 0.0001$<br>Losartan vs Losartan + Quinpirole: $t(18) = 18.33, **** p < 0.0001$<br>Quinpirole vs Losartan + Quinpirole: $t(18) = 0.5322, p = 0.6011$ | | | | | |

**Supplementary Table S10.** (associated with Figure S7). Statistical Table.

| **Fig S7B – Percentage of ChAT- NeuN+ and ChAT+ NeuN+ ROIs positive for AT1R-D2R dimers across genotype and PLA processing** |  |  |  |  |  |  |  |  |
| --- | --- | --- | --- | --- | --- | --- | --- | --- |
|  |  | ChAT- NeuN+ |  |  | ChAT- NeuN+ |  |  |  |
|  |  | Mean | SEM | n | Mean | SEM | n |  |
| Positive ROIs (ratio) | D2R L/L | 0.686 | 0.021 | 497 | 0.783 | 0.086 | 23 |  |
|  | ChAT-Cre +/- D2R L/L | 0.541 | 0.033 | 229 | 0.143 | 0.132 | 7 |  |
|  | D2R L/L No Plus Probe | 0.259 | 0.028 | 251 | 0.000 | 0.000 | 10 |  |
| | | Two-way repeated measures ANOVA:<br>Main Effect of Cell Type: $F(1, 1011) = 4.874$, * $p = 0.0275$<br>Main Effect of Genotype: $F(2, 1011) = 24.82$, **** $p < 0.0001$<br>Interaction: $F(2, 1011) = 3.882$, * $p = 0.0209$ | | | | | | |
| | | Post Hoc Tukey's multiple comparisons test:<br>ChAT- NeuN+:<br>D2R L/L vs. ChAT-Cre +/-; D2R L/L: $q(1011) = 5.513$, *** $p = 0.0003$<br>D2R L/L vs. D2R L/L – No Plus Probe: $q(1011) = 16.75$, **** $p < 0.0001$<br>ChAT-Cre +/-; D2R L/L vs. D2R L/L – No Plus Probe: $q(1011) = 9.372$, **** $p < 0.0001$<br>ChAT+ NeuN+:<br>D2R L/L vs. ChAT-Cre +/-; D2R L/L: $q(1011) = 4.503$, ** $p = 0.0043$<br>D2R L/L vs. D2R L/L – No Plus Probe: $q(1011) = 6.278$, **** $p < 0.0001$<br>ChAT-Cre +/-; D2R L/L vs. D2R L/L – No Plus Probe: $q(1011) = 0.8812$, ns $p = 0.8075$<br>D2R L/L:<br>ChAT- NeuN+ vs. ChAT+ NeuN-: $q(1011) = 1.381$, ns $p = 0.3290$<br>ChAT-Cre +/-; D2R L/L:<br>ChAT- NeuN+ vs. ChAT+ NeuN-: $q(1011) = 3.150$, * $p = 0.0261$<br>D2R L/L – No Plus Probe:<br>ChAT- NeuN+ vs. ChAT+ NeuN-: $q(1011) = 2.439$, ns $p = 0.0849$ | | | | | | |

**Supplementary Table S11.**
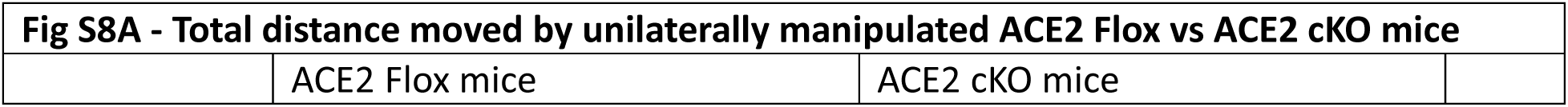
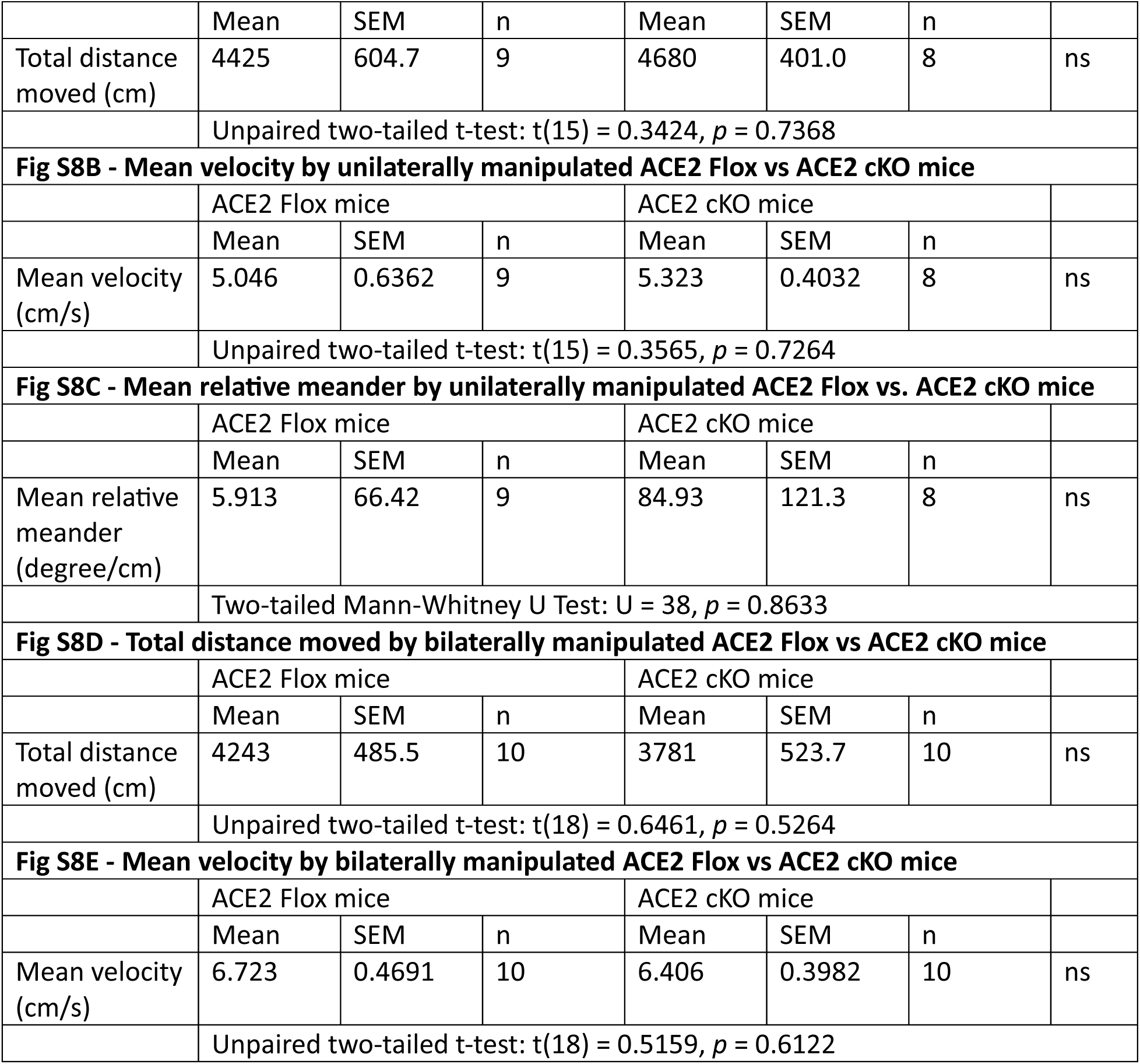
(associated with Figure S8). Statistical Table.

## Methods

Resource availability

## Lead contact

Further information and requests for resources and reagents should be directed to and will be fulfilled by the lead contact, Andreas Kottmann.

## Materials availability

The material reported in this article will be shared by the lead contact upon request.

## Data and code availability

All data supporting the findings of this study are available within the paper and its supplementary information.

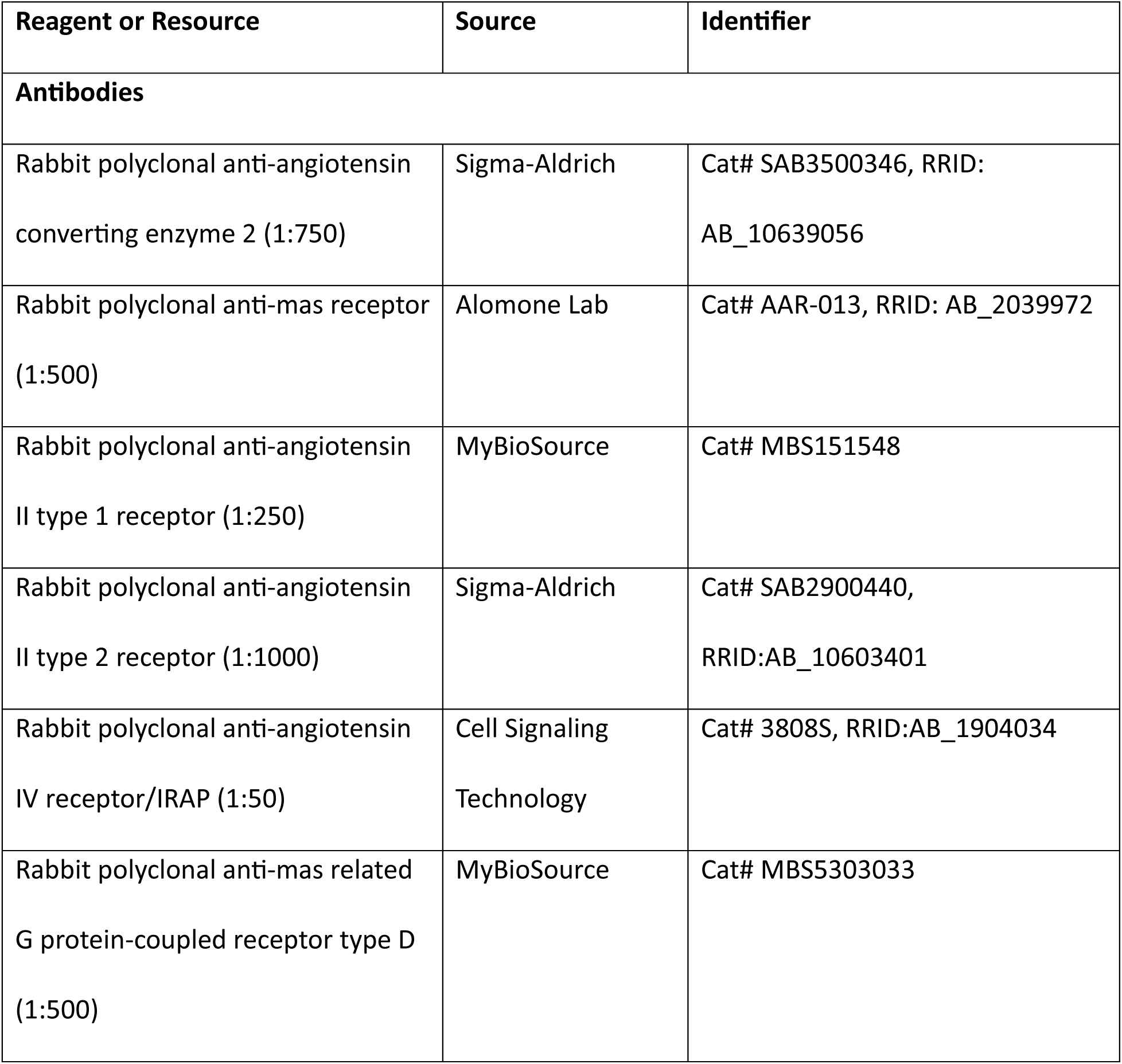
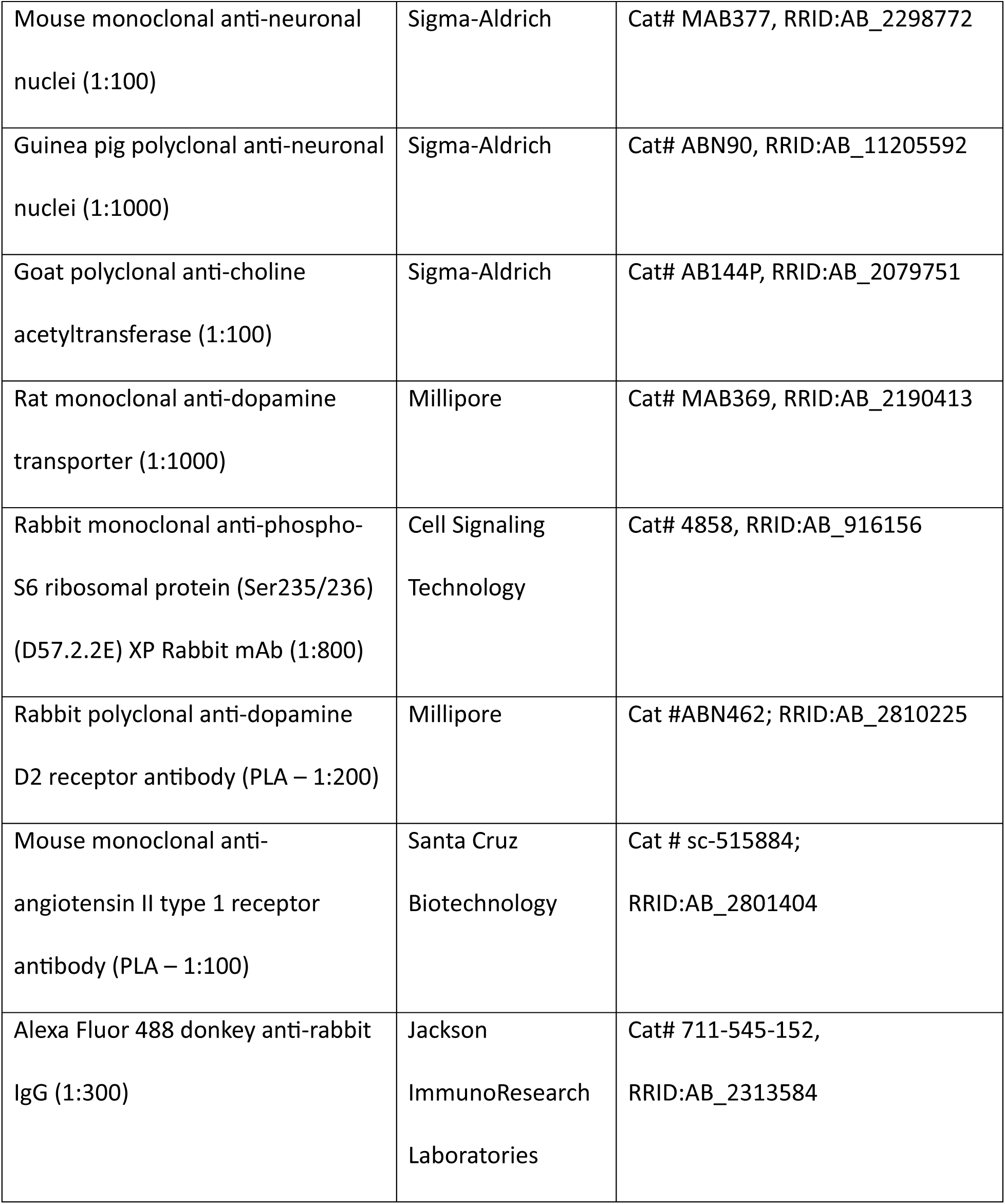
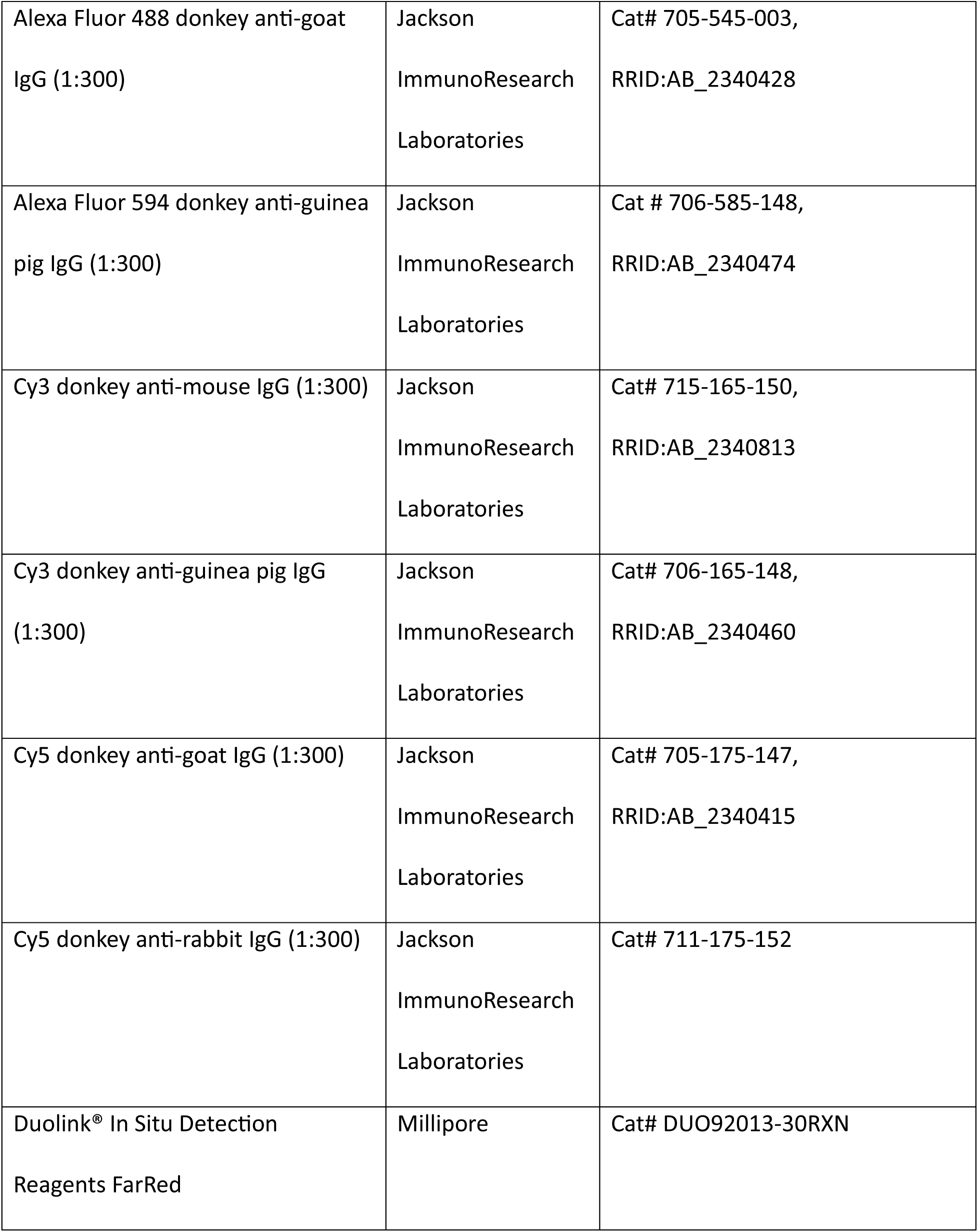
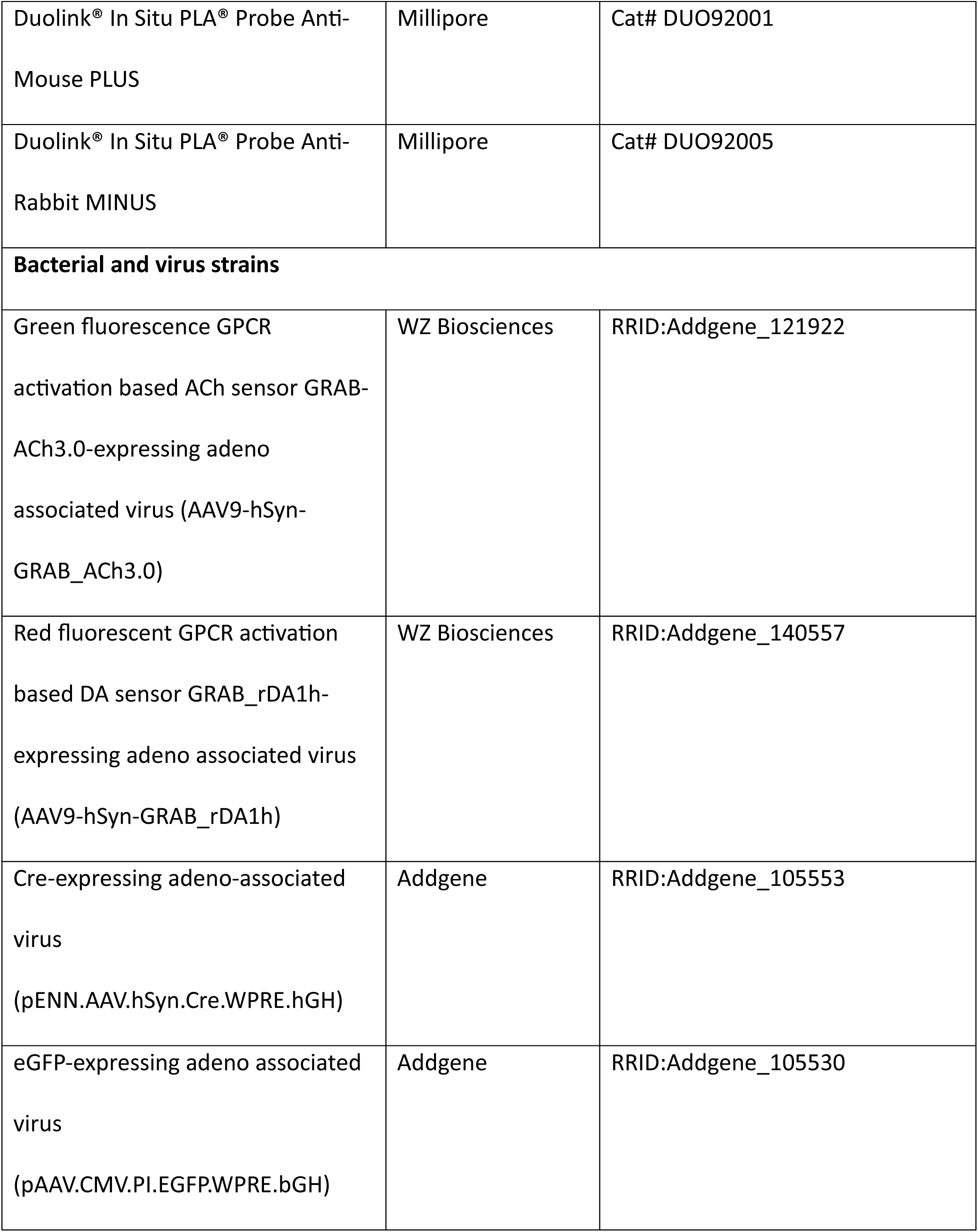
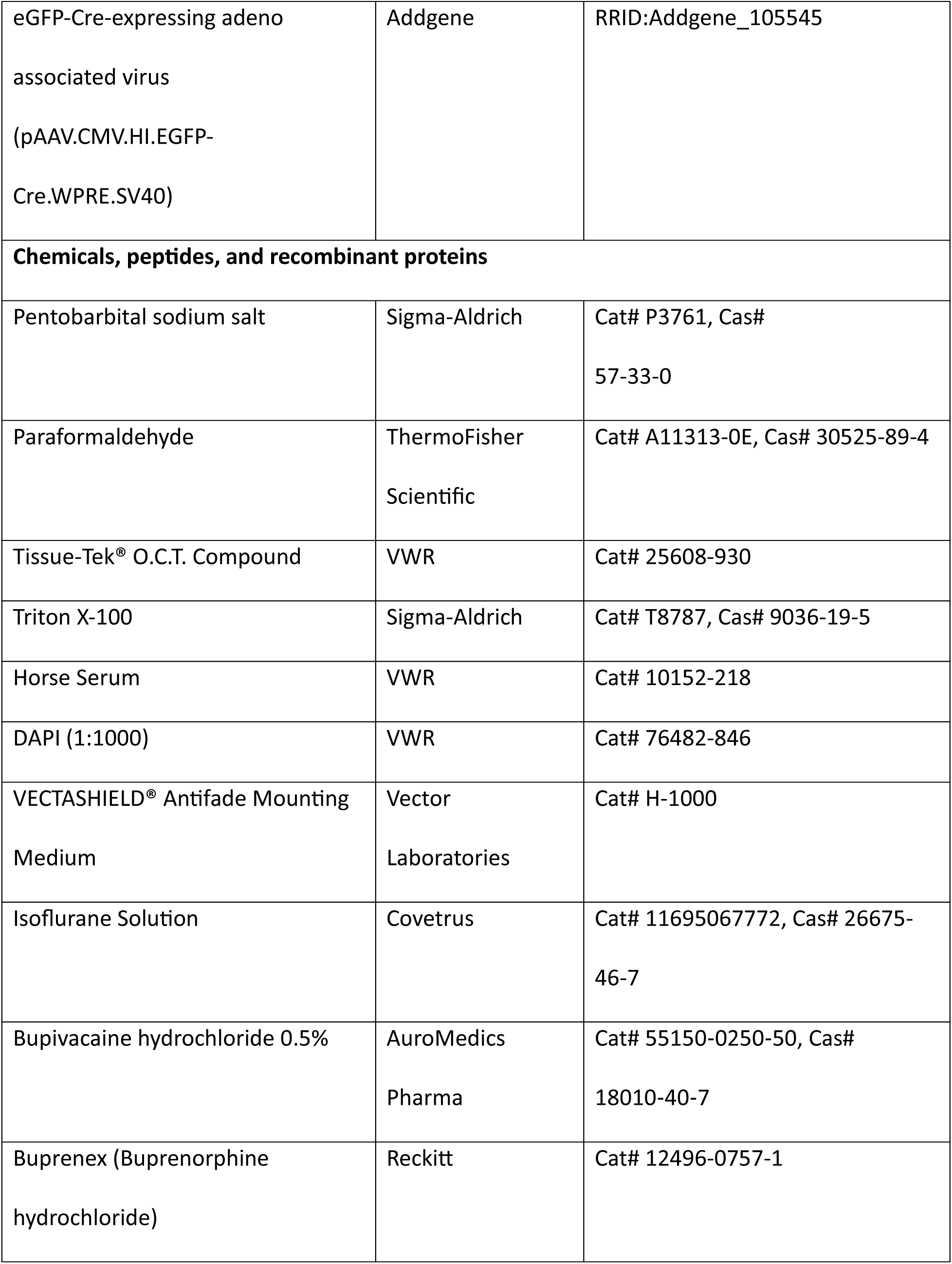
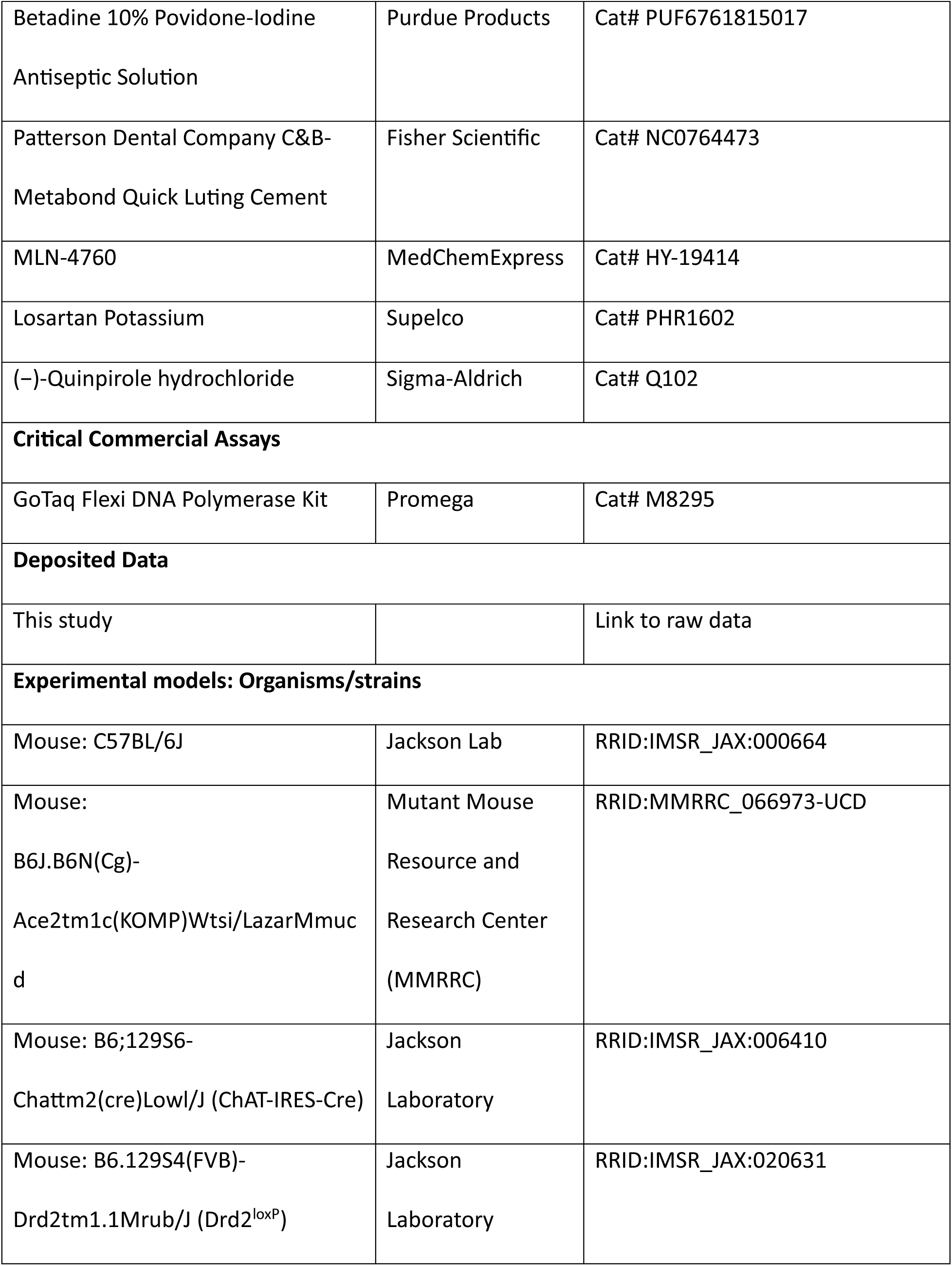
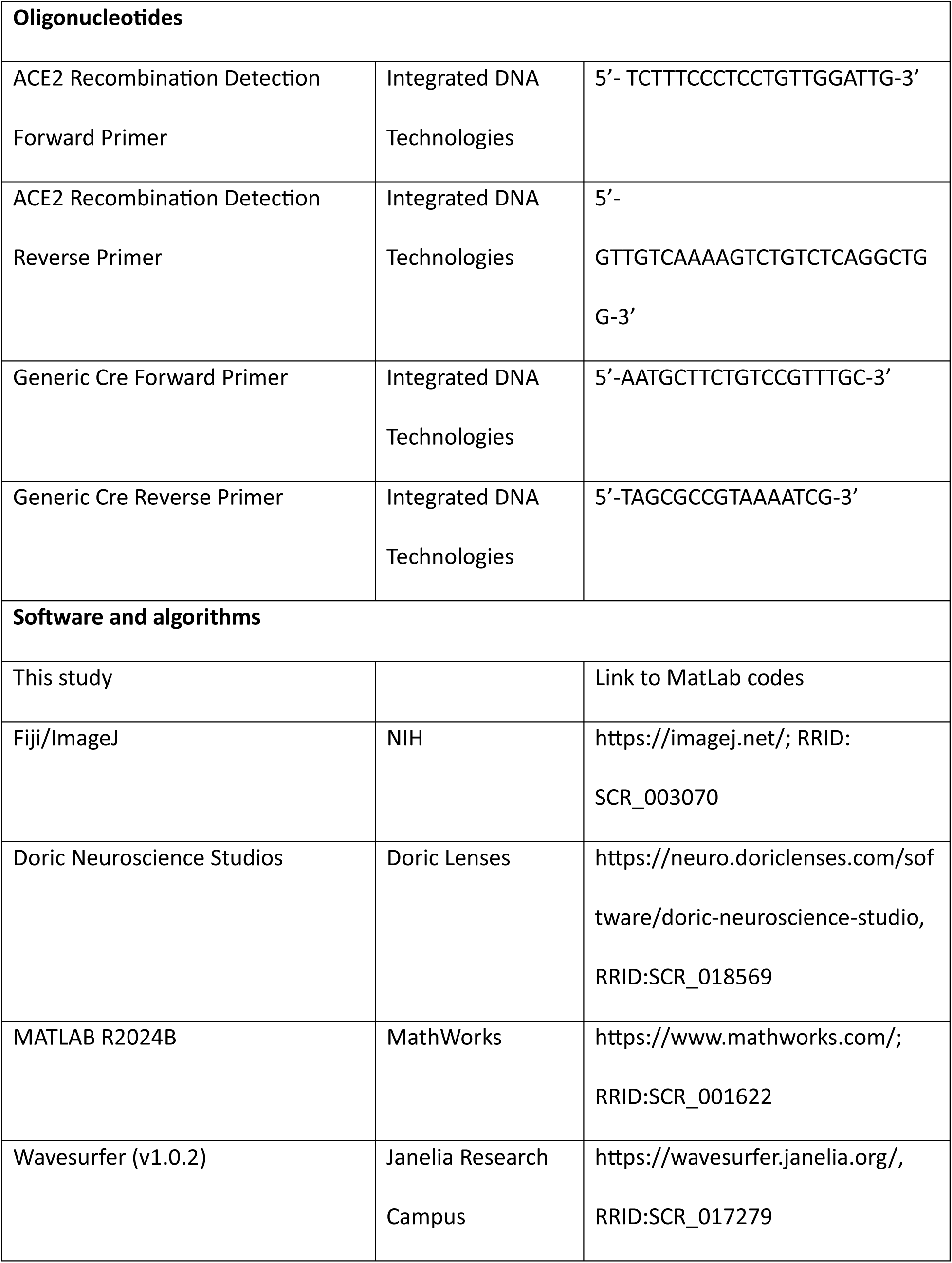
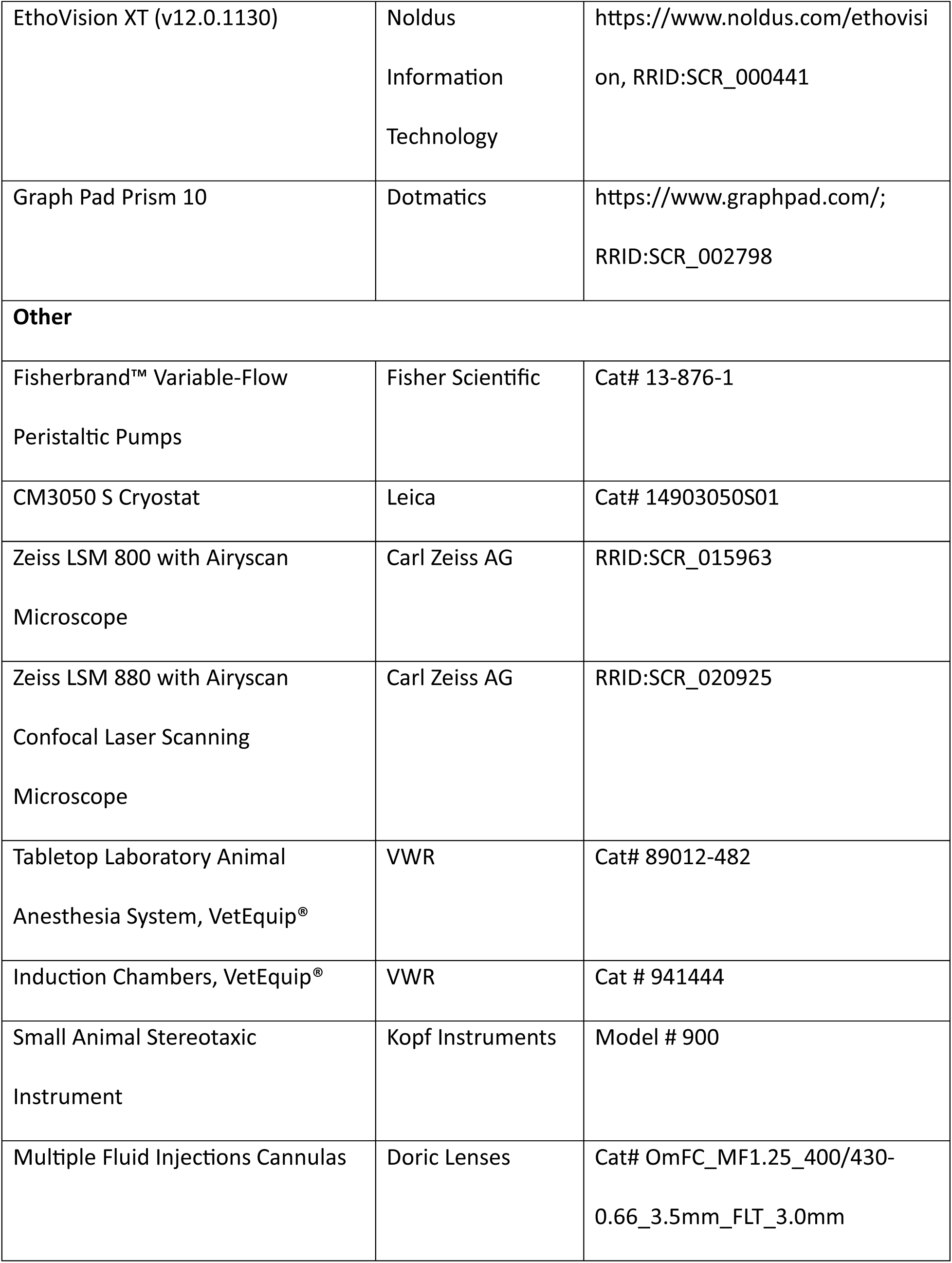
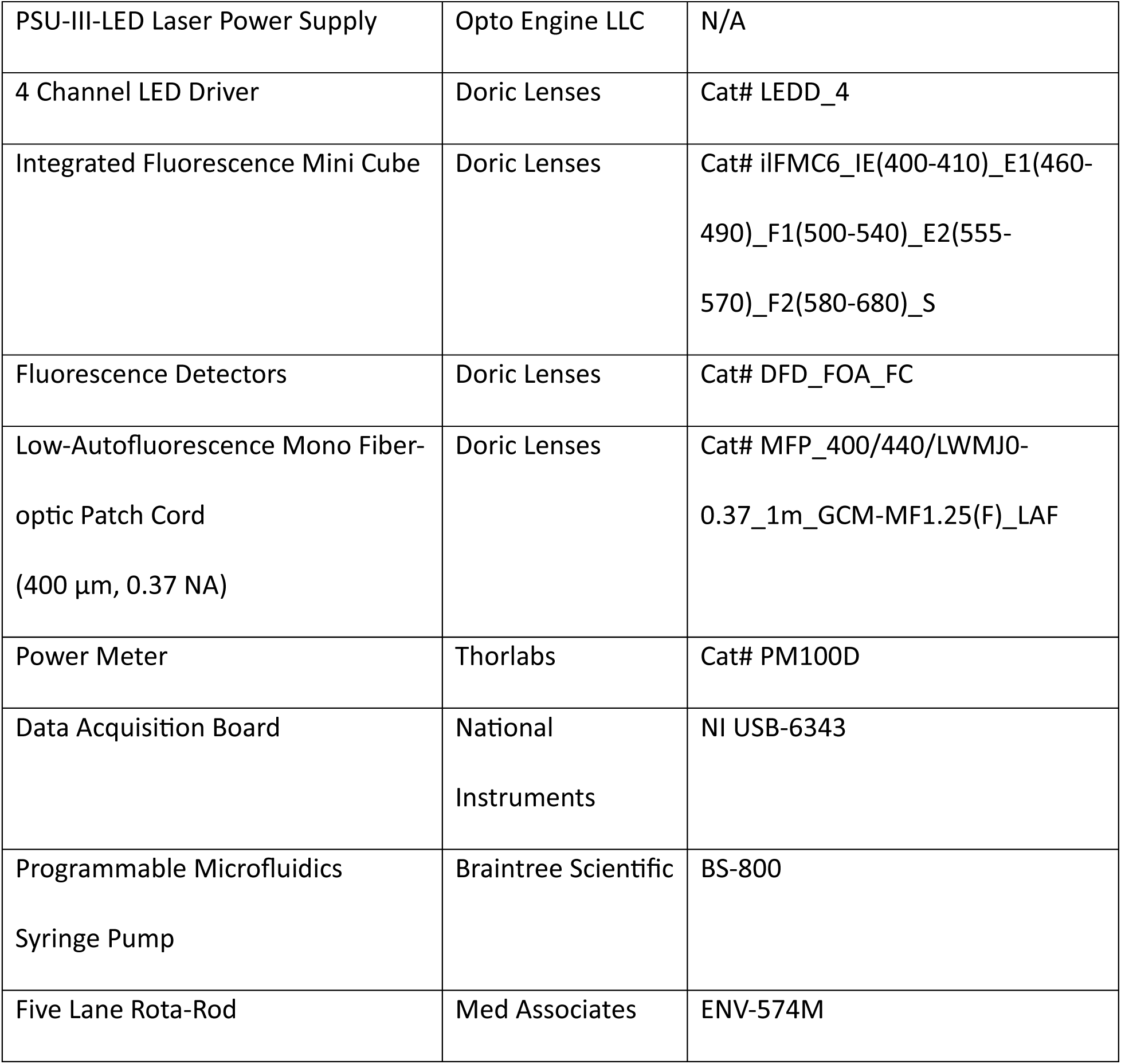
Key resource table.

## Experimental model and subject details

Male wildtype C57BL/6J mice were purchased from the Jackson Laboratory. ACE2^flox^ mice (stock #066973-UCD; hereafter referred to as “ACE2 Flox“) were obtained from the Mutant Mouse Resource and Research (MMRRC) at the University of California, Davis. Genotyping of transgenic animals was performed using the PCR-based protocol provided by MMRRC. Littermates of the same sex were randomly assigned to experimental groups.

Mice were group housed (2-5 per cage) in ventilated cages maintained at 23°C under a 12-hour light/dark cycle, with ad libitum access to food and water. Cage bedding was changed weekly. All procedures were conducted in accordance with protocols approved by the Institutional Animal Care and Use Committee (IACUC) of the City College of New York (CCNY).

## Method details

### Experimental Design

The study was designed to assess the role of ACE2 enzymatic activity in striatal cholinergic interneurons. ACE2 Flox mice received AAV-Cre or vehicle injections unilaterally into the DLS, and their behavioral and neuromodulatory profiles were assessed using fiber photometry, open field testing, and rotarod learning tasks.

### Immunohistochemistry

Mice were deeply anesthetized with intraperitoneal pentobarbital (15 mg/kg) and perfused transcardially with 0.9% saline followed by 4% paraformaldehyde (PFA) in 0.1 M phosphate-buffered saline (PBS). Brains were post-fixed overnight at 4°C in 4% PFA, then cryoprotected in 30% sucrose in PBS until they sank (∼48 hours). Cryoprotected brains were embedded in optimal cutting temperature (OCT) compound, frozen on dry ice, and stored at –80°C. Brains were cryosectioned coronally at 30 µm using a cryostat at –20°C and stored as free-floating sections in PBS with 0.1% sodium azide until staining.

On day 1 of staining, sections were washed 3 × 15 minutes in PBS to remove excess OCT, permeabilized with 0.3% Triton X-100 (v/v) in PBS for 15 minutes and blocked overnight at 4°C in PBS containing 10% horse serum and 0.3% Triton X-100.

On day 2, sections were incubated with primary antibodies diluted in blocking buffer at 4°C overnight. On day 3, sections were washed 3 ×15 minutes in PBS containing 2% horse serum and 0.03% Triton X-100, then incubated with fluorophore-conjugated secondary antibodies and DAPI in PBS for two hours at room temperature. Sections were washed again 3 ×15 minutes in PBS containing 2% horse serum and 0.03% Triton X-100 to remove excess secondary antibodies and mounted using a VectaShield Mounting Medium.

Fluorescent imaging was performed using a Zeiss LSM 800 or 880 confocal microscope. Quantitative image analysis was conducted in ImageJ.

### Proximity Ligation Assay

Free-floating sections were washed in PBS for 5 minutes at room temperature to remove superficial azide, followed by a 10-minute wash in 50:50 PBS:TBS to gradually shift buffer composition. Sections were then washed four times in TBS (0.1 M Tris, pH 7.3, 0.9% NaCl) for 15 minutes each at room temperature, permeabilized in 0.1% Triton X-100 in TBS for 15 minutes, and blocked in TBS containing 10% horse serum, 3% BSA, and 0.1% Triton X-100 first for 30–60 minutes at room temperature and then overnight at 4°C.

On day 2, sections were washed three times in TBS for 5 minutes each at room temperature and incubated overnight at 4°C with primary antibodies diluted in TBS containing 10% horse serum, 1% BSA, and 0.1% Triton X-100.

On day 3, sections were washed four times in TBS containing 0.05% Tween-20 for 30 minutes each at room temperature, then incubated for 2 hours at 37°C with Duolink PLA probes (Plus and Minus) diluted in TBS with 0.05% Tween-20 and 1% BSA. Control sections were processed with only Plus or Minus probe. Following incubation, sections were washed three times in TBS containing 0.05% Tween-20 and 300 mM NaCl for 15 minutes each at room temperature, then washed once in plain TBS for 10 minutes. Ligation was performed for 30 minutes at 37°C, followed by three additional washes in TBS with 0.05% Tween-20 and 300 mM NaCl. Amplification was then carried out at 37°C for 90 minutes, protected from light. Sections were washed twice in TBS with 0.05% Tween-20 and 300 mM NaCl for 15 minutes each, followed by two washes in PBS for 15 minutes each at room temperature.

For subsequent immunostaining, sections were blocked in PBS containing 10% horse serum and 0.1% Triton X-100 for 30–60 minutes at room temperature, then incubated overnight at 4°C with goat anti-ChAT and guinea pig anti-NeuN primary antibodies. On day 4, sections were washed four times in PBS containing 2% horse serum and 0.03% Triton X-100 for 15 minutes each, then incubated for 2 hours at room temperature with fluorophore-conjugated secondary antibodies (AF488 anti-goat, Cy3 anti-guinea pig) and DAPI. After three final washes in PBS containing 2% horse serum and 0.03% Triton X-100 for 15 minutes each, sections were mounted in VectaShield and imaged using a confocal microscope.

### Stereotaxic Surgery and Viral Injections

Mice were anesthetized with 2% isoflurane in a VetEquip induction chamber and placed in a Kopf stereotaxic frame. The scalp was shaved and sterilized with alternating betadine and sterile water. Ophthalmic ointment (Puralube) was applied to protect the eyes. Local anesthesia (100 µL of 0.5% bupivacaine) was injected subcutaneously beneath the scalp. An anterior-posterior incision was made to expose the skull, and connective tissue was cleared with 3% hydrogen peroxide.

To improve adhesive bonding, the skull surface was scored using a scalpel by making a cross-hatching pattern. A small hole was drilled at lambda to attach a headbar with a stainless-steel screw. A craniotomy was performed using a dental drill above the dorsolateral striatum (DLS), and a fiberoptic multiple fluid injection cannula (Doric Lenses; OmFC_MF1.25_400/430-0.66_3.5mm_FLT_3.0mm) was stereotaxically implanted at the following coordinates from bregma: AP +0.7 mm, ML +2.2 mm, DV –3.0 mm. The implant was secured using C&B Metabond Quick Adhesive Cement.

Post-operatively, mice received intraperitoneal injections of 400 µL 5% sucrose in 0.9% saline and a subcutaneous injection of 0.05 mg/kg buprenorphine. Mice recovered for three weeks prior to virus injection.

AAV stocks were used as received from Addgene/WZ Biosciences (titers ranging from 10¹²–10¹³ GC/mL). A 1:1 mixture of AAV-hSyn-GRAB_ACh3.0 and AAV9-hSyn-GRAB_rDA1h (rDA2.5h) was delivered through the cannula (0.5 µL total volume) at a rate of 5 µL/min using a programmable syringe pump. Mice were given an additional three weeks for viral expression prior to experimentation and were habituated to head fixation atop a running wheel in a dark, sound attenuated cabinet. ACE2 genetic ablation in the DLS was achieved by injecting 1 µL of AAV-hSyn-Cre-WPRE-hGH through the cannula of experimental animals. ACE2 Flox control mice received 1 µL of cortex buffer as a vehicle control.

### Fiber Photometry Recording

Recordings were performed using a Doric Lenses basic fiber photometry system composed of a 4-channel LED driver (LEDD_4), Integrated Fluorescence Mini Cube (ilFMC6_IE), dual-channel fluorescence detectors (DFD_FOA_FC), and a 400 µm 0.37 NA patch cable (MFP_400/440/LWMJ0-0.37_1m_GCM-MF1.25(F)_LAF).

Signal acquisition and synchronization were performed using WaveSurfer (v1.0.2; Janelia Research Campus), interfaced with a National Instruments USB-6343 data acquisition board. LED output was controlled through Doric Neuroscience Studio.

Prior to recording, the patch cable was photobleached using ≥25 mW laser power for ≥30 minutes to reduce autofluorescence. LED output for 470 nm and 568 nm channels was then calibrated to ∼30 µW at the tip of the patch cord using a PM100D power meter (Thorlabs). Excitation light sources included 405 nm (isosbestic control), 470 nm (GRAB-ACh3.0 excitation), and 568 nm (GRAB-rDA1h excitation). LEDs were pulsed in 10 ms intervals in a cyclic sequence (405 → 470 → 568), and emitted signals were sampled at 1000 Hz.

Each session included a 15-minute baseline recording, followed by a 45-minute recording period. For sessions involving drug delivery, pharmacological agents were injected intrastriatally via the implanted cannula using a Braintree Scientific programmable microfluidics syringe pump (Model BS-800) at a volume of 0.5 µL, delivered at a rate of 5 µL/min.

### Fiber Photometry Analysis

Raw data were processed using custom MATLAB scripts. Signals were extracted from WaveSurfer-exported.h5 files, demultiplexed, and concatenated into continuous time series. A precise time array was reconstructed based on the acquisition frequency.

Photobleaching was corrected using an exponential decay model. Each channel (isosbestic, ACh3.0, rDA1h) was filtered using a 20 Hz low-pass infinite impulse response filter, and signals were downsampled using a 100-point moving mean.

Normalized ΔF/F_0_ traces were calculated by performing a linear fit between the fluorescent signal and its isosbestic control to define the F_0_. ΔF/F_0_ values were then z-scored to normalize across sessions.

Burst event detection was performed by identifying periods when the normalized signal exceeded 3 median absolute deviations (MADs) above a filtered local baseline. The baseline was defined as a 150-second moving median, excluding high-transient data points (>2 MADs) from the calculation.

### Open Field Test

Mice were placed individually in a rat open field arena (43 × 43 cm) located in a quiet, dimly lit behavioral testing room. Each session lasted 15–30 minutes, during which animals were allowed to freely explore the arena. Behavior was recorded using infrared (IR) video cameras positioned either above or below the arena, depending on concurrent experimental setup. The arena was cleaned with 70% ethanol between sessions.

Videos were acquired and analyzed using EthoVision XT (v12.0.1130; Noldus Information Technology). Locomotor activity was quantified using center point tracking. Measured parameters included total distance moved, mean velocity, turn frequency, and relative meander.

### Accelerating Rotarod

Mice were habituated to the rotarod apparatus (Med Associates, ENV-574M) on day 0, which consisted of a 300-second period atop the stationary rod followed by 300 seconds in the holding basket beneath the apparatus. Beginning on day 1, mice were trained using ten trials per day for seven consecutive days.

Each trial began with the rotarod set to 4 rpm and accelerating linearly to 40 rpm over the first 300 seconds. This was followed by a constant 40 rpm phase lasting up to 500 additional seconds. If a mouse completed one full passive rotation, it was manually removed from the apparatus and recorded as having fallen. A minimum 300-second rest period was provided between trials to prevent fatigue.

Animals that failed to reach a latency to fall of 300 seconds or more during the training period were classified as non-learners and excluded from analysis. Rotarod data was statistically analyzed using GraphPad Prism.

### Quantification and Statistical Analysis

Raw fiber photometry data were stored as WaveSurfer-exported.h5 files and analyzed using custom MATLAB scripts (see Fiber Photometry Analysis). All other raw data, including fluorescence intensities extracted from regions of interest (ROIs) in confocal images, open field locomotor metrics, and latency to fall in the accelerating rotarod assay, were exported to Microsoft Excel and analyzed using GraphPad Prism v10 (Dotmatics).

Statistical tests were selected based on the distribution and variance of each dataset. Normality was assessed using Shapiro-Wilk, D’Agostino-Pearson omnibus, and Anderson-Darling tests, and homoscedasticity was evaluated using Brown-Forsythe or Bartlett’s tests where appropriate. For normally distributed data with equal variance, unpaired or paired two-tailed t-tests, one-way ANOVAs, or two-way repeated-measures ANOVAs were used. Welch’s ANOVA or mixed-effects models were applied when assumptions were violated. Post hoc comparisons were corrected using Holm–Šídák’s Multiple comparisons test unless otherwise noted. Significance was defined as p < 0.05.

Statistical tests for each figure panel are detailed in the figure legends and summarized in the file ‘Supplemental Statistical Analysis for Figures’.

## Acknowledgment

This work was supported by a City University of New York (CUNY) Interdisciplinary Research Grant (IRG, PI A.H.K) and the Louise Lennihan Arts and Science Grant to A.R.W.. A.H.K acknowledges support through NIH U54MD017979 (PI Maria Lima) and A.R.W. acknowledges the invaluable support received from the biology graduate program of the Graduate Center of the City University of New York.. We thank Sonia Bernal for her essential contributions to laboratory maintenance and organization. We also thank Michael Holmes, Layla Morgan, and Lillian Lee for their help with cytohistochemical data collection. We thank Dr. Santiago Uribe-Cano for his support and comments on early versions of this manuscript.

## Notes

### Competing Interest Statement

The authors have declared no competing interest.

